# Developmental variation in dopamine neurobiology, neurocognitive functioning, and impulsivity shape substance use trajectories in youth

**DOI:** 10.1101/2025.10.24.684358

**Authors:** Ashley C. Parr, Amar Ojha, Daniel J. Petrie, Finnegan J. Calabro, Brenden Tervo-Clemmens, Will Foran, Douglas Fitzgerald, Susan F. Tapert, Kate Nooner, Wesley Thompson, David Goldston, Duncan Clark, Beatriz Luna

## Abstract

Theoretical neurodevelopmental models implicate increases in dopamine (DA) function and limitations in neurocognitive control in risk-taking behavior, including substance use, during the transition from adolescence to adulthood. However, developmental relationships between DA, neurocognitive control, and the emergence of substance use are poorly understood. Here, we tested the role of basal ganglia tissue iron, reflecting DA neurophysiology, as well as impulsivity and inhibitory control in longitudinal developmental trajectories of substance use. We leveraged the National Consortium on Alcohol and NeuroDevelopment in Adolescence and Adulthood (NCANDA-A) cohort, a large, multisite longitudinal neuroimaging study of 807 participants (baseline ages 12 – 22 years old, 50% female, 1 – 9 annual visits per participant, 6164 sessions total). Substance use, inhibitory control, and tissue iron increased non-linearly during adolescence into young adulthood, concurrent with decreases in impulsivity. Non-linear Growth Mixture Models identified four common trajectories of substance use: *low* (no- or low levels of use across visits; 30% of participants), *youth peak* (peak use in adolescence/young adulthood followed by declines; 26%), *adolescent increasing* (early, steep linear increases in use from adolescence into adulthood; 17%), and *adult increasing* (low use in adolescence, followed by late linear increases into adulthood; 26%). We show that increased substance use was associated with a phenotype of *high* impulsivity, *low* inhibitory control, and *low* basal ganglia tissue iron, particularly in early adolescence in individuals who displayed youth peak patterns in substance use trajectories. These findings highlight that early developmental differences in DA-related neurobiology and associated impulsivity and cognitive control shape distinct trajectories of adolescent substance use, underscoring adolescence as a critical window for the early identification and implementation of neurodevelopmentally sensitive interventions for substance use disorders.

## Introduction

Illicit substance use increases dramatically during the teenage years. In 2024, only 15% of 8^th^ graders in the United States reported using illicit substances^1^, compared to approximately 37% of 12^th^ graders^1^. Substance use is a leading contributor to preventable morbidity and mortality in youth, driving a range of behavioral and adverse health outcomes including suicidality and self-harm^2,3^, interpersonal violence^2^, and accidental death (i.e., road injuries^2^ and drug-related toxicity^4,5^). As such, youth substance use poses a critical public health challenge, necessitating the implementation of evidence-based policies and developmentally tailored interventions. Neurodevelopmental theories conceptualize youth substance use in the context of dual risk models, including the Driven Dual Systems Model^6,7^, which proposes that ongoing neurobiological changes within prefrontal cortex (PFC) cognitive control systems and subcortical dopaminergic (DA) reward systems contribute to normative increases in adolescent sensation seeking, risk-taking, and reward-driven behavior, potentially creating a window of vulnerability heightened levels of substance use.

Specifically, ‘dual-risk’ theories of adolescent substance use propose that *hyper*-active reward reactivity in striatal regions coupled with *hypo*-function of cognitive control systems confers risk for problematic use^8–11^, potentially via an increase of an already-enhanced reward drive^12^ coupled with ongoing maturation of prefrontal executive control systems in adolescence^8^. This model has gained empirical support, with striatal reward *hyper-*activation identified as a consistent and reliable functional brain marker of adolescent substance use vulnerability^8,13^, as well as studies showing that cognitive performance and activation across cortical control networks are associated with substance use risk-factors (i.e., impulsivity and externalizing characteristics^14,15^) and substance use onset^16^, respectively. At the systems-level, normative adolescent increases in risk-taking have been associated with enhanced functional connectivity within mesocorticolimbic circuits, including between the ventromedial PFC and the ventral striatum (VS)^17^, and patterns of VS connectivity have been shown to predict substance use onset and frequency in adolescents and young adults^18,19^. Accumulating evidence further suggests that the facilitation of substance use behavior in adults may be driven by impaired inhibitory control, with distributed system-level alterations in cortical control regions (i.e., the vmPFC, cingulate, insula, and dorsolateral PFC^20,21^), potentially reflecting long-term dysregulation of top-down mechanisms over striatal DA processes as a core feature of substance use disorder and drug-seeking behavior^22–25^. Together, these findings implicate mesocorticolimbic networks underlying neurocognition and reward processing in substance use vulnerability. However, how developmental interactions between DA, inhibitory control, and impulsivity contribute to adolescent substance use – and function as neurodevelopmental markers that predict the time course of substance use across adolescence into adulthood – remains unclear.

Animal studies have shown dynamic changes within DAergic systems throughout adolescence, including increases in striatal DA availability^26,27^ and DA signaling^28^, that support enhanced reward drive, motivation, and risk-taking^27,29–31^, as well as drug-seeking behavior^28^. Human neuroimaging studies corroborate enhancements in reward-related VS blood oxygen level dependent (BOLD) activation^12,32–34^ during adolescence, which correspond to heightened risk-taking^32,35^, and have been shown to predict subsequent increases in substance use^13^. However, striatal BOLD is a distal proxy for functional activation of DA systems, and the lack of available non-invasive *in vivo* DA markers in pediatric populations remains an impediment to understanding its role in risk-taking and substance use behavior in youth. Therefore, much of our understanding concerning the neurobiology of substance use has come from molecular neuroimaging studies in adults, which have demonstrated distinct properties of DA signaling involved in different phases of substance use, as well as specific DAergic alterations following repeated exposure^36–40,40–42^. The few molecular imaging studies in youth have generally included small samples, primarily targeting young adulthood rather than childhood or adolescence, and have yielded mixed results – with studies showing either no changes (in D2 receptor availability and presynaptic DA release following binge drinking in young adults^43^) or blunted DA function (decreased D2 receptor availability and presynaptic DA release in young adults with familial risk for problematic substance use^44^). Based on these results, changes within D2 receptors and DA release could reflect a long-term consequence of increased midbrain DA biosynthesis in response to substance use (i.e., downregulation of striatal DA signaling)^20^. Alternatively, some animal studies have shown that lower baseline levels of striatal D2 receptor availability and DA release promotes increased drug-seeking behavior in animals with no prior history of exposure^45,46^, raising the possibility that these features could reflect predisposing risk markers. Given that substance use itself alters DA neurophysiology, it is difficult to discern from studies in adults whether DA system perturbations reflect preexisting vulnerability markers or long-term consequences of substance exposure. To this end, longitudinal studies in substance naïve youth are essential to identify individual differences in DAergic processes *prior* to the onset of substance use behavior, and would further allow for parsing out neurodevelopmental markers that may predict more long-term patterns of substance use (i.e., escalation into adulthood) from those associated with normative patterns of substance use that potentially reflect an instantiation of a more generalized adolescent risk-taking process.

Ascertaining a role for DA neurobiology in the development of substance use behavior in youth has been limited by restrictions on the use of PET in pediatric populations that provide direct estimates of DA. Magnetic Resonance (MR)-based tissue iron neuroimaging has emerged as a viable non-invasive index of DAergic physiology and maturation, as it can safely be obtained in pediatric cohorts. Scalable estimates of tissue iron can be obtained via normalizing and time-averaging T2*-weighted images (nT2*w) acquired during resting state scans, which quantifies the relative T2* relaxation across the brain and is sensitive to magnetic field inhomogeneities induced by iron ^47–49^. Tissue iron is involved in several neurodevelopmental processes, and is located in the dendrites of DA neurons ^50^, co-localizes with DA vesicles ^51^, and is involved in monoamine synthesis and production ^52–54^ as a co-factor for tyrosine hydroxylase ^51,55^, the rate limiting step in DA synthesis. Critically, tissue iron is found in highest concentrations within DAergic regions including the basal ganglia and midbrain ^56–59^, and MR indices of tissue iron have been shown to correspond to presynaptic vesicular DA storage measured using PET [^11^C]dihydrotetrabenazine (DTBZ)^60,61^. Additionally, neurodevelopmental studies have demonstrated increases in tissue iron accrual throughout the first two decades of life, with the rate of iron increase decelerating into adulthood ^62–66^, paralleling changes in DA availability observed in animal models ^67^. More recently, a role for basal ganglia tissue iron has been demonstrated in reward sensitivity ^68^, cognitive processing including inhibitory control^64,69,70^, and functional changes in mesocorticolimbic networks that support developmental decreases in risk-taking throughout adolescence^17^. Additionally, while initial studies have demonstrated a role for tissue iron, and related midbrain measures of dopamine neurophysiology (neuromelanin) in adolescent^71^ and young adult substance use^72^, critically, these studies have not characterized developmental trajectories in substance use to ascertain whether dopaminergic function serves as a neurobiological risk-marker for substance use escalation during the transition from adolescence to adulthood. Together, these results suggest that tissue iron reflects important aspects of DA-related neurophysiology, and changes in tissue iron play an important role in the specialization of networks that underlie adolescent risk-taking behavior. Here, we leverage these findings by examining the extent to which developmental variation in tissue iron, impulsivity, and inhibitory control predicts substance use behavior and distinct trajectories in substance use from adolescence to adulthood.

We leveraged a large, longitudinal, high-risk sample of adolescents and young adults, the majority of whom substance-naïve at baseline (having limited or no history of substance use at the first timepoint)^73^. We first characterized trajectories of substance use, as well as three critical markers that have been implicated in adult substance use and continue to undergo maturation in adolescence, namely, self-reported impulsivity, task-based assessments of inhibitory control, and MR-based estimates of basal ganglia tissue iron that provided *in vivo* estimates of basal ganglia DA-related neurophysiology. We then examined the extent to which individual differences (between-person) and developmental trajectories (within-person) of substance use corresponded to variation in impulsivity, inhibitory control, and tissue iron. Importantly, primary analyses examined substance use frequency and trajectories combined across multiple substances including nicotine, cannabis, and alcohol (including both all alcohol use and binge drinking), which allowed for the identification of neural and cognitive markers linked to adolescents’ general propensity toward substance use as a phenotypic risk-taking process, however, we also investigate each substance individually in order to assess substance-specific effects. Overall, capitalizing on repeated longitudinal sampling of brain and behavioral processes, the present study demonstrates that variation in impulsivity, inhibitory control, and DAergic processes plays a role in the timing and developmental progression of substance use behavior during the transition from adolescence to adulthood.

## Results

We examined how non-invasive neuroimaging markers of dopamine (DA) – related neurobiology and neurocognitive functioning shape patterns of substance use behavior from adolescence into adulthood in the National Consortium on Alcohol and Neurodevelopment in Adolescence and Adulthood (NCANDA-A) cohort, which combines yearly longitudinal neuroimaging with multidimensional assessments of executive function, trait impulsivity, and substance use in a large cohort of 807 adolescents (50% female) and young adults ages 12 – 30 across 1 – 9 longitudinal timepoints each, for a total of 6164 sessions. Importantly, the majority of participants were substance-naïve at baseline (with 83% having limited or no history of substance use at the first timepoint), and approximately 50% were recruited based on subclinical risk characteristics for alcohol use disorder^73^. Self-report substance use was assessed at each visit, with quantity assessed as the number of days of use in the past 30 days, frequency assessed as ‘never’, ‘ever’, or ‘regular (weekly)’, and intensity across multiple substances assessed as single-use, co-use (2 substances), or poly-use (>2 substances). Primary analyses examined generalized use patterns across all substances via a composite score, however, we also investigate substance-specific effects. Impulsivity was assessed using the UPPS-P Impulsive Behavior Scale^74–76^, and inhibitory control was assessed using the well-validated rewarded anti-saccade task^15,33,77^. Finally, basal ganglia DA – related neurobiology^61^ was assessed using MR-based estimates of tissue iron, obtained by time-averaged and normalized T2*-weighted imaging (nT2*w)^65,66^. Generalized Additive Mixed Models (GAMMs) were used in all primary analyses, allowing us to estimate flexible, data-driven trajectories and explore the shape of developmental trajectories for each measure, and to examine relationships between substance use and individual differences in impulsivity, inhibitory control, and DA-related neurobiology, controlling for non-linear age effects (see Methods). A series of participant-level sociodemographic sensitivity analyses explored how the magnitude, age-related trajectories, and developmental timing of substance use, neurocognitive, and DAergic metrics, varied as a function of 18 subgroups defined by sex, race/ethnicity, study site, and socioeconomic indicators. Finally, growth mixture models were used to identify distinct within-person trajectories in substance use across time, and GAMMs tested for individual differences neurocognitive and DAergic metrics across trajectory groups, and how developmental variation in these factors predicted patterns of youth substance use behavior.

### Patterns and Developmental Changes in Substance use, Impulsivity, Inhibitory Control, and Dopamine-Related Neurobiology

#### Substance Use Increases from Adolescence to Young Adulthood across the Full Sample

We first characterized patterns in the magnitude and developmental timing of past 30-day substance use across the full sample (Figure 1A – E). Use varied significantly by substance (*F* = 293.60, *p* < .001), with highest use (Figure 1A) and peak magnitudes (Figure 1Di) for cannabis, followed by alcohol, nicotine, and finally, binge drinking. GAMMs revealed non-linear age-related increases in generalized use patterns combined across all substances (composite use: s(age): F = 785.07, p < .001, Figure 1C & E), as well as across each substance individually (*p* < .001 in all 4 models; Figure 1C). However, substance-specific trajectories varied by shape (Figure 1C), age of initiation (Figure 1Dii), increase onset (Figure 1Diii), and rates of change (Figure 1Div). Cannabis and alcohol use showed earlier age of initiation, followed by steeper, more rapid age-related increases, while binge drinking and nicotine showed later age of initiation, followed by shallower, more gradual age-related increases. Peak use for all substances occurred around age 21–22 (Figure 1B).

**Figure 1.**
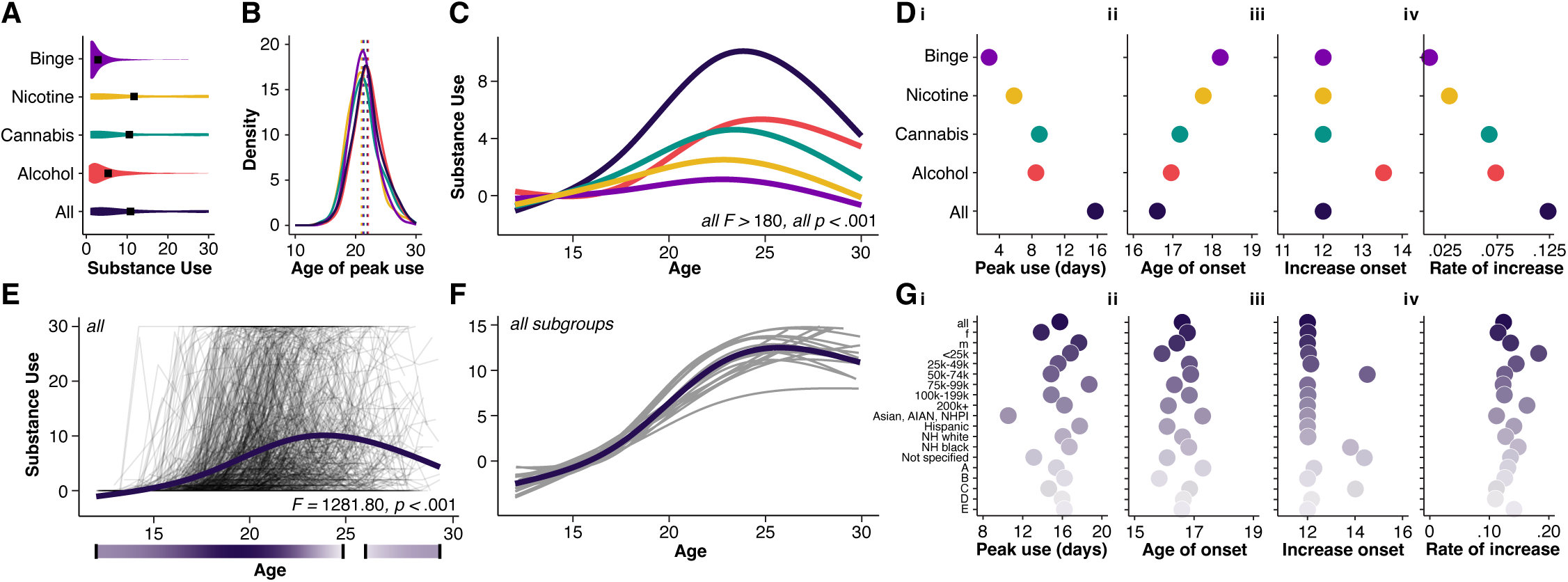
Patterns and developmental changes in substance use across the full sample and population subgroups. **(A)** Violin plots showing distribution of self-reported past 30-day substance use across individual substances as well as a composite measure across all substances (‘All’). Mean use is represented by black squares. **(B)** Histograms showing age of peak use across individual substances. Mean peak age is represented by hashed lines. **(C)** Nonlinear developmental models averaged across all visits for substance use for the full NCANDA sample (all participants), with each line representing individual substances (from A), and the dark line representing overall use across all substances. **(D) i.** Average peak use across individual substances (mean of per-participant peak value). **ii.** Average age of onset across individual substances. **iii.** Average age of increase onset across individual substances (first age where the confidence interval of the derivative did not include zero). **iv.** Rate of increase across individual substances (calculated as the average first derivative of the age spline). **(E)** Nonlinear developmental model for composite measure of substance use (as this forms the principal metric for analysis), with individual lines (black) reflecting connected visits for individual participants. **(F)** Overall substance use trajectories across population subgroups in NCANDA, including 18 population subgroups reflecting race/ethnicity, household income, guardian education, sex, and study site (geographical location), analyzed individually (grey lines, full sample is dark line). (**G) i.** Average peak overall substance use across population subgroups (maximum value from nonlinear fit). **ii.** Average age of increase onset in overall substance use across population subgroups (first age where the confidence interval of the derivative did not include zero). **iii.** Average age of onset across individual substances. **iv.** Rate of increase in overall substance use across population subgroups (calculated as the average first derivative of the age spline). Female (F). Male (M). American Indian and Alaska Native (AIAN). Native Hawaiian and Pacific Islander (NHPI). Non-Hispanic (NH).

#### The Developmental Timing, but not Magnitude, of Substance Use is Consistent across Sociodemographic Factors

We next examined whether developmental patterns in general substance use (composite) varied by sociodemographic factors (see Methods). Substance use significantly differed by sex (*F* = 7.23, *p* = .007), but not race/ethnicity, income, or study site (all *p* > .05). The *peak* magnitude of use (Figure 1Gi) differed significantly by sex (*F* = 21.83, *p* < .001) and race/ethnicity (*F* = 4.95, *p* < .001), but not income or study site (all *p* > .05). Independent models across the 18 population subgroups demonstrated consistency in developmental timing (Figure 1F); with ages of initiation ranging from 15.5 and 17.5 years in all subgroups (Figure 1Gii), developmental increases beginning between approximately 12 and 15 years in 17 of 18 population subgroups (Figure 1Giii), ages of peak use between approximately 20.5 and 22.5 years in all subgroups (not shown), and comparable rates of change (Figure 1Giv).

#### Impulsivity Decreases from Adolescence to Young Adulthood across the Full Sample

We assessed patterns in the magnitude and developmental timing of trait-level impulsivity across the full sample (Figure 2A – D). Impulsivity varied significantly by subscale (*F* = 3434.83, *p* < .001), with highest scores (Figure 2A) and peak scores (Figure 2Di) for sensation seeking, followed by negative urgency, perseverance, positive urgency, and finally, premeditation. GAMMs revealed linear age-related decreases in mean impulsivity (across all subscales: s(age): F = -75.66, p < .001, Figure 2C & E), as well as across each subscale individually (*p* < .001 in all 5 models; Figure 2C). However, subscale-specific trajectories varied by shape (Figure 2C), decrease onsets (Figure 2Dii), and rates of change (Figure 2Diii). Positive urgency, premeditation, and negative urgency began decreasing around age 12, whereas perseverance and sensation seeking showed protracted decrease onsets at approximately age 15 and 18, respectively. In comparison to other subscales, perseverance showed shallower and slower decreases over time. Peak impulsivity for all subscales occurred around age 18 (Figure 2B).

**Figure 2.**
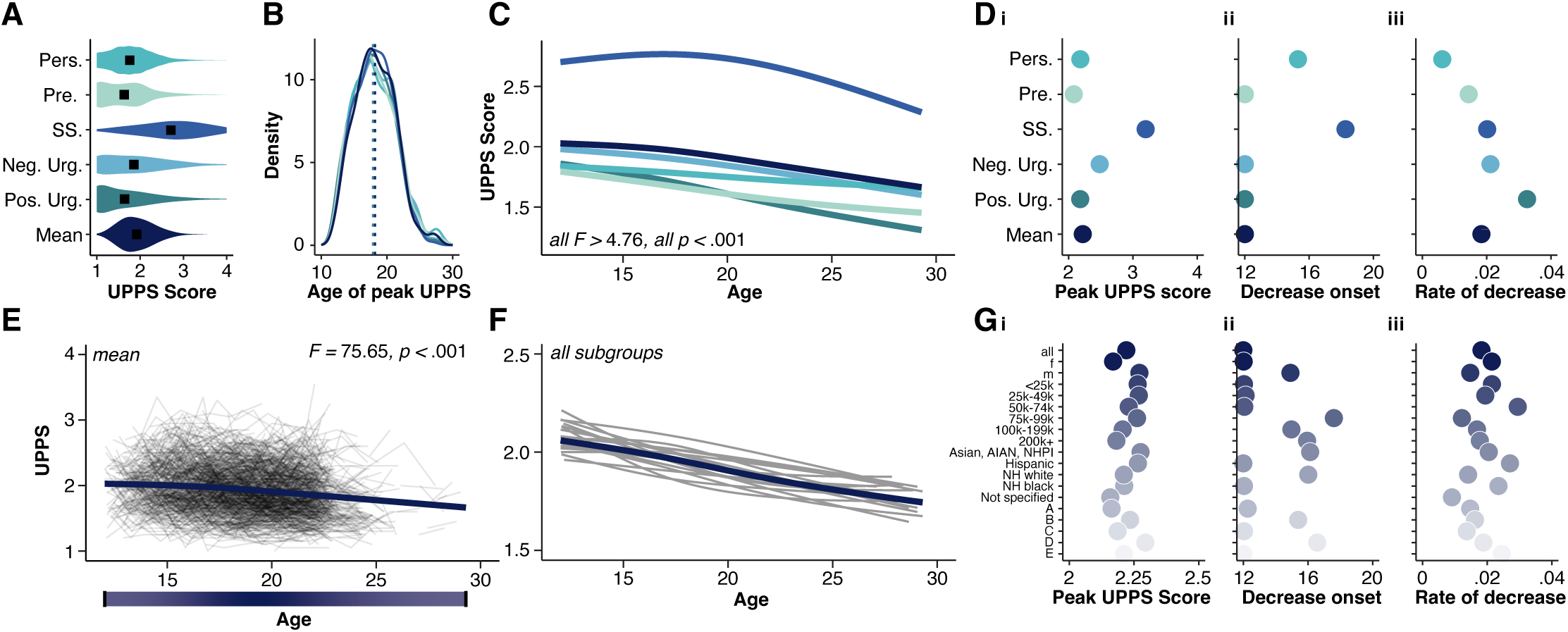
Patterns and developmental changes in impulsivity across the full sample and population subgroups. **(A)** Violin plots showing distribution of self-reported impulsivity, measured by the UPPS, across individual subscales as well as a the mean across all subscales. Mean scores are represented by black squares. **(B)** Histograms showing age of peak UPPS across individual subscales. Mean peak age is represented by hashed lines. **(C)** Nonlinear developmental models averaged across all visits for UPPS scores for the full NCANDA sample (all participants), with each line representing individual subscales (from A), and the dark line representing the mean across all subscales. **(D) i.** Average peak UPPS across individual subscales (mean of per-participant peak value). **ii.** Average age of decrease onset across individual subscales (first age where the confidence interval of the derivative did not include zero). **iii.** Rate of decrease across individual substances (calculated as the average first derivative of the age spline). **(E)** Nonlinear developmental model for mean UPPS scores (as this forms the principal metric for analysis), with individual lines (black) reflecting connected visits for individual participants. **(F)** Mean UPPS trajectories across population subgroups in NCANDA, including 18 population subgroups reflecting race/ethnicity, household income, guardian education, sex, and study site (geographical location), analyzed individually (grey lines, full sample is dark line). **(G) i.** Average peak mean UPPS score across population subgroups (maximum value from nonlinear fit). **ii.** Average age of decrease onset in mean UPPS across population subgroups (first age where the confidence interval of the derivative did not include zero). **iii.** Rate of decrease in mean UPPS scores across population subgroups (calculated as the average first derivative of the age spline). Female (F). Male (M). American Indian and Alaska Native (AIAN). Native Hawaiian and Pacific Islander (NHPI). Non-Hispanic (NH).

#### The Developmental Timing, but not Magnitude, of Impulsivity is Consistent across Sociodemographic Factors

Impulsivity (mean across all subscales) significantly differed by sex (*F* = 23.63, *p* < .001), race/ethnicity (*F* = 4.07, *p* < .001), and study site (*F* = 2.87, *p* < .001), but not income (*F* = .25, *p* = .94). The *peak* magnitude of impulsivity (Figure 2Gi) differed significantly by sex (*F* = 13.57, *p* < .001) and study site (*F* = 2.83, *p* = .02) but not race/ethnicity or income (all *p* > .05). Independent models across the 18 population subgroups demonstrated relatively consistent developmental timing (Figure 2F), with developmental decreases beginning between approximately 12 and 16 years of age in 15 of 18 population subgroups (Figure 2Gii), ages of peak impulsivity between approximately 17.5 and 19 years of age in all subgroups (not shown), and comparable rates of change (Figure 2Giii).

#### Inhibitory Control Performance Increases from Adolescence to Young Adulthood across the Full Sample

Task-based measures of inhibitory control were obtained using the anti-saccade task, which is a visuospatial inhibitory control task in which participants are presented with a peripheral cue and required to direct their gaze to its mirror location, requiring the suppression of the reflexive response to look toward the stimulus and the generation of a voluntary eye movement in the opposite direction (Figure 3A; see Methods, note that the anti-saccade task was administered at 2 participating research sites). Reward was administered on half of the trials (‘rewarded trials’) and the other half were non-rewarded (‘neutral’ trials), and accuracy was assessed as the percentage of correct trials.

**Figure 3.**
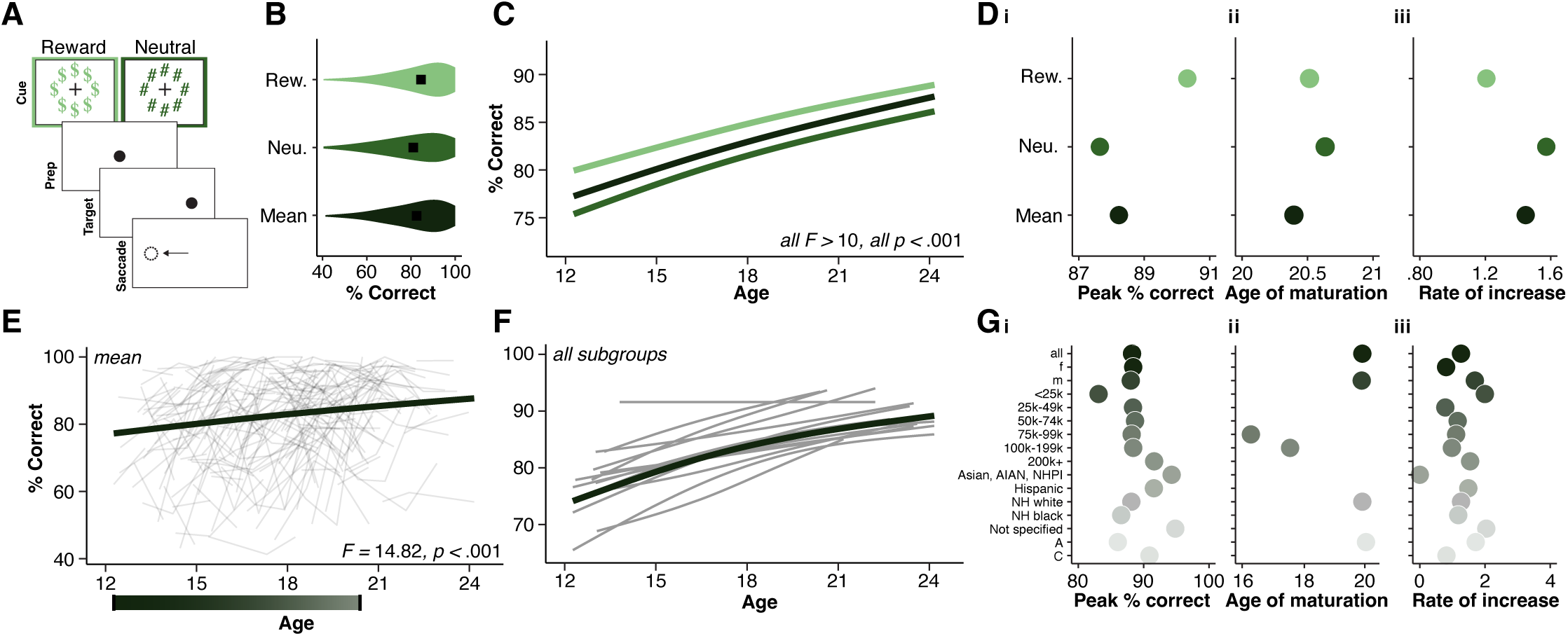
Patterns and developmental changes in inhibitory control across the full sample and population subgroups. **(A)** Anti-Saccade task. Each trial began with a cue epoch during which a fixation point was surrounded by either a ring of dollar signs, indicating the chance to earn reward for correct performance (‘reward’ trials), or a ring of pound signs, indicating that there was no reward at stake for the trial (‘neutral trials’), followed up a response preparation period. Next, a peripheral cue appeared in an unknown location along the horizontal midline and participants were instructed to direct their gaze to the mirror location of the stimulus, completing an anti-saccade. Each trial was followed by a variable intertrial interval (ITI; not depicted). **(B)** Violin plots showing distribution of anti-saccade performance (% correct), across individual trial types (reward and neutral) as well as a the mean across trial types. Mean scores are represented by black squares. (**C)** Nonlinear developmental models averaged across all visits for anti-saccade performance for the participating NCANDA sample who performed the anti-saccade task (Duke and Pittsburgh sites only), with each line representing individual trial types (from A), and the dark line representing the mean across all trials. **(D) i.** Average peak anti-saccade performance across individual trial types (mean of per-participant peak value). **ii.** Average age of maturation across individual trial types (first age where the confidence interval of the derivative included zero). **iii.** Rate of increase across individual trial types (calculated as the average first derivative of the age spline). **(E)** Nonlinear developmental model for mean anti-saccade performance (as this forms the principal metric for analysis), with individual lines (black) reflecting connected visits for individual participants. **(F)** Mean anti-saccade trajectories across population subgroups in NCANDA, including 15 population subgroups reflecting race/ethnicity, household income, guardian education, sex, and study site (geographical location, note only two participating sites), analyzed individually (grey lines, full sample is dark line). **(G) i.** Average peak mean anti-saccade performance across population subgroups (maximum value from nonlinear fit). **ii.** Average age of increase onset in mean anti-saccade performance across population subgroups (first age where the confidence interval of the derivative did not include zero). **iii.** Rate of increase in overall anti-saccade performance across population subgroups (calculated as the average first derivative of the age spline). Female (F). Male (M). American Indian and Alaska Native (AIAN). Native Hawaiian and Pacific Islander (NHPI). Non-Hispanic (NH).

We assessed patterns in the magnitude and developmental timing of inhibitory control across the full sample (Figure 3B – D). Anti-saccade performance varied significantly by trial type (*F* = 46.85, *p* < .001), with highest performance (Figure 3B) and peak performance (Figure 3Di) for reward trials in comparison to neutral trials. GAMMs revealed inverse age-related increases in mean anti-saccade performance (across all trial types; s(age): F = 14.82, p < .001, Figure 3C & E), as well as across each trial type individually (*p* < .001 in both models; Figure 3C). Trial-type specific trajectories did not vary by shape (Figure 3C), increase onsets, age of maturation (Figure 3Dii), age of peak performance, or rates of change (Figure 3Diii). Note that we show age of maturation (Figure 3Dii) instead of age of increase onset given common increase onsets across trial types and given that the age at which inhibitory control reaches its developmental plateau is more insightful.

#### The Developmental Timing and Magnitude of Response Inhibition is Consistent across Sociodemographic Factors

Anti-saccade performance (mean across trial types) significantly differed across study site (*F* = 32.01, *p* < .001) and race/ethnicity (*F* = 2.57, *p* = .04), but not sex or income (all *p* > .05). The *peak* magnitude of inhibitory control performance (Figure 3Gi) similarly differed significantly across sex (*F* = 13.57, *p* < .001) and study site (*F* = 2.83, *p* = .02), but not race/ethnicity, or income (all *p* > .05). Independent models across the 15 population subgroups (because anti-saccade task data were only collected across two participating sites, see Methods) were less statistically powered in comparison to other metrics, thus we were only able to detect significant periods of change in 5 of the 15 population subgroups. However, independent models across these 5 population subgroups revealed relatively consistent developmental timing (Figure 3F), with developmental increases beginning between approximately 12 and 16 years of age in 4 of the 5 subgroups (not shown), stabilizing between 17.5 and 20 years of age in 4 of 5 population subgroups (Figure 3Gii), ages of peak performance between approximately 16.5 and 18.5 years of age in all subgroups (not shown), and comparable rates of change (Figure 3Giii).

#### Basal Ganglia Tissue Iron Decreases from Adolescence to Young Adulthood across the Full Sample

MR-based indices of basal ganglia tissue iron, reflecting DA-related neurobiology^61^ were obtained via time-averaged and normalized T2*-weighted imaging (nT2*w) across a 10-minute resting state scan, and individual estimates were extracted across each basal ganglia region of interest, including the caudate nucleus, nucleus accumbens (NAcc), pallidum, and putamen, as well as one estimate reflecting the mean across all voxels in the basal ganglia (Figure 4A; see Methods, note that nT2*w is inversely related to iron content).

**Figure 4.**
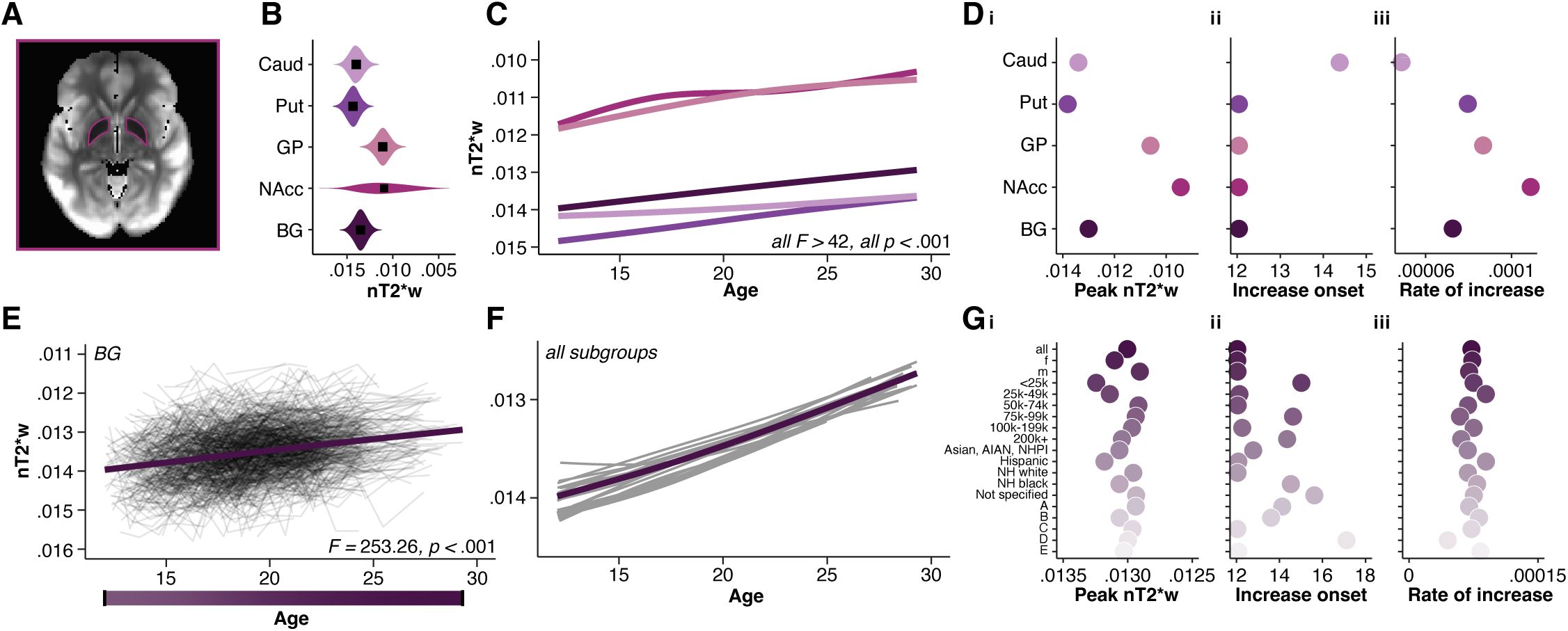
Patterns and developmental changes in tissue iron reflecting dopamine-neurobiology across the full sample and population subgroups. **(A)** Mean nT2*w data across all visits and all participants. **(B)** Violin plots showing distribution of tissue iron, measured by time-averaged and normalized T2*-weighted images (nT2*w), across individual basal ganglia regions of interest (ROI), as well as the mean across all voxels in the basal ganglia. Mean values are represented by black squares. **(C)** Nonlinear developmental models averaged across all visits for nT2*w for the full NCANDA sample (all participants), with each line representing individual ROIs (from A), and the dark line representing the whole basal ganglia. **(D) i.** Average peak nT2*w across individual ROIs (mean of per-participant peak value). **ii.** Average age of increase onset across individual ROIs (first age where the confidence interval of the derivative did not include zero). **iii.** Rate of increase across individual ROIs (calculated as the average first derivative of the age spline). **(E)** Nonlinear developmental model for basal ganglia nT2*w (as this forms the principal metric for analysis), with individual lines (black) reflecting connected visits for individual participants. **(F)** Basal ganglia nT2*w trajectories across population subgroups in NCANDA, including 18 population subgroups reflecting race/ethnicity, household income, guardian education, sex, and study site (geographical location), analyzed individually (grey lines, full sample is dark line). **(G) i.** Average peak basal ganglia nT2*w across population subgroups (maximum value from nonlinear fit). **ii.** Average age of increase onset in basal ganglia nT2*w across population subgroups (first age where the confidence interval of the derivative did not include zero). **iii.** Rate of increase in basal ganglia nT2*w across population subgroups (calculated as the average first derivative of the age spline). Note that x-axes in (B), (D), (I), and y-axes in (C), (G), and (H) are reversed for visualization to ease interpretability given the inverse relationship between nT2*w and tissue iron concentration. Time-averaged and normalized T2*w (nT2*w). Caudate (Caud). Putamen (Put). Globus Pallidum (GP). Nucleus Accumbens (NAcc). Basal ganglia (BG). Female (F). Male (M). American Indian and Alaska Native (AIAN). Native Hawaiian and Pacific Islander (NHPI). Non-Hispanic (NH).

We assessed patterns in the magnitude and the developmental timing of nT2*w across the full sample (Figure 5B – D). As expected, nT2*w varied significantly by region (*F* = 11281.35, *p* < .001), with lowest nT2*w, indicating highest iron, (Figure 4B) and lowest peak nT2*w (Figure 4Di) for the NAcc, followed by the pallidum, the caudate, and finally the putamen. GAMMs revealed age-related decreases in nT2*w in basal ganglia nT2*w (mean across all voxels; *F* = -253.25, *p* < .001, Figure 4C & E), as well as across each region individually (*p* < .001 in all 4 models; Figure 4C). Region-specific trajectories varied by shape (Figure 4C), decrease onsets (Figure 4Dii), rates of change (Figure 4Diii), and age of peak nT2*w. The caudate showed shallower, later, and more gradual age-related decreases over time beginning around age 14, while the NAcc showed steeper age-related decreases in comparison to other regions. Additionally, to ascertain the age at which the most rapid developmental changes in tissue iron occurred, we examined peak ages of change, which were earliest for the NAcc and the pallidum (around age 13, not shown), in comparison to other basal ganglia regions (around age 19 – 20) reflecting relatively early, rapid changes within these regions. All regions showed peak nT2*w values in young adulthood around age 19-20.

**Figure 5.**
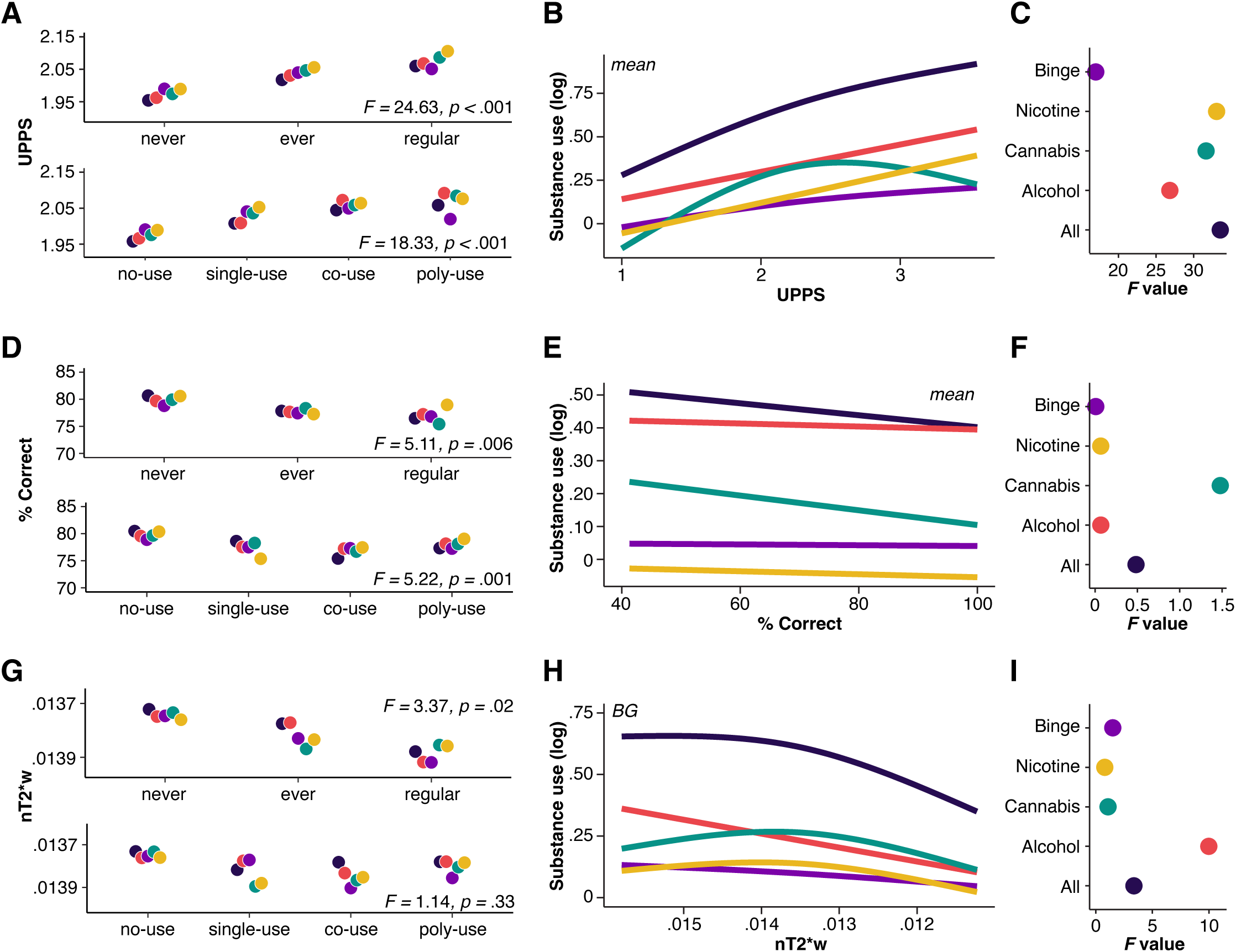
Associations between substance use and impulsivity, response inhibition, and tissue iron across the full sample. **(A) Top panel**: predicted UPPS values within substance use frequency categories. **Bottom panel:** predicted UPPS values within substance use use-type categories. Statistics represent a main effect of frequency and use-type category on UPPS scores from an ANOVA test, controlling for age and sociodemographic variables. **(B)** Predicted values from non-linear models assessing associations between substance use and impulsivity, measured by mean UPPS scores, averaged across all visits for the full NCANDA sample (all participants). **(C)** F-values reflecting significance of the smooth term for UPPS for each individual substance. **(D) Top panel:** predicted anti-saccade performance within substance use frequency categories. **Bottom panel:** predicted anti-saccade performance within substance use use-type categories. Statistics represent a main effect of frequency and use-type category on anti-saccade performance from an ANOVA test, controlling for age and sociodemographic variables. **(E)** Predicted values from non-linear models assessing associations between substance use and response inhibition, measured by mean performance on the anti-saccade task, averaged across all visits for the NCANDA sample (at participating sites). **(F)** F-values reflecting significance of the smooth term for anti-saccade performance for each individual substance. **(G) Top panel:** predicted basal ganglia nT2*w within substance use frequency categories. **Bottom panel:** predicted basal ganglia nT2*w within substance use use-type categories. Statistics represent a main effect of frequency and use-type category on nT2*w from an ANOVA test, controlling for age and sociodemographic variables. **(H)** Predicted values from non-linear models assessing associations between substance use and tissue iron, measured by basal ganglia nT2*w, averaged across all visits for the full NCANDA sample (all participants). **(I)** F-values reflecting significance of the smooth term for basal ganglia nT2*w for each individual substance. In (B), (E), and (H), y-axes show log-transformed past 30-day substance use, and each line represents an individual substance, and the dark line represents overall use across all substances. In (A), (D), and (G), categorizations in the top panel reflect whether participants reported never having used substances (‘never’), ever having used substances (‘ever’), and regularly using substances (>1/week) at each visit, and statistics report a main effect of frequency category. Categorizations in the bottom panel reflect whether participants reported no-use (i.e., never), the use of one substance (‘single-use’), the use of two substances (‘co-use’), or greater than two substances (‘poly-use’) at each visit, and statistics report a main effect of use-type category. Note that the y-axis in (G) and the x-axis in (H) is reversed for visualization to ease interpretability given the inverse relationship between nT2*w and tissue iron concentration.

#### The Developmental Timing, but not Magnitude, of Basal Ganglia Tissue Iron is Consistent across Sociodemographic Factors

Basal ganglia nT2*w significantly differed by sex (*F* = 51.94, *p* < .001) and race/ethnicity (*F* = 4.76, *p* < .001), but not income or study site (all *p* > .05). The *peak* magnitude of basal ganglia nT2*w (Figure 4Gi) differed significantly by sex (*F* = 27.77, *p* < .001), race/ethnicity (*F* = 2.54, *p* < .04), and income (*F* = 3.06, *p* = .01) but not study site (*F* = 1.38, *p* = .24). Independent models across the 18 population subgroups demonstrate consistent developmental timing of basal ganglia nT2*w, with developmental decreases, indicating increases in tissue iron, beginning between 12 and 15 years of age in 16 of 18 population subgroups (Figure 4Gii), ages of peak nT2*w between approximately 19.5 and 21 years of age in all subgroups (not shown), and comparable rates of change (Figure 4Giii).

Together, these results demonstrate robust, longitudinal changes in substance use, trait-level impulsivity, inhibitory control, and basal ganglia tissue iron across the full sample, with variability in the magnitude of substance use, impulsivity, inhibitory control, and basal ganglia tissue iron, but largely conserved developmental timing, across sociodemographic contexts. See Supplementary Table S1, S3, & S5 for full model statistics across individual substances, UPPS subscales, anti-saccade trial types, basal ganglia regions, and Supplementary Table S2, S4, & S6 for population subgroups.

### Increased Substance Use is Associated with Heightened Impulsivity, Decreased Inhibitory Control, and Reduced Basal Ganglia Tissue Iron

To assess whether individual differences in neurocognitive and neurobiological metrics were associated with qualitative shifts in use patterns, substance use was grouped into categories at each visit reflecting substance use frequency (i.e., never, ever, regular; see above) and use-type (i.e., no-use, single-use, co-use, poly-use indicating higher use across multiple substances; see Table S7A for group characteristics). Models then interrogated individual differences in impulsivity, inhibitory control, and tissue iron within each category, and associations with past 30-day substance use. All models controlled for a smoothed term of age, allowing us to investigate these relationships above and beyond age effects.

#### Impulsivity

Impulsivity (mean across subscales) significantly differed across substance use frequency categories (Figure 5A top panel; *F* = 24.63, *p* < .001), with higher scores among those who reported regular use in comparison to never and ever, and higher scores in those who report ever using compared to never (all *p* < .001). Impulsivity also significantly differed across substance use-type categories (Figure 5A bottom panel; *F* = 18.33, *p* < .001), with higher scores associated with poly-substance use (3+) in comparison to no-use and single substance-use, higher scores in co-substance use in comparison to no-use and single-substance use, and higher scores in single-substance use in comparison to no-use (all *p* < .01). Finally, impulsivity was significantly associated with past 30-day substance use (Figure 5B-C; *F* = 33.54, *p* < .001), with higher UPPS scores associated with increased days of use. These effects were similar across all individual UPPS subscales (see Methods for lack of interaction effects).

#### Inhibitory Control

Inhibitory control (mean across trial types) significantly differed across substance use frequency categories (Figure 5D top panel; *F* = 5.11, *p* = .006), with lower performance in those who report regular use in comparison to never and ever (all *p* < .01). Inhibitory control performance also significantly differed across substance use-type categories (Figure 5D bottom panel; *F* = 5.22, *p* = .001), with lower anti-saccade performance in poly-substance use in comparison to no-use (*p* = .02), and lower performance in co-substance use in comparison to no-use and single-substance use (both *p* < .01). In contrast, we did not observe a significant association between mean anti-saccade performance and past 30-day substance use (Figure 5E-F; *F* = .48, *p* = .49). These effects were similar across anti-saccade trial types (see Methods for lack of interaction effects).

#### Basal Ganglia DA-Related Neurobiology

Basal ganglia nT2*w significantly differed across substance use frequency categories (Figure 5G top panel; *F* = 3.37, *p* = .02), with higher nT2*w, indicating lower iron, in those who report regular use in comparison to never and ever (all *p* < .01), but no significantly difference in nT2*w across substance use-type categories (Figure 5G bottom panel; *F* = 1.14, *p* = .33). Finally, basal ganglia nT2*w was significantly associated with past 30-day substance use (Figure 5H-I;*F* = 4.22, *p* = .01), with higher nT2*w, indicating lower iron, associated with increased days of use. These effects were similar across individual ROIs (see Methods for lack of interaction effects).

All together, these results suggest that increased substance use was associated with heightened impulsivity, decreased inhibitory control, and lower basal ganglia tissue iron. See Supplementary Table S7 for full model statistics across individual substances, UPPS subscales, anti-saccade trial types, and basal ganglia regions of interest.

### Examining Within-Person Patterns of Substance use and Relationships with Impulsivity, Inhibitory Control, and Dopamine-Related Neurobiology

In order to understand whether individual differences in impulsivity, inhibitory control, and dopamine-related neurobiology contributed to the developmental time course of substance use from adolescence to young adulthood, we first characterized subgroups reflecting within-person trajectories in substance use, and next investigated individual differences across trajectory groups, as well as developmental variation in these factors across trajectories that might indicate that individual differences occurred early in development (i.e., prior to substance use onset) or later in development (i.e., following substance exposure).

#### Characterizing Variation in Within-Person Trajectories in Substance Use

Growth Mixture Models (GMMs) were applied to capture variation in within-person trajectories (patterns of change) across timepoints and classify individuals based on generalized patterns of overall substance use (composite) across development (Figure 6A – D). These models revealed four classes of common trajectories in substance use across the full sample (Figure 6A, top panel), all of which showed significant age-related increases in substance use (*p* < .001 in all 4 models; Figure 6A). However, trajectory groups differed in the magnitude of substance use (Figure 6B; *F* = 113.88, *p* < .001), and in the shape (Figure 6A), age of initiation (Figure 6Dii), age of peak use (Figure 6C), increase onsets (Figure 6Diii), and rate of change (Figure 6Div). The *low* group (n = 241, 30% of participants) showed only minimal increases across adolescence (mean age of initiation: 17.77 years), with near-zero growth. The *youth peak* group (n = 139, 17% of participants), comprised of individuals who showed the highest level of substance use, showed a quadratic pattern, with steep, rapid age-related increases beginning around age 12 (mean age of initiation: 15.65 years), followed by declines into young adulthood beginning around age 26. In contrast, the *adolescent increasing* group (n = 139, 17% of participants) showed steep, linear increases beginning around age 16 (mean age of initiation: 16.39 years). Finally, the *adult increasing* group (n = 211, 26% of participants) showed slower, more gradual linear increases beginning around age 16 (mean age of initiation: 17.38 years), and a later age of peak use (around age 23) in comparison to other groups, which peaked earlier (around age 21 – 22). Together, these findings highlight distinct developmental patterns of substance use across trajectory groups.

**Figure 6.**
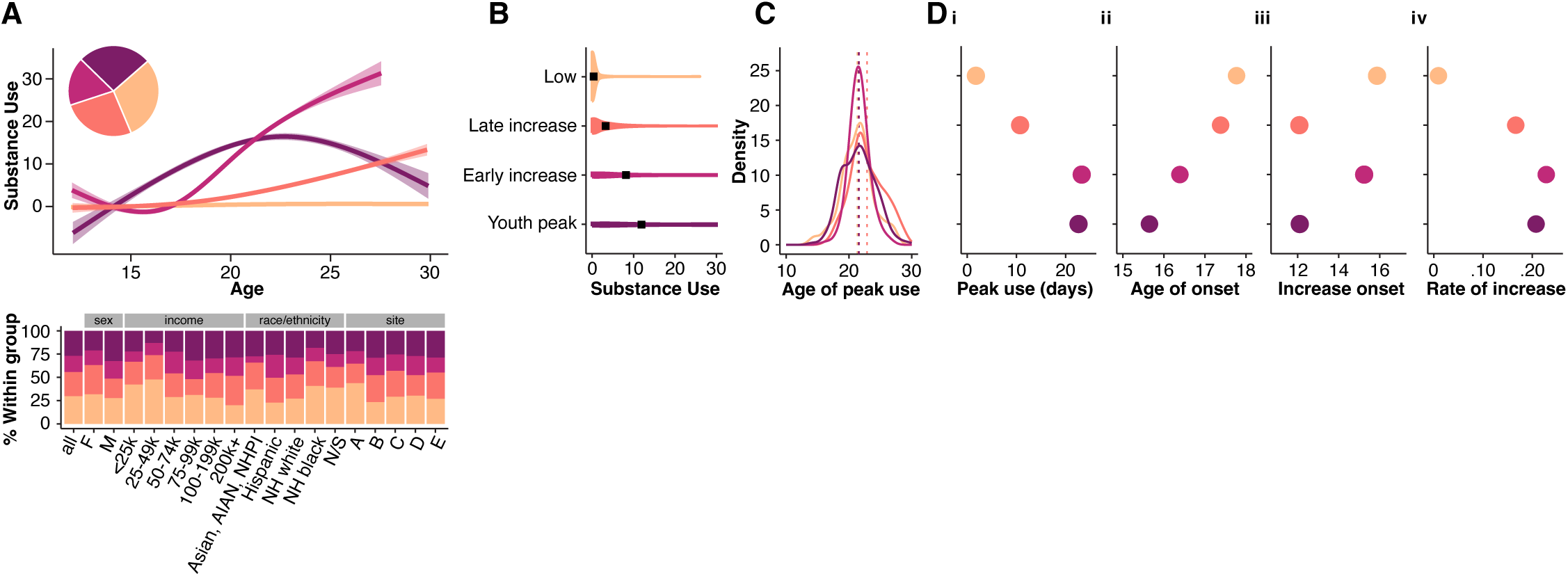
Distinct within-person trajectories in substance use. **(A) Top panel:** Growth Rate Mixture Models revealed four distinct patterns of within-person trajectories in substance use, including a *low* group, comprised of individuals who show no- or perpetually low levels of substance use across visits (n = 241, 30% of participants), *youth peak*, comprised of individuals who showed the earliest increases, with peak use in adolescence/young adulthood followed by decreases into adulthood (n = 211, 26% of participants), and finally, two *increasing* groups; an *adolescent increasing* group comprised of individuals who showed early, steep increases in use from adolescence into adulthood (n = 139, 17% of participants) and an *adult increasing* group comprised of individuals who showed low use in adolescence followed by linear increases in use into adulthood (n = 211, 26% of participants). The main plot shows nonlinear developmental models for overall substance use for individual trajectory groups. The pie chart indicates the percentage of participants within each trajectory subgroup; color in pie chart corresponds to age model color in main plot. **Bottom panel:** Group membership within each trajectory group across population subgroups in NCANDA, including 18 population subgroups reflecting race/ethnicity, household income, guardian education, sex, and study site (geographical location). **(B)** Violin plots showing distribution of self-reported past 30-day overall substance use across individual trajectory group. Mean use is represented by black squares. **(C)** Histograms showing age of peak use across individual substances. Mean peak age is represented by hashed lines. **(D) i.** Average peak use across individual trajectory groups (mean of per-participant peak value). **ii.** Average age of onset across individual trajectory groups. **iii.** Average age of increase onset across individual trajectory group (first age where the confidence interval of the derivative did not include zero). **iv.** Rate of increase across individual trajectory groups (calculated as the average first derivative of the age spline). X-axis reflects the proportion of each trajectory group across each demographic factor. Female (F). Male (M). American Indian and Alaska Native (AIAN). Native Hawaiian and Pacific Islander (NHPI). Non-Hispanic (NH).

Importantly, sensitivity analyses illustrate that group membership was not significantly associated with variation in sociodemographic factors (Figure 6A, bottom panel; i.e., that certain sociodemographic factors did not disproportionately fall into a particular trajectory category). While we found a significant effect of age on trajectory groups (*F* = 19.90, *p* < .001), there was only minor variation in age among trajectory groups (ranging from 18.8 years to 20.5 years).

#### Distinct Developmental Patterns of Impulsivity across Trajectory Groups

Trait-level impulsivity (mean across subscales) significantly differed across trajectory groups (Figure 7B; *F* = 14.13, *p* < .001), highest scores in the *youth peak* group, followed by the *adolescent increasing* group, the *low* group, and lastly, the *adult increasing* group. Independent GAMMs modelling the developmental time course of impulsivity within each trajectory group (Figure 7A – C) revealed significant age-related decreases in all groups (*p* < .001 in all 4 models; Figure 7A). However, trajectories differed in shape (Figure 7A), decrease onsets (Figure 7Cii), and rate of change (Figure 7Ciii). The *low, youth peak,* and *adult increasing* groups showed steep, early decreases beginning early in adolescence. In contrast, the *adolescent increasing* group showed later, shallower, and slower decreases.

**Figure 7.**
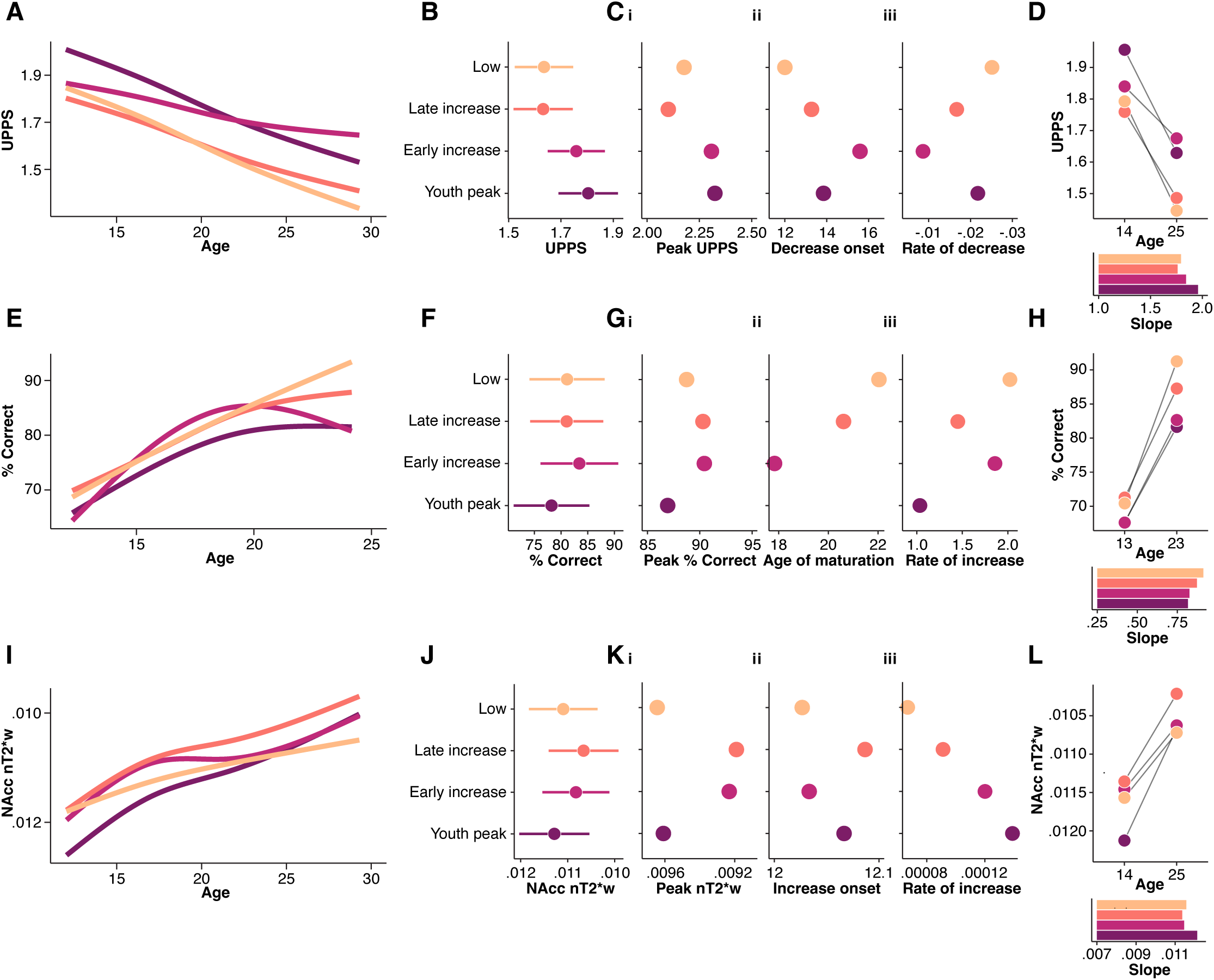
Variation in impulsivity, response inhibition, and tissue iron within substance use trajectory subgroups. **(A)** Nonlinear developmental models for mean UPPS scores for individual substance use trajectory groups. **(B)** Predicted mean UPPS scores within individual substance use trajectory groups. **(C) i.** Average peak mean UPPS scores across individual substance use trajectory groups (mean of per-participant peak value). **ii.** Average age of decrease onset across individual substance use trajectory groups (first age where the confidence interval of the derivative did not include zero). **iii.** Rate of decrease across individual substance use trajectory groups (calculated as the average first derivative of the age spline). **(D) Top panel:** Predicted mean UPPS scores at age 14 (i.e., beginning of study) and age 25 (i.e., end of study) for individual substance use trajectory groups. **Bottom panel:** Predicted slope of age-related change in mean UPPS score for individual substance use trajectory groups. **(E)** Nonlinear developmental models for mean anti-saccade performance for individual substance use trajectory groups. **(F)** Predicted mean anti-saccade performance within individual substance use trajectory groups. **(G) i.** Average peak mean anti-saccade performance across individual substance use trajectory groups (mean of per-participant peak value). **ii.** Average age of maturation across individual substance use trajectory groups (first age where the confidence interval of the derivative included zero). **iii.** Rate of increase across individual substance use trajectory groups (calculated as the average first derivative of the age spline). **(H) Top panel:** Predicted mean anti-saccade performance at age 13 (i.e., beginning of study) and age 23 (i.e., end of study) for individual substance use trajectory groups. **Bottom panel:** Predicted slope of age-related change in mean anti-saccade performance for individual substance use trajectory groups. **(I)** Nonlinear developmental models for NAcc nT2*w for individual substance use trajectory groups. **(J)** Predicted NAcc nT2*w within individual substance use trajectory groups. **(K) i.** Average peak NAcc nT2*w across individual substance use trajectory groups (mean of per-participant peak value). **ii.** Average age of increase onset across individual substance use trajectory groups (first age where the confidence interval of the derivative did not include zero). **iii.** Rate of increase across individual substance use trajectory groups (calculated as the average first derivative of the age spline). **L. Top panel:** Predicted NAcc nT2*w at age 14 (i.e., beginning of study) and age 25 (i.e., end of study) for individual substance use trajectory groups. **Bottom panel:** Predicted slope of age-related change in NAcc nT2*w for individual substance use trajectory groups. Nucleus accumbens (NAcc). Time-averaged and normalized T2*w (nT2*w). Note that the y-axis in (M), and the top panel of (R), and the x-axes in (N) and (O) are reversed for visualization to ease interpretability given the inverse relationship between nT2*w and tissue iron concentration.

##### Variation in Impulsivity across Age within Trajectory Groups

A significant trajectory group by age interaction emerged for impulsivity (*F* = 9.89, *p* < .001), and post-hoc tests revealed in early adolescence (estimates for age 14, see methods, Figure 7D top panel), the *youth peak* group showed significantly higher predicted UPPS scores in comparison to all other trajectory groups (all *p*s < .01), and the *adolescent increasing* group showed significantly higher scores in comparison to the *adult increasing* group (*p* = .03). By young adulthood (estimates for age 25, see methods, Figure 7D top panel), UPPS scores remained higher in the *youth peak* and *adolescent increasing* groups in comparison to both the *low* and *adult increasing* groups (all *p*s < .001). Additionally, the *adolescent increasing* group showed significantly flatter slopes (Figure 7D bottom panel), indicating diminished age-related decreases in impulsivity over time (all *p*s < .05) in comparison to other groups. In contrast, the *youth peak* group showed comparable slopes to the *low* and *adult increasing* groups, potentially reflecting normative decreases in impulsivity over time. Together, these findings highlight distinct developmental patterns of impulsivity across trajectory groups, and suggest early and persistent elevations in impulsivity, particularly in the *youth peak* and *adolescent increasing* groups, with the *adolescent increasing* group also showing weaker developmental declines in impulsivity.

#### Distinct Developmental Patterns of Inhibitory Control across Trajectory Groups

Inhibitory control performance (mean across trial types) did not differ significantly across substance use trajectory categories (Figure 7F; *F* = 1.52, *p* = .21), however, the *youth peak* group tended to have lower inhibitory control performance, compared to the *low*, *adolescent increasing*, and *adult increasing* groups. Independent GAMMs modelling the developmental time course of inhibitory control within each trajectory group (Figure 7E – H) revealed significant age-related increases in all groups (*p* ranging from .01 to <.001). However, trajectories differed in shape (Figure 7E), increase onsets (Figure 7Gii), age of maturation (Figure 7Gii), and rate of change (Figure 7Giii). The *low, adult increasing,* and *adolescent increasing* groups showed steep, rapid, age-related increases throughout adolescence, followed by stabilization in young adulthood. In contrast, the *youth peak* group slower and overall attenuated age-related increases in comparison to other groups. Note that we show age of maturation (Figure 7Dii) instead of age of increase onset given common increase onsets across trial types and given that the age at which inhibitory control reaches its developmental plateau is more insightful.

##### Variation in Inhibitory Control across Age within Trajectory Groups

We did not observe a significant trajectory group by age interaction for inhibitory control (*F* = 1.13, *p* = .32), and post-hoc tests revealed no significant differences in predicted anti-saccade performance in among groups in early adolescence (Figure 7H top panel, all *p* > .05), or late adolescence (Figure 7H top panel, all *p* > .05). There were no significant differences in slopes between trajectory groups (Figure 7H bottom panel). These effects were similar across individual trial types (see Supplement). Together, these findings highlight statistically comparable developmental patterns of inhibitory control across trajectory groups.

#### Distinct Developmental Patterns of NAcc Tissue Iron across Trajectory Groups

Differences in nT2*w across trajectories, as well as interactions with age, were not present across most individual regions of interest (see Supplement), with the exception of the NAcc which showed the strongest effects and is therefore presented below and in Figure 7, but we additionally describe results for the whole basal ganglia and other regions in the Supplementary section. NAcc nT2*w differed significantly across substance use trajectory categories (Figure 7J; *F* = 3.07, *p* = .03), with higher overall NAcc nT2*w (indicating lower iron) in the *youth peak* group, followed by the *low* group, the *adolescent increasing* group, and finally, the *adult increasing* group. Independent GAMMs modelling the developmental time course of NAcc nT2*w in each trajectory group (Figure 7I – K) revealed significant age-related decreases across all groups (*p* < .001 in all 4 models; Figure 7I). However, trajectories differed in shape (Figure 7I) and rate of change (Figure 7Kiii). The *low* and *adult increasing* groups showed consistent, yet shallow, linear age-related decreases over time (indicating increases in tissue iron). In contrast, the *youth peak* and *adolescent increasing* groups showed curvilinear trajectories, with steeper, more rapid age-related decreases, particularly early on.

##### Variation in NAcc nT2*w across Age within Trajectory Groups

A significant trajectory group by age interaction emerged for NAcc tissue iron (*F* = 3.32, *p* = .007), and post-hoc tests revealed that in early adolescence (Figure 7L top panel), the *youth peak* group showed significantly higher nT2*w, indicating lower tissue iron, relative to all other trajectory groups (all *p*s < .05), however, this difference was diminished by young adulthood (all *ps* > .05; Figure 7L top panel). Additionally, the *youth peak* group showed significantly higher slopes in comparison to the *low* group (*p* = .006). Together, these findings highlight distinct developmental patterns of tissue iron across trajectory groups, and suggest that while the *youth peak* showed significantly lower tissue iron early in adolescence, they showed steeper increases over time, which may reflect relatively delayed NAcc tissue iron development and/or normalization of iron levels in this cohort.

All together, these results suggest variability in the magnitude and developmental trajectory of impulsivity, inhibitory control, and NAcc tissue iron across substance use trajectory subgroups. See Supplementary Table S8-S13 for full model statistics and post-hoc comparisons across trajectory groups, substances, UPPS subscales, anti-saccade trial types, and basal ganglia regions of interest.

## Discussion

This study examined how non-invasive neuroimaging markers of dopamine (DA) – related neurobiology and neurocognitive functioning shape patterns of substance use from adolescence into adulthood. By integrating MR-based assessments of basal ganglia tissue iron – an indirect correlate of presynaptic DA availability^61,62^ – with up to 9 repeated assessments per participant of impulsivity, inhibitory control, and substance use, we identified key biological and cognitive factors associated with the frequency and developmental time course of youth substance use. Across the large heterogeneous community and high-risk sample^73^ we found that increased impulsivity, reduced inhibitory control, and lower tissue iron were each independently associated with higher frequency and intensity of substance use. Using growth mixture modelling, we identified four distinct substance use trajectories, including no- or low use across the study period, youth peaks, linear increases beginning in adolescence, and finally, low use in adolescence but linear increases in adulthood. Notably, our findings highlight distinct developmental patterns in impulsivity, inhibitory control, and nucleus accumbens (NAcc) tissue iron concentration across distinct within-person trajectories of substance use; individuals who displayed *youth peak* and *adolescent increasing* patterns showed *increased* impulsivity, particularly early in adolescence, and *youth peak* patterns of substance use additionally showed *lower* levels of tissue iron, suggesting that this neurobehavioral pattern may reflect a predisposing phenotype for early engagement in substance use. These results suggest that biological changes within brain systems that are still undergoing development in adolescence, including DAergic and prefrontal-dependent cognitive systems, may contribute to youth patterns of substance use. To our knowledge, this is the first study to longitudinally link aspects of DA neurophysiology to substance use across adolescence *in vivo* in humans, prior to the escalation of substance use, allowing us to understand neurodevelopmental processes underlying youth propensity toward substance use.

### Age-Related Changes in Substance Use across the Full Sample and Within-Person Trajectories in Substance Use

Across the full sample, substance use exhibited non-linear trajectories across the study period, marked by increases in use in early adolescence that continued until young adulthood, followed by decreases that continued toward the upper limit of the study period (30 years of age). Despite variation in the magnitude of substance use across individual substances, all exhibited ages of initiation between 17 and 18 years of age, in addition to peak use between 21 and 22 years of age. Importantly, we show that while the magnitude of youth substance use varied across sociodemographic factors, including biological sex assigned at birth, the developmental timing, including age of initiation, early adolescent increase onsets, early adulthood peaks in substance use, and rates of change were relatively consistent across sociodemographic factors. This finding is in agreement with recent reports showing that the developmental timing of adolescent risk-taking is consistent across social, environmental, and psychological factors^78^.

Given that a primary objective of the current study was to identify neurodevelopmental differences that were predictive of variation in substance use trajectories (i.e., adolescent-specific vs long-term escalation into adulthood), we utilized an innovative growth mixture modelling approach that captured variation in within-person trajectories across timepoints and robustly identified subgroups based on patterns of substance use across development. These models indicated four distinct subtypes of trajectories (Figure 6). The *low* substance use group (slightly under 1/3 of the sample) was comprised of individuals with no- or low-levels of use across the entire study period and late age of initiation, with only minimal increases across adolescence, indicating that a significant proportion of youth are abstaining entirely or until adulthood, which echoes recent statistics showing that fewer youth are using alcohol^79^. The *youth peak* substance use group (approximately 1/4 of the sample) was comprised of individuals who showed the earliest age of initiation and highest rates of use overall, but with quadratic patterns across adolescence marked by rapid early increases, peaks in use in adolescence/young adulthood, notably, followed by decreases into adulthood. This group may reflect an instantiation of increased risk-taking propensity in adolescence, particularly given findings showing that more regular patterns of substance use are associated with higher levels of risk-taking in youth, both in this study and others^78,80^. The *adolescent increasing* group (approximately 17% of participants) showed early mean age of initiation, and the second highest rates of substance use overall, with rapid linear increases in use beginning at in mid-adolescence, potentially reflecting a group at increased risk for the development of long-term patterns of substance use into adulthood, though future work is needed to characterize long-term outcomes. Finally, the *adult increasing* group (approximately 1/4 of the sample) showed low levels of use during adolescence, and lower rates overall, with a later mean age of initiation, young adulthood peaks in use, and slower rates of change relative to *youth peak* and *adolescent increasing*, but continued increases into adulthood. We additionally show that population subgroups defined by sex, race/ethnicity, study site, and socioeconomic indicators were not disproportionately represented within any one trajectory group, further supporting the notion that the developmental timing, represented here as distinct trajectories within the population, was generalizable across subgroups.

### Impulsivity Relates to Developmental Trajectories of Substance Use

Across the full sample, impulsivity was heightened early in adolescence and linearly decreased with age. The magnitude of impulsivity varied depended on subscale, however, all subscales exhibited peak levels of impulsivity in late adolescence and decreased linearly until the mid-twenties. Notably, sensation seeking showed a distinct trajectory marked by a prolonged period of heightened sensation seeking across adolescence, with age-related decreases not beginning until approximately 18 years of age and continuing until the end of the study period, consistent with prior reports of adolescent sensation seeking^81,82^ and risk-taking more generally^17,78^. We additionally show that while the magnitude of impulsivity varied across sociodemographic factors, including biological sex assigned at birth (as has been shown previously in cross-sectional analyses^83,84^), and study site (potentially indexing geographical location), the developmental timing, including early adolescent decrease onsets, late adolescent peaks in impulsivity, and rates of change were relatively consistent across sociodemographic factors, in line with recent reports regarding the generalizability of adolescent risk-taking propensity across these factors^78^.

We show that heightened impulsivity was associated with increased substance use, both in terms of increased days of use, and further illustrated by higher levels of impulsivity in those who report greater substance use, with impulsivity scaling with frequency (e.g., ‘regular’ > ‘ever’ > ‘never’), and with the number of substances used (i.e., in those who report patterns of poly-substance use). These data provide compelling evidence that variation in trait-level impulsivity plays a critical role in the magnitude of substance use. We further showed that impulsivity varied as a function of the developmental time course of substance use; with significantly heightened impulsivity in the *youth peak* and *adolescent increasing* groups, both of which showed the highest rates and the earliest and most rapid changes in substance use, in comparison to the *low* group and the *adult increasing* group, both of which showed low use in adolescence. Both youth-onset trajectory groups showed early and persistent elevations in impulsivity, that importantly, preceded the mean age of initiation of substance use initiation. However, the *youth peak* group showed earlier and more substantial age-related decreases in impulsivity in comparison to the *adolescent increasing* group, evidenced by both higher slopes and higher rates of change that were comparable to the *low* substance use trajectory group, while the *adolescent increasing* group showed diminished age-related declines in impulsivity (Figure 7). These results may suggest that the increased patterns of substance use in the *youth peak* and *adolescent increasing* are supported by heightened impulsivity early on; but developmental decreases in substance use may have been supported by normative age-related decreases in impulsivity in the *youth peak* group. In contrast, the *adolescent increasing* group showed attenuated age-related decreases in impulsivity, potentially underlying prolonged vulnerability and elevated risk for sustained or persistent increases in substance use into adulthood. That impulsivity was elevated prior to substance use escalation suggests that it may be a predisposing factor for youth patterns of substance use, highlighting its role in shaping the trajectory of youth substance use.

### Inhibitory Control Relates to Developmental Trajectories of Substance Use

Across the full sample, inhibitory control robustly improved across adolescence, stabilizing in young adulthood, as has been observed in prior studies using the well-validated anti-saccade task^33,85–87^ and examining the maturational timing of executive function more broadly^87,88^. Performance was significantly enhanced during rewarded anti-saccade trials in comparison to neutral (non-rewarded) trials; however, in contrast to prior studies that show that this reward enhancement was more prominent early in adolescence^34,68,89^, we saw similar developmental trajectories across trial types. Across sociodemographic factors, inhibitory control performance varied only by study site, and the developmental timing was fairly consistent across the population subgroups for which we were well powered to detect significant maturational effects, including early adolescent increase onsets, late adolescent/early adulthood stabilization, late adolescent peaks in performance, and comparable rates of change.

We further show that across the full sample, lower inhibitory control was associated with increased levels of substance use, illustrated by decreased performance in those that report more regular patterns of use and with poly-substance use. This result is similar to findings with impulsivity, reflecting that decreased inhibitory control may also play an important role in substance use, which is complementary to prior reports in NCANDA showing decreased inhibitory control performance in youth at increased risk for substance use^14,15^. Additionally, associations did not differ across trial types, suggesting generalized inhibitory control involvement (i.e., rather than reward modulation of cognitive control). Although differences in inhibitory control were not significantly different across trajectory groups, qualitatively, the *youth peak* group showed a trend toward lower performance, in addition to diminished age-related improvements (Figure 7), potentially suggesting delayed or attenuated development of inhibitory control that could contribute to elevated levels of substance use in youth.

### Dopamine Neurobiology Relates to Developmental Trajectories of Substance Use

Consistent with prior studies using EPI-derived measures of tissue iron ^61,64,66,68,90,91^, nT2*w decreased linearly across each basal ganglia region of interest, reflecting age-related increases in tissue iron that parallel changes in animal models of DA availability during analogous developmental periods^30,92,93^. Importantly, we show that while the magnitude of basal ganglia nT2*w varied across sociodemographic factors, including biological sex assigned at birth, the developmental timing, was primarily consistent, including early adolescent increases in tissue iron, late adolescent/early adulthood peaks in tissue iron, and comparable rates of change. Interestingly, while analyses revealed increases in tissue iron across each region of interest, the caudate and the putamen exhibited the most rapid change between 25 and 27 years of age, but the NAcc exhibited a considerably earlier peak change at approximately 13 years, demonstrating relatively early, rapid, increases. Through the association between tissue iron and DA function^61,65,94,95^ within mesocorticolimbic systems, relatively earlier peak increases in the NAcc may be mechanistically related the outsized influence of limbic systems^96,97^ and peak reward reactivity^12,29,33,98,99^ in adolescence, contributing to unique behavioral patterns including increased risk-taking and youth patterns of substance use.

Across the full sample, lower basal ganglia tissue iron was associated with increased levels of substance use, both in terms of increased days of use, and further illustrated by lower levels of tissue iron in those who report more regular patterns of substance use. These data suggest that lower levels of DA neurophysiology are associated with increased substance use, potentially reflecting compensatory sensation seeking/exogenous DA stimulation^27,100^. We further showed that tissue iron varied as a function of the developmental time course of substance use, particularly within the NAcc; with the *youth peak* group, who had the earliest onset and earliest age-related increases in use, showing significantly lower tissue iron in comparison to all other trajectory groups, most prominently early in adolescence, that preceded the mean age of initiation. However notably, the *youth peak* group showed substantial age-related increases in tissue iron, evidenced by both high slopes and high rates of change, which may suggest normalization throughout adolescence as we observed no significant differences in NAcc nT2*w later in young adulthood (Figure 7). These findings suggest that low tissue iron, reflecting low DA levels, in early adolescence may predispose youth to earlier and more frequent substance use. However, the observed ‘catch-up’ in tissue iron maturation may support later reductions in use into adulthood. Of particular interest, the neurodevelopmental trajectory effects were especially pronounced in the NAcc relative to other basal ganglia regions, which aligns with the well-established role of the NAcc in reward-driven and risk-taking behaviors in adolescence^12,29,32,89,101^, as well as in substance use more broadly^18,19,71^. The prominence of NAcc-specific effects may reflect the unique contribution of its ongoing maturation in youth patterns of substance use, potentially by influencing motivational processes relevant to substance use initiation and early drug-seeking behavior^102^, in contrast to more dorsal striatal regions, which may play a greater role in the maintenance and habitual aspects of substance use^102^. The finding that *low* NAcc iron predicted increased use further aligns with neurodevelopmental models suggesting that low DA signaling might promote compensatory risk-taking and reward seeking behavior^27,100^, and with ‘inverted-U’ hypotheses of DA function^103^ whereby low levels of DA can increase risk behavior^104^. In light of the current findings, it is possible that low tissue iron may reflect low basal DA, or blunted endogenous DA, and may promote increased motivational salience of external rewards, including drugs, as a means of acutely boosting DA signaling^105–107^ and providing hedonic stimulation^108^ through reinforcing effects of substances, particularly in youth who show early substance use engagement.

### Strengths of the Current Study

A key strength of the current study is its longitudinal design, which enabled the assessment of within-person changes over time – essential for characterizing developmental trajectories and disentangling predisposing neurobiological factors from long-term consequences of substance use. Critically, enrolling primarily substance-naïve individuals (83% had limited or no history of substance use) allowed us to capture biological processes that may prospectively predict substance use frequency and trajectory, offering a window into early markers of vulnerability. Given the heterogeneity of substance use patterns among youth, this approach supports a nuanced understanding of how individual differences unfold across adolescence. Identifying early contributing neurobiological factors is an essential step toward elucidating the etiology of substance use disorders. Further, the broad age range captured in this study (12–30 years) enabled detailed mapping of long-term use patterns – allowing us to distinguish transient, potentially normative increases in use during adolescence and young adulthood (e.g., sensation seeking) from more persistent trajectories that extend into adulthood and may reflect heightened risk.

Although prior research has consistently shown that chronic substance use can lead to long-term alterations in DA function, it remains unclear how pre-existing differences in biological systems predispose individuals to substance use, or whether substance use itself disrupts the normative maturation of these systems. The current findings suggest that individual differences in impulsivity and DA-related brain function may serve as early markers that precede the substance use onset. Moreover, developmental trajectories of these systems may help distinguish youth-onset from adult-onset patterns of use. From a translational perspective, these results underscore the importance of early cognitive and DAergic system development in shaping adolescent behavior and highlight potential neurobiological risk markers for youth substance use initiation and engagement. Identifying such markers could inform the timing and targeting of early interventions – such as psychosocial or cognitive-behavioral strategies – designed to mitigate risk before problematic use emerges.

By examining substance use as a composite construct—including nicotine, cannabis, and alcohol—in addition to isolating individual substances, the current study was able to capture adolescents’ general propensity toward substance use and identify underlying DAergic and neurocognitive patterns linked to broader risk-taking phenotypes. This approach offers an integrative view of substance use vulnerability during a critical developmental window. Notably, the consistency of developmental findings across sociodemographic subgroups—defined by biological sex assigned at birth, race/ethnicity, socioeconomic indicators, and study site—suggests that both the developmental timing of risk and the biological mechanisms underlying substance use propensity are largely conserved across diverse populations. Although prior research has documented variability in the magnitude of substance use and neurocognitive functioning across demographic groups, far less attention has been paid to the developmental timing of these associations. This distinction is crucial for advancing mechanistic models of the etiology and pathogenesis of substance use disorders, which often emerge during adolescence, and for informing the design of developmentally sensitive interventions. Furthermore, while previous studies have reported age-related changes in DAergic and neurocognitive metrics, the precise timing, magnitude, and periods of maturation have remained poorly defined. Leveraging the large and deeply phenotyped sample from NCANDA-A (N = 831; total visits = 6,268), and applying nonlinear and growth mixture modeling approaches, we were able to delineate specific developmental windows during which substance use behaviors and their associated biological underpinnings undergo significant change.

### Opportunities for Ongoing Investigations and Methodological Limitations

Several key limitations should be acknowledged, each of which highlights opportunities for important future research. First, we did not assess whether trajectory group membership was associated with differential long-term clinical outcomes (i.e., following long-term use). Future work should explore how these trajectories predict distinct adult outcomes, including neuropsychiatric diagnoses and substance use disorders. Second, we did not observe significant differences in impulsivity, inhibitory control, or tissue iron among individuals in the *adult increasing group,* with developmental trajectories that were comparable to those of the *low*-use group. This pattern suggests that adult-onset increases in use may be influenced by other factors not captured in the current analyses – such as contextual or environmental stressors. Future research should aim to identify and characterize factors that contribute to adult-onset patterns of substance use. Third, future research should extend beyond the neurocognitive and biological metrics examined in this study to investigate the influence of social factors known to shape youth substance use patterns. These include peer norms, peer and parental substance use attitudes, and societal attitudes^1,109^ toward substance use, and youths’ own motivations – such as using substances to cope with stress or alleviate boredom^110–112^ – as potential drivers or distinguishing factors in substance use trajectories. Finally, changing policies and legislation, such as the introduction of smoke-free laws for tobacco use^113^ and, conversely, the legalization of cannabis in various countries and states^1^, may influence societal attitudes toward substance use, and, consequently, youth engagement with these substances. In addition, broader social and environmental factors, including societal disruptions like social distancing during the COVID-19 pandemic^1^, are important to consider when examining shifts in youth substance use patterns over time. Ongoing surveillance efforts, such as the *Monitoring the Future* study^1^, along with future research that integrates these contextual factors, will be critical in advancing our understanding of youth substance use risk and propensity.

Understanding the factors that contribute to variation in iron content leading into adolescence, for example, lower tissue iron levels that in turn increase susceptibility to maladaptive behaviors such as substance use, remains an open question and represents a critical direction for future work. Early life factors that influence DA system development may shape individual differences in reward sensitivity and impulse control, thereby increasing vulnerability to later substance use. Genetic and epigenetic variation in DA-related genes^114,115^, prenatal exposures (i.e., maternal stress and inflammation^116,117^), and early environmental factors (i.e., adversity^118,119^, nutritional deficiencies, and toxin exposure^120,121^) – have all been linked to disruptions in the development of DA systems and frontostriatal circuitry. Thus, early alterations in DAergic functioning may represent a mechanistic vulnerability contributing to the developmental onset of substance use.

Methodologically, while T2*-based indices provide a valuable, non-invasive proxy for tissue iron – and by extension, DAergic physiology – it remains an indirect measure. Future research would benefit from incorporating more direct, yet still non-invasive, methods to further elucidate the role of dopamine in substance use trajectories. Additionally, because the T2* signal is influenced by myelin content^122,123^, our analyses focused on the basal ganglia, where iron concentrations are highest and most reliably drive T2* signal variation^48,49,124^. This targeted approach enhances interpretability but also opens important avenues for future work to explore DA-related processes in other mesocorticolimbic regions – such as the PFC – and across broader distributed circuits implicated in substance use risk and regulation.

Finally, the inhibitory control task was administered at only two research sites, which provided sufficient power to detect associations and characterize periods of significant change across the full sample. However, statistical power was substantially reduced when subdividing participants into subgroups, limiting the ability to detect windows of significant change within smaller groups. While these considerations should inform the interpretation of subgroup comparisons, they also highlight an opportunity for future studies with larger, more diverse samples and repeated measurement designs to better characterize developmental patterns of inhibitory control across distinct risk and trajectory profiles.

### Conclusions

Together, these findings suggest that high impulsivity, low inhibitory control, and low DA may contribute to youth substance use behavior, potentially reflecting increased risk-taking behavior during adolescence as reward and cognitive systems continue to undergo maturation and adult trajectories are being established. These neurobehavioral differences may emerge early in adolescence, prior to substance use initiation, and may normalize over time in some individuals, potentially through age-related maturation of DA systems, normative decreases in impulsivity, and improvements in cognitive control. In contrast, attenuation of age-related changes that lead to persistent impulsivity may contribute to prolonged substance use in some individuals. This study contributes to a growing body of work that suggests that developmental trajectories of mesocorticolimbic DA and cognitive control systems are central to adolescent substance use vulnerability. Early identification of these mechanisms can improve prevention efforts and better target interventions that align with neurodevelopmentally sensitive periods. Ultimately, understanding how maturing DA and neurocognitive systems shape health-risk behaviors can inform clinical practice, public health strategies, and basic neuroscience.

## Materials and Methods

### Participants

Data for this project were provided from participants recruited by the National Consortium on Alcohol & Neurodevelopment in Adolescence and Adulthood (NCANDA-A) study^73^ at five NCANDA-A sites across the United States: University of Pittsburgh Medical Center (UPMC), SRI International (SRI), Duke University (Duke), Oregon Health & Science University (OHSU), and University of California San Diego (UCSD). NCANDA-A is a longitudinal, prospective study of healthy 12-22 year olds^73^, aimed at investigating whether and how normative neurodevelopmental trajectories of brain structure and function contribute to and/or are impacted by substance use. At each visit, participants completed a neuropsychological battery, 3T magnetic resonance imaging (MRI) neuroimaging session, and a comprehensive assessment of substance use, psychiatric symptoms, and diagnoses as well as functioning in major life domains^73^. Eight hundred and seven adolescents and young adults (400 female; age range = 12 – 22 years old at baseline) participated in an accelerated longitudinal study (total participant visits = 6164; median number of visits per-participant = 5; range of visits per-participant = 1-9; age range = 12 – 30 years old across all included longitudinal visits; see Figure S1 for participant age by visit structure). Participants were screened for study exclusion criteria including MRI contraindications (e.g., claustrophobia, non-removable metal in the body), head injury with significant loss of consciousness, psychiatric disorders that might influence study completion (e.g., psychosis), and psychiatric medication^73^. Given that a central goal of NCANDA-A was to examine the transition to significant substance use during adolescence into adulthood, approximately 50% of participants were recruited based on subclinical risk factors for alcohol use disorder, however, included participants had limited or no history of alcohol or other drug use at baseline as per National Institute on Alcohol Abuse and Alcoholism (NIAAA) guidelines for risky drinking^73^. To optimize generalizability and representation (see^78,125,126^ for discussion) and due to studies demonstrating that the developmental timing of adolescent risk-taking behavior is consistent across sociodemographic factors^78^, no participant-level demographic exclusion criteria were applied to the analysis sample, and we additionally show the potential effect of such factors in a series of sensitivity analyses. Participants over 18 provided informed consent, while participants younger than 18 provided written assent and parental consent. All experimental procedures obtained full Institutional Review Board (IRB) approval at each site, and complied with the Code of Ethics of the World Medical Association (Declaration of Helsinki, 1964)^127^. Each site recruited representative community samples with respect to local racial/ethnic distributions, ensuring balance for equal distribution of sex by age group. Participant demographic information is provided in Table 1 (but see^73^ for detailed sampling strategy, recruitment information, and assessment protocol).

**Table 1.** Participant characteristics at baseline. Coding for race/ethnicity, household income, guardian education, sex, age at baseline, study site, and MRI scan manufacturer. Guardian education reflects average across guardians. All variables were assessed via self-report. See ‘Participants’ for information on recruitment. Non-Hispanic (NH). African American (AA). American Indian and Alaska Native (AIAN). Native Hawaiian and Pacific Islander (NHPI).

| Characteristic | Category | % | N |
| --- | --- | --- | --- |
| Race/Ethnicity | Asian, AIAN, NHPI | 8.18 | 66 |
|  | Hispanic | 11.90 | 96 |
|  | NH Black/AA | 12.02 | 97 |
|  | NH White | 62.95 | 508 |
|  | Unspecified | 4.96 | 40 |
| Household Income | <25k | 5.45 | 44 |
|  | 25k-49k | 10.29 | 83 |
|  | 50k-74k | 11.40 | 92 |
|  | 75k-99k | 12.14 | 98 |
|  | 100k-199k | 41.39 | 334 |
|  | >200k | 19.33 | 156 |
| Guardian Education (Average) | < Grade 9 | .87 | 7 |
|  | High school | 7.43 | 60 |
|  | College | 49.44 | 399 |
|  | Graduate School | 42.26 | 341 |
| Sex | F | 49.63 | 400 |
|  | M | 50.37 | 407 |
| Age at Baseline | 12-14 | 31.92 | 258 |
|  | 15-17 | 38.40 | 310 |
|  | 18-22 | 29.68 | 239 |
| Site | A | 14.84 | 120 |
|  | B | 20.07 | 162 |
|  | C | 20.57 | 166 |
|  | D | 18.70 | 151 |
|  | E | 25.81 | 208 |
| Scanner | GE | 66.46 | 536 |
|  | Siemens | 33.54 | 271 |

#### Substance Use Assessments

Participants completed the self-report Customary Drinking and Drug-use Record (CDDR) at each visit, which assesses the frequency (never, ever, regular (weekly) use) and quantity of recent (i.e., number of days in the past 30 days/past year) and lifetime alcohol and substance use, including nicotine, cannabis, and other substances (e.g., hallucinogens, opiates^73,128^). Additionally, binge drinking was assessed based on the NIAAA guidelines of consuming greater than four or five drinks for females and males, respectively, within an occasion. We additionally examined poly-substance use patterns by computing whether participants reported using one substance in a given month (single-use), two substances (co-use), or greater than two substances (poly-use). Participants further report on the age at first use and age of initiation of regular (weekly) substance use. Data (frequency and quantity) were collected separately for each substance type at each visit. Here, across each substance, frequency was assessed as the number of days of use in the past 30 days, focusing solely on alcohol use, nicotine, cannabis, and binge drinking.

#### Impulsivity Measures

##### UPPS

To assess constructs of impulsivity, the self-report Short UPPS-P Impulsive Behavior Scale (SUPPS-P) was administered at each session^74–76^, which includes subscales assessing five distinct dimensions of impulsivity, including sensation seeking, negative urgency, positive urgency, perseverance (lack of), and premeditation (lack of). Each subscale ranged from 1-4, with 4 being the maximum score. We also constructed an average score by calculating the mean across all subscales, reflecting impulsivity independent of subscale.

#### Executive Function Measures

##### Rewarded Anti-saccade Task Design and Timing

Two hundred and seventy-six participants (150 female, 12 – 24 years old, 1-5 visits, 682 total sessions) at two participating sites (Duke & UPMC) additionally completed a rewarded anti-saccade task (Figure 3A), as has been previously been described in detail^15,33,77^. The full protocol included a total of 56 full incentivized (‘reward’) and 56 full non-incentivized (‘neutral’) anti-saccade trials, completed across 4 fMRI runs, each consisting of 28 trials for a total of 112 trials. See Supplemental Methods for details regarding stimulus presentation, eye tracking recording and apparatus. On each full trial, participants were presented with one of two cues (cue epoch) during which a fixation point was surrounded by either a ring of dollar signs (“$”; each subtending approximately 1° of visual angle), indicating the chance to earn reward for correctly performing the forthcoming trial (‘reward’ trials), or an equivalently sized, isoluminant ring of pound signs (“#”), indicating that there was no money at stake for that trial, regardless of performance (‘neutral’ trials) (1.5 s, pseudorandomly interleaved conditions). Following the disappearance of the incentive ring, the central fixation cue turned red to indicate the response preparation period (1.5 s). Finally, a peripheral cue (yellow ellipse) appeared in an unknown location along the horizontal midline (75 ms; ± 3°, 6°, or 9° visual angle to the left or right, relative to the central fixation), and participants were instructed to direct their gaze to the mirror location of the stimulus, completing an anti-saccade (response epoch; 1.5 s). Each trial was followed by a fixation period with a variable intertrial interval (ITI; jittered between 1.5, 3, or 4 s). An additional 24 (12 rewarded, 12 neutral) partial trials that presented either the incentive cue alone (6 of each reward and neutral trial types) or the incentive cue and preparation epoch (6 of each reward and neutral trial types), but not the response epoch, were included as ‘catch trials’ in order to estimate the hemodynamic response for each trial epoch (incentive cue and response preparation epochs, respectively), however, these trials were not included in our analyses. Each run was approximately 5 min 9 s in duration and consisted of 14 neutral trials and 14 reward trials. Four runs were completed in total, for a total of 56 full reward and 56 full neutral trials. In order to prevent participants computing a running total of rewards and/or engaging working memory processes, they were not explicitly told how much money could be earned on a given ‘reward’ trial (‘$’) and were not given explicit ongoing feedback throughout the task. However, they were informed prior to the task that they could win up to an additional $10 contingent on their performance and that no penalty would be accrued for incorrect performance (i.e., loss of monetary reward).

The behavioral indicator of interest was accuracy (the percentage of correct anti-saccade responses) which was calculated as the number of correct trials divided by the number of scorable (tracked) trials and was calculated separately for reward and neutral trials. Sessions with fewer than 25 viable trials total, as well as correct response rates lower than 40%, were excluded from analysis, resulting in a final sample of 257 12- to 24-year-old participants (134 female), for a total of 586 sessions.

#### MRI Data Processing

##### Structural Imaging Acquisition & Preprocessing

Multisite protocols ensured that comparable 3T MR data were acquired across the 5 imaging sites (methods have been reported in detail^15,129^). Two sites used Siemens TIM-Trio scanners (UPMC and OHSU) and three sites used GE Discovery MR750 scanners (UCSD, SRI International, & Duke); however, all participants underwent the similar scan protocols at each timepoint. Structural (T1-weighted) images were acquired on Siemens scanners using a magnetization-prepared rapid gradient-echo (MPRAGE) pulse sequence (TR, 6 ms; TE, 3 ms; flip angle, 9°; matrix, 256x 256; FOV, 24 cm; 1.2 x .94 x .94 mm voxel size, 160 slices), and on GE scanners using an Inversion Recovery-Spoiled Gradient Recalled echo (IR-SPGR) sequence (TR, 6 ms; TE, 2 ms; flip angle, 11°; matrix, 256 x 256; FOV, 24cm; 1.2 x .94 x .94 mm voxel size, 146 slices). Structural data were preprocessed to extract the brain from the skull, and warped to the MNI standard brain using both linear (FLIRT) and non-linear (FNIRT) transformations.

#### Acquisition & Preprocessing of T2*-Weighted Indices Reflecting Tissue Iron Properties

T2*-weighted MR data were acquired over a 10-minute fixation resting state scan using 3D blood oxygen level dependent (BOLD) echoplanar sequence (TR, 2,200 ms; TE, 30 ms; flip angle, 79°; 3.75 x 3.75 x 5 mm voxel size; 32 slices; 275 volumes) maximally covering the cortex and basal ganglia. T2* data were minimally preprocessed as required for this measure, and included 4D slice-timing and head motion correction, skull stripping, co-registration to the structural image, and nonlinear warping to MNI space ^65,68^.

##### Time-Averaging and Normalization of T2 Data (nT2*w)

T2*-weighted BOLD images were preprocessed as in ^65,66,68,130,131^, including the following steps: Each volume was first normalized to the whole-brain mean (z-score normalization) using a coverage map created from all inputs for each participant including only non-zero values. This normalized signal was then aggregated voxel-wise across all volumes in the resting state scan using the median, which reduces the effect of outlier volumes resulting in one normalized T2*-weighted image for each participant (nT2*w). Volumes containing frame-wise displacement (FD) > 0.3mm were identified as high motion timepoints and were excluded from analyses ^132^. This procedure ensures enhanced SNR by averaging across multiple timepoints ^65^, and the normalization provides the T2* decay in the basal ganglia relative to the whole brain, which allows for comparison of nT2*w values across participants. Critically, as shorter T2* relaxation times indicate stronger magnetic field inhomogeneities induced by iron, nT2*w is inversely related to iron, with lower values indicating *higher* iron content, and higher values indicating *lower* iron content.

##### Regions of Interest

Given evidence from prior studies showing mixed results in terms of specificity across basal ganglia ROIs in modulation of cognitive and reward function during adolescence ^68^, we extracted nT2*w values across all basal ganglia regions of interest (Figure 4A) including the globus pallidus (GP), nucleus accumbens (NAcc), putamen, and caudate nucleus, and additionally computed one per-participant estimate of nT2*w that reflected the mean across all voxels in the basal ganglia across both hemispheres (left and right combined) defined using the Harvard-Oxford subcortical atlas ^133^. Consistent with prior work ^68,130^, basal ganglia estimates of nT2* were longitudinally stable across sessions, evidenced by high intra class correlation coefficient (ICCs, Psych Package, version 2.1.6, ^134^) of nT2*w values across visits (.89).

nT2*w indices of basal ganglia physiology were available in 799 12- to 29-year-old participants (395 female) for a total of 4363 sessions.

##### Harmonization of nT2*w across Scanners

nT2*w estimates from BOLD data approximates the T2* decay time based on the mean amplitude of BOLD signal normalized to the whole-brain amplitude, which could result in variation in absolute values for nT2*w across different scanners, software versions, and scan parameters. Given the multi-site nature of NCANDA-A, as well as variation in scanner manufacturer (GE, Siemens) across sites, and changes to scanner model version within sites over time, we harmonized nT2*w data using NeuroCombat (*neuroCombat* package version 1.0.13 in R^135^). We found that harmonizing based on scanner model version ameliorated substantial variability in nT2*w values relative to harmonizing based on site or scanner manufacturer alone, and additionally improved the consistency of age-effects and minimized between-site variation in nT2*w (see Supplement for additional details). Thus, all nT2*w values reflect adjusted estimates based on harmonization procedures.

### Data Analysis

#### Characterization of Substance Use Patterns

Participants’ responses from the CDDR were assessed at each visit in the following ways for past 30-day substance use: (1) Number of days of use within the past 30 days [alcohol, cannabis, nicotine], number of binge drinking days within the past 30 days (based on NIAAA guidelines, see above); (2) Responses were additionally grouped into the following frequency categorizations for each substance: never (reporting never having used at that timepoint), ever (reporting having ever used), and regular (reporting having used at least once/week); (3) Responses were grouped into the following use-type categorizations for each substance: no-use (reporting never having used that substance), co-use (reporting using that substance plus another substance in the past 30 days), and poly-use (reporting using that substance plus 2 or more other substances in the past 30 days). The frequency and use-type categorizations allowed us to ascertain whether effects occurred in a dose-dependent manner that scales with the number of substances that were used concurrently.

Primary analyses investigated substance use patterns overall via a probabilistic score that estimated the number of days (out of 30) that individuals used at least one substance (see Supplement). Follow-up sensitivity analyses investigated effects across each substance specifically, including alcohol, binge drinking, cannabis, and nicotine in order to ascertain whether risk-makers were associated with general patterns of use or whether they reflected individualized risk for different substances (that might have different etiological contributors and developmental time courses).

#### General Statistical Approach and Theoretical Models

All analyses were conducted in R via RStudio version 4.3.1 (R Core Team (2023), https://www.r-project.org). Two streams of analyses were implemented, the first to describe age-related changes in substance use, impulsivity, inhibitory control, and basal ganglia tissue iron, and to examine effects of the latter three metrics on concurrent substance use (termed ‘between-person analyses’ below), and the second to predict whether variation in these factors were predictive of distinct trajectories in substance use across the study period (termed ‘within-person analyses’ below).

##### Age Analyses across the Full Sample

We applied sequential models to first characterize age-related changes in substance use across all substances, as well impulsivity (UPPS), inhibitory control (anti-saccade), and basal ganglia tissue iron (nT2*w; *Model 1*), across the full sample. We next tested for associations between substance use and 1.Impulsivity (UPPS), 2. Inhibitory control (anti-saccade performance), and 3. Tissue iron (nT2*w; *Models 2 & 3),* While primary models were conducted across a composite measure of substance use (see Methods), a series of sensitivity analyses repeated these analyses separately across substances (see Supplement).

All statistical analyses utilized generalized additive mixed models (GAMMs; *mgcv* package in R ^136^, version 3.5.2), including random intercepts and random slopes estimated for each participant to account for the longitudinal aspect of the dataset. All continuous variables were modelled as smooth terms (age, impulsivity, anti-saccade, and tissue iron), and sociodemographic variables including biological sex assigned at birth, family income, and study site were coded as categorical variables (factors) and were modelled as parametric terms.

We characterized age-related changes across all measures (‘outcome variable’, *Model 1*) using GAMMs, which were chosen to characterize non-linear developmental trajectories given that non-linear fits have been shown to provide a better approximation of developmental changes throughout adolescence ^86,88,137–139^. Penalized natural cubic regression splines were implemented via the *mgcv* package to characterize linear and/or non-linear age trajectories ^140,141^, and a maximum basis complexity (*k*) of 4 was chosen consistent with prior studies utilizing GAMMs in adolescent populations ^64,142–146^ based on theoretical expectations that there are unlikely to be more than three resolvable inflection points (e.g., childhood, adolescence, adulthood stages), and to prevent overfitting the age smooth function. We used Restricted Maximum Likelihood (REML) to fit our GAMMs with unbiased estimates of the variance components in the context of random effects^147^. To assess age-ranges of significant change, we calculated the average first derivative of the age spline from the GAMM using finite differences (as were conducted in^64,148^), and 95% confidence intervals of the derivatives were generated (*gratia* package in R ^149^). Confidence intervals of the derivative that did not include zero indicated intervals of significant change. Average derivative was computed as an estimate of the average rate of change per year, across the entire age range. To additionally quantify developmental trends in the data, we computed the mean per-participant value (mean), as well as the peak value (maximum across all timepoints) across all metrics [substance use, impulsivity, inhibitory control performance, tissue iron], and we took the age (years) of this peak value to estimate the age of peak [substance use, impulsivity, inhibitory control performance, tissue iron].

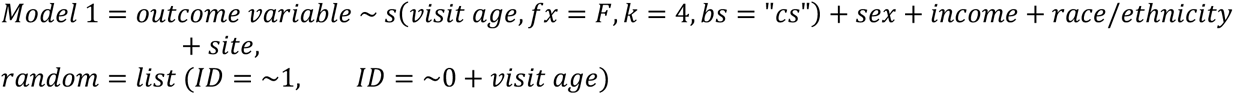

##### Participant-Level Sociodemographic Sensitivity Analyses

Sensitivity analyses explored how age-related trajectories changed as a function of 18 subgroups defined via levels of four variables (sex, race/ethnicity, study site, and socioeconomic indicators (family income), see Table 1) that have previously been shown to be associated with adolescent risk-taking propensity^78^. Independent models assessing developmental trajectories, mean, peak, and age of peak (as above) across each variable of interest [substance use, impulsivity, inhibitory control, and tissue iron] were conducted separately across these 18 subgroups to determine generalizability of developmental models.

##### Characterizing Relationships between Substance Use and Impulsivity, Inhibitory Control, and Basal Ganglia Tissue Iron

To ascertain whether risk-markers were associated with qualitative shifts in use patterns, substance use was grouped into categories at each visit reflecting substance use frequency (i.e., never, ever, regular; see above) and use-type (i.e., no-use, single-use, co-use, poly-use indicating higher use across multiple substances). Models then interrogated whether 1.Impulsivity (UPPS), 2. Inhibitory control (Anti-saccade performance), and 3. Tissue iron (nT2*w) differed within each category (*outcome variable; Model 2)*, independent of age (controlling for a smoothed term of age).

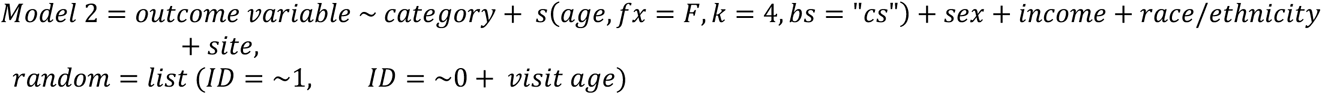

We then investigated the relationship between continuous measures of substance use (number of days in the past 30 days) and 1.Impulsivity (UPPS), 2. Inhibitory control (Anti-saccade performance), and 3. Basal ganglia tissue iron (nT2*w). Substance use was the dependent variable, and all independent variables were modelled as smoothed terms. Importantly, we covaried for a smoothed term of age in the model as a covariate to account for non-linear effects of age (*Model 3*). Continuous measures of substance use were log transformed for these analyses. Penalized thin plate regression splines were implemented via the *mgcv* package and a maximum basis complexity (*k*) of 3 was chosen as we did not expect these associations to largely deviate from linearity.

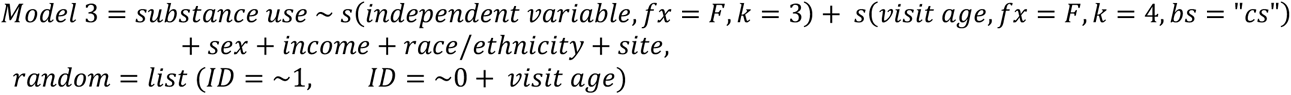

To ascertain specificity of impulsivity (UPPS) subscales, inhibitory control task conditions (anti-saccade, reward vs neutral trials), and basal ganglia regions of interest on substance use, initial models tested an interaction effect across each level of these measures. These models revealed that the interaction term was not significant for the effects of UPPS subscale on substance use (*F* = .40, *p* = .85), indicating uniform effects of each subscale on substance use, and we therefore utilized UPPS total scores for all primary analyses. The interaction model was also not significant for the effects of anti-saccade trial condition on substance use (*F* = .19, *p* = .83), indicating similar effects of reward and neutral anti-saccade performance on substance use, and we therefore used performance combined across conditions for all primary analyses. Finally, the interaction model was not significant for the effects of tissue iron within each basal ganglia region of interest on substance use (*F* = 1.99, *p* = .09), indicating lack of regional specificity in associations with substance use, and we therefore utilized an estimate across the whole basal ganglia for all primary analyses. However, we present results across each subscale of the UPPS, given prior results showing effects primarily for specific subscales (i.e., negative urgency^15,150^), each trial condition of the inhibitory control task, given prior results showing inconsistent results with regard to specificity across trial types (i.e., rewarded condition^77,151,152^), and each region within the basal ganglia, given prior results showing effects of specific regions on adolescent behavior and cognitive function (i.e., the nucleus accumbens^17,64^) in the supplemental results section.

##### Within-Person Trajectory Classification Analyses

We characterized *trajectories* in substance use by applying growth mixture models (i.e., Latent Class Mixed Model; *LCMM* package in R^153,154^, version 2.1.0) that captured variation in within-person trajectories (patterns of change) across timepoints and classify individuals based on patterns of substance use across development^155^ (*Model 4)*. Iterative class models were performed that tested for 1 – 4 separate classes of substance use trajectories, and the best fitting model was chosen based on the minimum Bayesian information criteria (BIC), Akaike information criteria (AIC), entropy, and integrated completed likelihood (ICL). Random intercepts were modelled per class, allowing each class to follow its own developmental trajectory, with participant and age as random effects. The same covariate structure as above was used, with the addition of baseline age as a between-participant variable to account for the longitudinal cohort design and to account for between- and within-participant effects, and modeling center age and a quadratic trend for center age (*Model 4)*. Given that fitting mixture models can be sensitive to initial values, we utilized a gridsearch function to improve and stabilize estimations of the models, and to improve the chance of finding the global maximum likelihood^153,154^. The model was estimated 100 times with different random starting values, with each repetition running up to 30 iterations. We limited our models to 1 – 4 classes based on theoretical expectations of developmental heterogeneity^156^, recommendations from prior mixture modeling research, and to ensure adequate class sizes, interpretability, and model parsimony^157,158^.

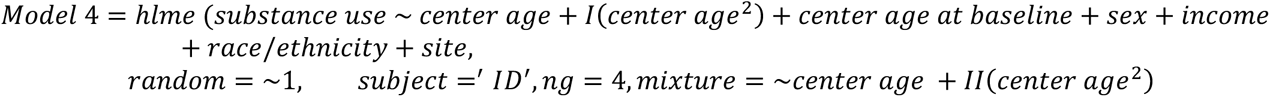

##### Participant-Level Sociodemographic Sensitivity Analyses

In order to confirm that sociodemographic factors were not disproportionately represented within each trajectory group, sensitivity analyses explored how trajectory group membership varied as a function of the 18 sociodemographic subgroups defined above.

##### Characterizing whether Variation in Impulsivity, Inhibitory Control, and Tissue Iron contributed to Within-Person Trajectories in Substance Use

Utilizing GAMMs (as above), we then tested for differences in 1.Impulsivity (UPPS), 2. Inhibitory control (anti-saccade performance), and 3. Basal ganglia tissue iron (nT2*w) across trajectory groups to assess whether these metrics differed within each trajectory group (*outcome variable; Model 2)*.

##### Characterizing Significant Trajectory group by Age Interactions on Substance use

Finally, we tested for interactions with age and trajectory group on each variable that might suggest that the effect of these factors varied across development (i.e., to ascertain whether trajectory groups had differences in DA early in development prior to the onset of substance use, or in young adulthood following substance use exposure; *Model 5*). These models used the same covariate structures as above.

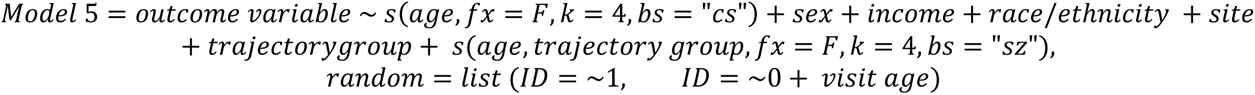

We characterized the nature of significant age interactions within each trajectory group using the *ggpredict* function in R (*ggeffects,* version 1.1.5^159^). To characterize significant interactions, we specified fixed values for the age that approximated the range of our data (age 14 for early adolescence; age 25 for young adulthood, with the exception of interactions involving the anti-saccade task in which we tested age 23 for young adulthood due to the restricted age-range). We then plotted the predicted relationship between age and dependent variable for each trajectory group and tested for differences at each age using the *hypothesis test* function (*ggeffects)*. Furthermore, we assessed differences between slopes across each measure within trajectories. Finally, to further characterize whether differences in maturational timing of impulsivity, inhibitory control, and tissue iron contributed to substance use trajectories, we repeated the growth rate modelling above within each trajectory group (*Model 1*).

## Data availability

Access to the full NCANDA dataset is available via the NIAAA Data Archive (collection 4513, https://nda.nih.gov/edit_collection.html?id=4513).

## Code availability

All original code generated for this study is available on github at https://github.com/LabNeuroCogDevel/NCANDA-Substance-Use-Trajectories-DA-Neurocog. In addition, code for computing the T2*-based estimates of basal ganglia tissue iron is available on github through https://zenodo.org/records/8302458. Original study code integrated publicly available software, including AFNI (available at https://afni.nimh.nih.gov/pub/dist/doc/htmldoc/index.html; version 23.1.10). All statistical analyses were conducted in R version 4.1 (https://cran.r-project.org/bin/macosx/).

## Acknowledgements

We thank the participants, the NCANDA research assistants, and the Laboratory for Neurocognitive Development (LNCD) for helpful comments on an earlier draft.

## Author Contributions

A.C.P, F.J.C., S.F.T., K.N., D.G., D.C., and B.L. designed the study; D.F., D.G., and D.C. collected the data; A.C.P., A.O, D.J.P., F.J.C., B.T.C, W.F., and W.T. analyzed the data; A.C.P. drafted the manuscript with input from A.O., D.J.P., F.J.C., B.T.C., S.F.T., D.G., D.C., and B.L.

