## Supplemental data tables for "Developmental variation in dopamine neurobiology, neurocognitive functioning, and impulsivity shape substance use trajectories in youth"

**Supplementary Methods**

***Eye Tracking During the Anti-Saccade Task***

*Stimulus Presentation and Eye Tracking Recording and Apparatus during the Anti-Saccade Task*

Monocular eye position data was recorded using a long-range optics eye-tracking system (Applied Science Laboratories, Bedford, MA; see^1^ for details). Eye-position was monitored via corneal reflection. Online video monitoring was performed to ensure task compliance. Eye position was calibrated using a 9-point calibration routine prior to each session and between runs, if necessary. All visual stimuli were presented on a flatscreen behind the MRI bore with E-Prime software^1,2^ and were made visible to the participant with a mirror mounted on the head coil.

A correct response was defined as the first eye movement during the saccade response epoch with a velocity greater than or equal to 30º/s that was made toward the mirror location of the peripheral cue and extended beyond a 2.5º visual angle central fixation zone. In contrast, an incorrect response was defined as the first eye movement during the saccade response epoch directed toward the peripheral cue and extending beyond at least 2.5º visual angle central fixation zone. Trials in which no saccade occurred were defined as ‘non-response’ trials and were excluded from the analysis. Trials in which the eye-tracker lost the participants’ eye were defined as dropped trials and were excluded from the analysis.

1.	Tervo-Clemmens, B. *et al.* Neural Correlates of Rewarded Response Inhibition in Youth at Risk for Problematic Alcohol Use. *Front. Behav. Neurosci.* **11**, (2017).

2.	Schneider, W., Eschman, A. & Zuccolotto, A. *E-Prime: User’s Guide*. (Psychology Software Incorporated, 2002).

**Supplementary Figures**

**Supplementary Figure S1.** Age Range and study Design of NCANDA Dataset

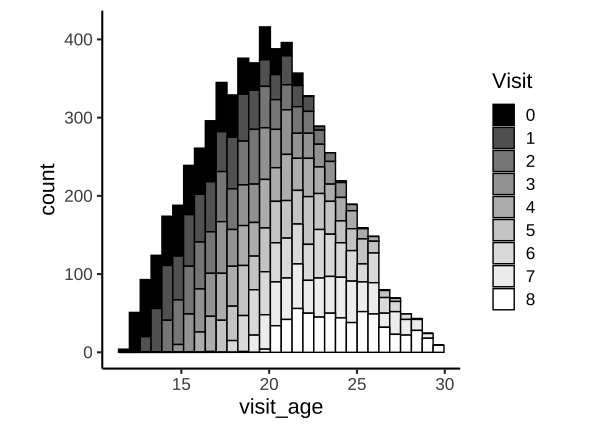

Histogram of participant age and visit number. NCANDA-A used an accelerated longitudinal design with up to 9 visits per-participant (807 participants; 6164 sessions total following exclusions), spanning the full adolescent period and relevant transitional periods from adolescence to young adulthood (total age-range: 12-30 years old).

**Supplementary Tables**

**Supplementary Table S1. Full Sample: Statistics Across Individual Substances, Subscales, Trial Types, and Regions of Interest**

| **Substance Use: Statistics Across Individual Substances** | | | | | | | |
| --- | --- | --- | --- | --- | --- | --- | --- |
| **Substance** | **Population** | **Mean n Days** | **SE n Days** | **Peak Use (days)** | **SE Peak Use (days)** | **Mean Age of Onset (years)** | **Age of peak use (years)** |
| All | Full Sample | 5.69 | .11 | 15.75 | .14 | 16.66 | 21.94 |
| Alcohol | Full Sample | 2.54 | .06 | 8.74 | .09 | 17.01 | 22.11 |
| Binge | Full Sample | .63 | .02 | 3.87 | .07 | 18.20 | 21.25 |
| Cannabis | Full Sample | 2.73 | .09 | 13.35 | .20 | 17.16 | 21.39 |
| Nicotine | Full Sample | 1.57 | .07 | 13.59 | .26 | 17.73 | 21.03 |
| **Impulsivity (UPPS): Statistics Across Individual Subscales** | | | | | | | |
| **Subscale** | **Population** | **Mean Score** | **SE Score** | | **Peak Score** | **SE peak Score** | **Age of Peak Score (years)** |
| Mean across all subscales | Full Sample | 1.92 | 0.01 | | 2.22 | 0.00 | 18.21 |
| Negative Urgency | Full Sample | 1.86 | 0.01 | | 2.48 | 0.01 | 18.25 |
| Perseverance | Full Sample | 1.76 | 0.01 | | 2.18 | 0.01 | 18.21 |
| Positive Urgency | Full Sample | 1.64 | 0.01 | | 2.18 | 0.01 | 17.84 |
| Premeditation | Full Sample | 1.63 | 0.01 | | 2.08 | 0.01 | 18.17 |
| Sensation Seeking | Full Sample | 2.71 | 0.01 | | 3.20 | 0.01 | 18.07 |
| **Inhibitory Control (Anti-Saccade): Statistics Across Individual Trial Types** | | | | | | | |
| **Trial Type** | **Population** | **Mean Performance (%)** | **SE Performance (%)** | | **Peak Performance (%)** | **SE Peak Performance (%)** | **Age of Peak Performance** |
| Mean | Full Sample | 82.56 | 0.55 | | 88.22 | 0.44 | 18.00 |
| Reward | Full Sample | 84.50 | 0.54 | | 90.31 | 0.42 | 17.89 |
| Neutral | Full Sample | 81.06 | 0.59 | | 87.64 | 0.48 | 18.04 |
| **Tissue Iron (nT2*w): Statistics Across Individual Regions of Interest** | | | | | | | |
| **Region of Interest** | **Population** | **Mean nt2*w** | **SE nT2*w** | | **Peak nT2*w** | **SE Peak nT2*w** | **Age of Peak nT2*w (years)** |
| Basal Ganglia | Full Sample | 0.0135 | .00001 | | 0.0130 | .000009 | 20.45 |
| Caudate | Full Sample | 0.0110 | .00004 | | 0.0094 | .00004 | 19.65 |
| NAcc | Full Sample | 0.0144 | .00001 | | 0.0138 | .000009 | 20.59 |
| Pallidum | Full Sample | 0.0140 | .00001 | | 0.0134 | .00001 | 19.91 |
| Putamen | Full Sample | 0.0111 | .00001 | | 0.0106 | .00001 | 20.85 |

**Supplementary Table S2. Sociodemographic Subgroups: Statistics Across Individual Substances, Subscales, Trial Types, and Regions of Interest**

| **Substance Use: Statistics Across Individual Substances by Sociodemographic Factors** | | | | | | | | | | | | | | | |
| --- | --- | --- | --- | --- | --- | --- | --- | --- | --- | --- | --- | --- | --- | --- | --- |
| **Substance** | **Population** | | **Level** | | **Mean n Days** | | **SE n Days** | | **Mean Peak Use (Days)** | | **SE Peak Use Days)** | | **Mean Age of Onset (Years)** | | **Mean Age of Peak Use (Years)** |
| All | Sex | | F | | 4.73 | | 0.14 | | 13.87 | | 0.19 | | 16.81 | | 22.08 |
| All | Sex | | M | | 6.64 | | 0.18 | | 17.67 | | 0.21 | | 16.51 | | 21.80 |
| All | Income | | <25k | | 5.51 | | 0.53 | | 16.82 | | 0.72 | | 16.06 | | 21.09 |
| All | Income | | 25-49k | | 4.93 | | 0.37 | | 15.58 | | 0.52 | | 17.04 | | 21.66 |
| All | Income | | 50-74k | | 5.10 | | 0.33 | | 14.86 | | 0.44 | | 16.86 | | 21.96 |
| All | Income | | 75-99k | | 6.62 | | 0.35 | | 18.71 | | 0.41 | | 16.40 | | 22.14 |
| All | Income | | 100k-199k | | 5.56 | | 0.17 | | 14.88 | | 0.22 | | 16.88 | | 22.24 |
| All | Income | | 200k+ | | 6.17 | | 0.26 | | 16.21 | | 0.29 | | 16.20 | | 21.50 |
| All | Race/ethnicity | | A, A, N | | 3.55 | | 0.31 | | 10.54 | | 0.47 | | 17.27 | | 21.77 |
| All | Race/ethnicity | | Hispanic | | 6.57 | | 0.36 | | 17.75 | | 0.41 | | 16.09 | | 21.82 |
| All | Race/ethnicity | | NH White | | 6.00 | | 0.15 | | 16.06 | | 0.18 | | 16.68 | | 22.08 |
| All | Race/ethnicity | | NH Black/AA | | 5.06 | | 0.34 | | 16.71 | | 0.45 | | 16.89 | | 21.86 |
| All | Race/ethnicity | | Unspecified | | 4.38 | | 0.50 | | 13.12 | | 0.68 | | 16.27 | | 20.73 |
| All | Site | | A | | 5.62 | | 0.30 | | 15.39 | | 0.39 | | 17.35 | | 22.54 |
| All | Site | | B | | 6.23 | | 0.27 | | 16.22 | | 0.31 | | 15.86 | | 21.19 |
| All | Site | | C | | 4.98 | | 0.24 | | 14.63 | | 0.32 | | 16.91 | | 21.96 |
| All | Site | | D | | 5.80 | | 0.26 | | 15.98 | | 0.33 | | 16.69 | | 21.87 |
| All | Site | | E | | 5.76 | | 0.22 | | 16.21 | | 0.28 | | 16.68 | | 22.23 |
| Alcohol | Sex | | F | | 2.25 | | 0.07 | | 7.70 | | 0.12 | | 17.06 | | 22.05 |
| Alcohol | Sex | | M | | 2.82 | | 0.09 | | 9.81 | | 0.14 | | 16.96 | | 22.18 |
| Alcohol | Income | | <25k | | 1.22 | | 0.15 | | 5.22 | | 0.28 | | 16.76 | | 21.20 |
| Alcohol | Income | | 25-49k | | 1.29 | | 0.13 | | 5.90 | | 0.28 | | 17.83 | | 22.05 |
| Alcohol | Income | | 50-74k | | 2.10 | | 0.16 | | 7.94 | | 0.28 | | 17.33 | | 22.14 |
| Alcohol | Income | | 75-99k | | 2.91 | | 0.18 | | 9.95 | | 0.27 | | 16.87 | | 22.37 |
| Alcohol | Income | | 100k-199k | | 2.77 | | 0.09 | | 8.97 | | 0.15 | | 17.13 | | 22.37 |
| Alcohol | Income | | 200k+ | | 3.06 | | 0.14 | | 9.97 | | 0.20 | | 16.38 | | 21.64 |
| Alcohol | Race/ethnicity | | A, A, N | | 1.71 | | 0.14 | | 5.90 | | 0.24 | | 17.62 | | 22.27 |
| Alcohol | Race/ethnicity | | Hispanic | | 2.24 | | 0.15 | | 8.30 | | 0.25 | | 16.53 | | 22.03 |
| Alcohol | Race/ethnicity | | NH White | | 2.92 | | 0.08 | | 9.57 | | 0.12 | | 16.94 | | 22.18 |
| Alcohol | Race/ethnicity | | NH Black/AA | | 1.53 | | 0.13 | | 6.58 | | 0.27 | | 17.59 | | 21.88 |
| Alcohol | Race/ethnicity | | Unspecified | | 1.91 | | 0.26 | | 7.82 | | 0.49 | | 16.99 | | 21.65 |
| Alcohol | Site | | A | | 2.47 | | 0.14 | | 8.09 | | 0.22 | | 17.80 | | 22.71 |
| Alcohol | Site | | B | | 2.68 | | 0.13 | | 8.91 | | 0.20 | | 16.08 | | 21.49 |
| Alcohol | Site | | C | | 2.27 | | 0.12 | | 8.13 | | 0.21 | | 17.28 | | 21.86 |
| Alcohol | Site | | D | | 3.11 | | 0.16 | | 10.01 | | 0.25 | | 17.01 | | 22.18 |
| Alcohol | Site | | E | | 2.26 | | 0.10 | | 8.44 | | 0.17 | | 17.10 | | 22.40 |
| Binge | Sex | | F | | 0.52 | | 0.03 | | 3.41 | | 0.09 | | 18.34 | | 21.34 |
| Binge | Sex | | M | | 0.74 | | 0.04 | | 4.33 | | 0.10 | | 18.07 | | 21.17 |
| Binge | Income | | <25k | | 0.31 | | 0.05 | | 2.76 | | 0.17 | | 18.72 | | 21.66 |
| Binge | Income | | 25-49k | | 0.33 | | 0.06 | | 3.29 | | 0.26 | | 18.99 | | 21.54 |
| Binge | Income | | 50-74k | | 0.55 | | 0.08 | | 4.17 | | 0.27 | | 18.73 | | 21.14 |
| Binge | Income | | 75-99k | | 0.62 | | 0.06 | | 3.73 | | 0.18 | | 18.17 | | 21.45 |
| Binge | Income | | 100k-199k | | 0.66 | | 0.04 | | 3.91 | | 0.10 | | 18.33 | | 21.40 |
| Binge | Income | | 200k+ | | 0.85 | | 0.06 | | 4.16 | | 0.12 | | 17.39 | | 20.73 |
| Binge | Race/ethnicity | | A, A, N | | 0.43 | | 0.06 | | 3.19 | | 0.16 | | 18.87 | | 21.65 |
| Binge | Race/ethnicity | | Hispanic | | 0.47 | | 0.05 | | 3.07 | | 0.15 | | 18.09 | | 21.20 |
| Binge | Race/ethnicity | | NH White | | 0.71 | | 0.03 | | 4.06 | | 0.08 | | 18.07 | | 21.20 |
| Binge | Race/ethnicity | | NH Black/AA | | 0.40 | | 0.06 | | 3.48 | | 0.24 | | 19.06 | | 21.47 |
| Binge | Race/ethnicity | | Unspecified | | 0.78 | | 0.16 | | 5.50 | | 0.49 | | 17.77 | | 20.97 |
| Binge | Site | | A | | 0.62 | | 0.06 | | 4.38 | | 0.20 | | 18.68 | | 21.73 |
| Binge | Site | | B | | 0.82 | | 0.06 | | 4.38 | | 0.14 | | 17.33 | | 20.42 |
| Binge | Site | | C | | 0.60 | | 0.05 | | 3.65 | | 0.15 | | 18.38 | | 21.38 |
| Binge | Site | | D | | 0.58 | | 0.05 | | 3.65 | | 0.14 | | 18.58 | | 21.42 |
| Binge | Site | | E | | 0.55 | | 0.04 | | 3.55 | | 0.13 | | 18.24 | | 21.48 |
| Cannabis | Sex | | F | | 2.12 | | 0.11 | | 11.60 | | 0.27 | | 17.54 | | 21.62 |
| Cannabis | Sex | | M | | 3.33 | | 0.14 | | 15.08 | | 0.28 | | 16.82 | | 21.15 |
| Cannabis | Income | | <25k | | 3.59 | | 0.47 | | 16.70 | | 0.93 | | 16.04 | | 20.56 |
| Cannabis | Income | | 25-49k | | 3.53 | | 0.34 | | 18.29 | | 0.67 | | 17.11 | | 21.46 |
| Cannabis | Income | | 50-74k | | 2.66 | | 0.27 | | 12.54 | | 0.56 | | 17.26 | | 21.18 |
| Cannabis | Income | | 75-99k | | 3.14 | | 0.28 | | 14.54 | | 0.56 | | 17.08 | | 21.84 |
| Cannabis | Income | | 100k-199k | | 2.52 | | 0.14 | | 12.70 | | 0.31 | | 17.48 | | 21.79 |
| Cannabis | Income | | 200k+ | | 2.29 | | 0.18 | | 11.36 | | 0.40 | | 16.85 | | 20.63 |
| Cannabis | Race/ethnicity | | A, A, N | | 1.06 | | 0.19 | | 7.61 | | 0.62 | | 17.54 | | 21.17 |
| Cannabis | Race/ethnicity | | Hispanic | | 3.70 | | 0.30 | | 14.25 | | 0.52 | | 16.68 | | 21.42 |
| Cannabis | Race/ethnicity | | NH White | | 2.79 | | 0.12 | | 13.88 | | 0.25 | | 17.27 | | 21.46 |
| Cannabis | Race/ethnicity | | NH Black/AA | | 2.80 | | 0.27 | | 14.05 | | 0.56 | | 17.16 | | 21.53 |
| Cannabis | Race/ethnicity | | Unspecified | | 1.97 | | 0.37 | | 9.80 | | 0.86 | | 16.50 | | 20.25 |
| Cannabis | Site | | A | | 2.35 | | 0.23 | | 14.62 | | 0.60 | | 17.55 | | 22.06 |
| Cannabis | Site | | B | | 3.06 | | 0.21 | | 12.93 | | 0.41 | | 16.65 | | 20.62 |
| Cannabis | Site | | C | | 2.05 | | 0.18 | | 11.61 | | 0.44 | | 17.51 | | 21.81 |
| Cannabis | Site | | D | | 2.74 | | 0.21 | | 13.76 | | 0.44 | | 17.38 | | 21.51 |
| Cannabis | Site | | E | | 3.14 | | 0.19 | | 14.00 | | 0.38 | | 17.02 | | 21.37 |
| Nicotine | Sex | | F | | 1.00 | | 0.09 | | 12.00 | | 0.41 | | 17.96 | | 21.28 |
| Nicotine | Sex | | M | | 2.13 | | 0.12 | | 14.55 | | 0.33 | | 17.61 | | 20.88 |
| Nicotine | Income | | <25k | | 1.93 | | 0.38 | | 20.20 | | 1.15 | | 17.33 | | 20.86 |
| Nicotine | Income | | 25-49k | | 1.60 | | 0.25 | | 16.07 | | 0.84 | | 17.00 | | 20.68 |
| Nicotine | Income | | 50-74k | | 1.39 | | 0.21 | | 12.53 | | 0.87 | | 17.74 | | 20.68 |
| Nicotine | Income | | 75-99k | | 1.96 | | 0.24 | | 14.72 | | 0.69 | | 17.58 | | 21.19 |
| Nicotine | Income | | 100k-199k | | 1.43 | | 0.11 | | 12.93 | | 0.40 | | 17.88 | | 21.22 |
| Nicotine | Income | | 200k+ | | 1.58 | | 0.16 | | 12.52 | | 0.52 | | 17.98 | | 20.90 |
| Nicotine | Race/ethnicity | | A, A, N | | 1.10 | | 0.22 | | 10.23 | | 0.89 | | 17.82 | | 20.55 |
| Nicotine | Race/ethnicity | | Hispanic | | 1.78 | | 0.22 | | 15.51 | | 0.72 | | 17.13 | | 21.17 |
| Nicotine | Race/ethnicity | | NH White | | 1.54 | | 0.09 | | 13.00 | | 0.31 | | 17.81 | | 21.11 |
| Nicotine | Race/ethnicity | | NH Black/AA | | 1.70 | | 0.24 | | 16.21 | | 0.86 | | 17.95 | | 20.90 |
| Nicotine | Race/ethnicity | | Unspecified | | 1.78 | | 0.40 | | 18.28 | | 1.39 | | 18.40 | | 20.49 |
| Nicotine | Site | | A | | 1.78 | | 0.22 | | 17.12 | | 0.84 | | 17.74 | | 21.52 |
| Nicotine | Site | | B | | 1.64 | | 0.17 | | 13.04 | | 0.51 | | 17.61 | | 20.55 |
| Nicotine | Site | | C | | 1.40 | | 0.16 | | 11.57 | | 0.57 | | 18.00 | | 21.24 |
| Nicotine | Site | | D | | 1.54 | | 0.17 | | 13.22 | | 0.55 | | 17.93 | | 21.14 |
| Nicotine | Site | | E | | 1.54 | | 0.14 | | 14.28 | | 0.50 | | 17.50 | | 21.00 |
| **Impulsivity (UPPS): Statistics Across Individual Subscales by Sociodemographic Factors** | | | | | | | | | | | | | | | |
| **Subscale** | | **Population** | | **Level** | | **Mean Score** | | **SE Score** | | **Mean Peak Score** | | **SE Peak Score** | | **Mean Age of Peak Score (Years)** | |
| Mean | | Sex | | F | | 1.86 | | 0.01 | | 2.17 | | 0.01 | | 18.27 | |
| Mean | | Sex | | M | | 1.98 | | 0.01 | | 2.27 | | 0.01 | | 18.15 | |
| Mean | | Income | | <25k | | 1.95 | | 0.03 | | 2.26 | | 0.02 | | 17.53 | |
| Mean | | Income | | 25-49k | | 1.93 | | 0.02 | | 2.27 | | 0.02 | | 18.21 | |
| Mean | | Income | | 50-74k | | 1.90 | | 0.02 | | 2.23 | | 0.01 | | 17.59 | |
| Mean | | Income | | 75-99k | | 1.97 | | 0.02 | | 2.26 | | 0.02 | | 18.15 | |
| Mean | | Income | | 100k-199k | | 1.91 | | 0.01 | | 2.21 | | 0.01 | | 18.80 | |
| Mean | | Income | | 200k+ | | 1.92 | | 0.01 | | 2.18 | | 0.01 | | 17.50 | |
| Mean | | Race/ethnicity | | A, A, N | | 2.04 | | 0.02 | | 2.28 | | 0.02 | | 18.67 | |
| Mean | | Race/ethnicity | | Hispanic | | 1.96 | | 0.02 | | 2.27 | | 0.01 | | 17.79 | |
| Mean | | Race/ethnicity | | NH White | | 1.92 | | 0.01 | | 2.21 | | 0.01 | | 18.27 | |
| Mean | | Race/ethnicity | | NH Black/AA | | 1.86 | | 0.02 | | 2.21 | | 0.02 | | 18.13 | |
| Mean | | Race/ethnicity | | Unspecified | | 1.89 | | 0.03 | | 2.16 | | 0.02 | | 17.88 | |
| Mean | | Site | | A | | 1.87 | | 0.02 | | 2.16 | | 0.01 | | 18.84 | |
| Mean | | Site | | B | | 1.98 | | 0.01 | | 2.23 | | 0.01 | | 17.59 | |
| Mean | | Site | | C | | 1.86 | | 0.01 | | 2.19 | | 0.01 | | 17.96 | |
| Mean | | Site | | D | | 1.97 | | 0.01 | | 2.29 | | 0.01 | | 18.57 | |
| Mean | | Site | | E | | 1.92 | | 0.01 | | 2.21 | | 0.01 | | 18.24 | |
| Negative Urgency | | Sex | | F | | 1.85 | | 0.01 | | 2.47 | | 0.01 | | 18.24 | |
| Negative Urgency | | Sex | | M | | 1.87 | | 0.01 | | 2.50 | | 0.01 | | 18.26 | |
| Negative Urgency | | Income | | <25k | | 2.03 | | 0.05 | | 2.70 | | 0.05 | | 17.50 | |
| Negative Urgency | | Income | | 25-49k | | 1.85 | | 0.03 | | 2.50 | | 0.03 | | 18.20 | |
| Negative Urgency | | Income | | 50-74k | | 1.87 | | 0.03 | | 2.53 | | 0.03 | | 17.88 | |
| Negative Urgency | | Income | | 75-99k | | 1.89 | | 0.03 | | 2.56 | | 0.03 | | 18.36 | |
| Negative Urgency | | Income | | 100k-199k | | 1.84 | | 0.02 | | 2.46 | | 0.01 | | 18.86 | |
| Negative Urgency | | Income | | 200k+ | | 1.82 | | 0.02 | | 2.40 | | 0.02 | | 17.30 | |
| Negative Urgency | | Race/ethnicity | | A, A, N | | 2.00 | | 0.04 | | 2.63 | | 0.03 | | 18.60 | |
| Negative Urgency | | Race/ethnicity | | Hispanic | | 1.96 | | 0.03 | | 2.56 | | 0.03 | | 17.81 | |
| Negative Urgency | | Race/ethnicity | | NH White | | 1.83 | | 0.01 | | 2.44 | | 0.01 | | 18.43 | |
| Negative Urgency | | Race/ethnicity | | NH Black/AA | | 1.83 | | 0.03 | | 2.54 | | 0.03 | | 17.84 | |
| Negative Urgency | | Race/ethnicity | | Unspecified | | 1.88 | | 0.05 | | 2.49 | | 0.04 | | 17.32 | |
| Negative Urgency | | Site | | A | | 1.82 | | 0.03 | | 2.46 | | 0.03 | | 19.19 | |
| Negative Urgency | | Site | | B | | 1.96 | | 0.02 | | 2.53 | | 0.02 | | 17.53 | |
| Negative Urgency | | Site | | C | | 1.77 | | 0.02 | | 2.43 | | 0.02 | | 17.80 | |
| Negative Urgency | | Site | | D | | 1.93 | | 0.02 | | 2.56 | | 0.02 | | 18.70 | |
| Negative Urgency | | Site | | E | | 1.82 | | 0.02 | | 2.45 | | 0.02 | | 18.28 | |
| Perseverance | | Sex | | F | | 1.76 | | 0.01 | | 2.20 | | 0.01 | | 18.33 | |
| Perseverance | | Sex | | M | | 1.76 | | 0.01 | | 2.16 | | 0.01 | | 18.09 | |
| Perseverance | | Income | | <25k | | 1.84 | | 0.03 | | 2.35 | | 0.03 | | 17.93 | |
| Perseverance | | Income | | 25-49k | | 1.76 | | 0.02 | | 2.19 | | 0.02 | | 18.05 | |
| Perseverance | | Income | | 50-74k | | 1.70 | | 0.02 | | 2.16 | | 0.02 | | 18.11 | |
| Perseverance | | Income | | 75-99k | | 1.78 | | 0.02 | | 2.20 | | 0.02 | | 18.35 | |
| Perseverance | | Income | | 100k-199k | | 1.75 | | 0.01 | | 2.17 | | 0.01 | | 18.65 | |
| Perseverance | | Income | | 200k+ | | 1.79 | | 0.02 | | 2.17 | | 0.01 | | 17.36 | |
| Perseverance | | Race/ethnicity | | A, A, N | | 1.75 | | 0.03 | | 2.11 | | 0.02 | | 18.61 | |
| Perseverance | | Race/ethnicity | | Hispanic | | 1.82 | | 0.02 | | 2.23 | | 0.01 | | 18.03 | |
| Perseverance | | Race/ethnicity | | NH White | | 1.77 | | 0.01 | | 2.18 | | 0.01 | | 18.32 | |
| Perseverance | | Race/ethnicity | | NH Black/AA | | 1.71 | | 0.02 | | 2.19 | | 0.02 | | 17.75 | |
| Perseverance | | Race/ethnicity | | Unspecified | | 1.71 | | 0.03 | | 2.18 | | 0.03 | | 17.49 | |
| Perseverance | | Site | | A | | 1.74 | | 0.02 | | 2.15 | | 0.02 | | 19.12 | |
| Perseverance | | Site | | B | | 1.82 | | 0.02 | | 2.19 | | 0.01 | | 17.37 | |
| Perseverance | | Site | | C | | 1.75 | | 0.02 | | 2.16 | | 0.01 | | 17.98 | |
| Perseverance | | Site | | D | | 1.79 | | 0.02 | | 2.23 | | 0.01 | | 18.50 | |
| Perseverance | | Site | | E | | 1.73 | | 0.01 | | 2.17 | | 0.01 | | 18.28 | |
| Positive Urgency | | Sex | | F | | 1.54 | | 0.01 | | 2.04 | | 0.01 | | 17.74 | |
| Positive Urgency | | Sex | | M | | 1.75 | | 0.01 | | 2.32 | | 0.01 | | 17.93 | |
| Positive Urgency | | Income | | <25k | | 1.84 | | 0.05 | | 2.48 | | 0.05 | | 17.48 | |
| Positive Urgency | | Income | | 25-49k | | 1.71 | | 0.03 | | 2.30 | | 0.03 | | 18.04 | |
| Positive Urgency | | Income | | 50-74k | | 1.67 | | 0.03 | | 2.23 | | 0.03 | | 17.30 | |
| Positive Urgency | | Income | | 75-99k | | 1.72 | | 0.03 | | 2.26 | | 0.03 | | 17.93 | |
| Positive Urgency | | Income | | 100k-199k | | 1.61 | | 0.01 | | 2.13 | | 0.01 | | 18.30 | |
| Positive Urgency | | Income | | 200k+ | | 1.56 | | 0.02 | | 2.06 | | 0.02 | | 17.05 | |
| Positive Urgency | | Race/ethnicity | | A, A, N | | 1.83 | | 0.04 | | 2.32 | | 0.04 | | 18.22 | |
| Positive Urgency | | Race/ethnicity | | Hispanic | | 1.70 | | 0.03 | | 2.29 | | 0.03 | | 17.56 | |
| Positive Urgency | | Race/ethnicity | | NH White | | 1.60 | | 0.01 | | 2.12 | | 0.01 | | 17.91 | |
| Positive Urgency | | Race/ethnicity | | NH Black/AA | | 1.70 | | 0.03 | | 2.32 | | 0.03 | | 17.62 | |
| Positive Urgency | | Race/ethnicity | | Unspecified | | 1.61 | | 0.04 | | 2.10 | | 0.04 | | 17.40 | |
| Positive Urgency | | Site | | A | | 1.66 | | 0.03 | | 2.16 | | 0.03 | | 18.35 | |
| Positive Urgency | | Site | | B | | 1.70 | | 0.02 | | 2.18 | | 0.02 | | 17.00 | |
| Positive Urgency | | Site | | C | | 1.56 | | 0.02 | | 2.15 | | 0.02 | | 17.61 | |
| Positive Urgency | | Site | | D | | 1.66 | | 0.02 | | 2.25 | | 0.02 | | 18.22 | |
| Positive Urgency | | Site | | E | | 1.64 | | 0.02 | | 2.17 | | 0.02 | | 18.04 | |
| Premeditation | | Sex | | F | | 1.63 | | 0.01 | | 2.10 | | 0.01 | | 18.24 | |
| Premeditation | | Sex | | M | | 1.64 | | 0.01 | | 2.06 | | 0.01 | | 18.09 | |
| Premeditation | | Income | | <25k | | 1.58 | | 0.03 | | 2.07 | | 0.03 | | 17.23 | |
| Premeditation | | Income | | 25-49k | | 1.67 | | 0.02 | | 2.16 | | 0.02 | | 18.17 | |
| Premeditation | | Income | | 50-74k | | 1.59 | | 0.02 | | 2.10 | | 0.02 | | 17.45 | |
| Premeditation | | Income | | 75-99k | | 1.69 | | 0.02 | | 2.13 | | 0.02 | | 18.16 | |
| Premeditation | | Income | | 100k-199k | | 1.64 | | 0.01 | | 2.06 | | 0.01 | | 18.68 | |
| Premeditation | | Income | | 200k+ | | 1.62 | | 0.02 | | 2.04 | | 0.01 | | 17.72 | |
| Premeditation | | Race/ethnicity | | A, A, N | | 1.67 | | 0.03 | | 2.02 | | 0.02 | | 18.38 | |
| Premeditation | | Race/ethnicity | | Hispanic | | 1.63 | | 0.02 | | 2.06 | | 0.02 | | 18.01 | |
| Premeditation | | Race/ethnicity | | NH White | | 1.65 | | 0.01 | | 2.10 | | 0.01 | | 18.30 | |
| Premeditation | | Race/ethnicity | | NH Black/AA | | 1.55 | | 0.02 | | 2.07 | | 0.02 | | 17.56 | |
| Premeditation | | Race/ethnicity | | Unspecified | | 1.63 | | 0.04 | | 2.04 | | 0.04 | | 17.77 | |
| Premeditation | | Site | | A | | 1.61 | | 0.02 | | 2.01 | | 0.02 | | 18.67 | |
| Premeditation | | Site | | B | | 1.68 | | 0.02 | | 2.07 | | 0.02 | | 17.42 | |
| Premeditation | | Site | | C | | 1.56 | | 0.02 | | 2.02 | | 0.02 | | 17.99 | |
| Premeditation | | Site | | D | | 1.72 | | 0.02 | | 2.20 | | 0.02 | | 18.20 | |
| Premeditation | | Site | | E | | 1.62 | | 0.01 | | 2.09 | | 0.01 | | 18.53 | |
| Sensation Seeking | | Sex | | F | | 2.52 | | 0.01 | | 3.05 | | 0.01 | | 17.90 | |
| Sensation Seeking | | Sex | | M | | 2.90 | | 0.01 | | 3.35 | | 0.01 | | 18.22 | |
| Sensation Seeking | | Income | | <25k | | 2.47 | | 0.05 | | 3.10 | | 0.03 | | 17.46 | |
| Sensation Seeking | | Income | | 25-49k | | 2.64 | | 0.03 | | 3.16 | | 0.03 | | 18.13 | |
| Sensation Seeking | | Income | | 50-74k | | 2.67 | | 0.03 | | 3.19 | | 0.02 | | 17.97 | |
| Sensation Seeking | | Income | | 75-99k | | 2.75 | | 0.03 | | 3.21 | | 0.02 | | 18.06 | |
| Sensation Seeking | | Income | | 100k-199k | | 2.72 | | 0.02 | | 3.20 | | 0.01 | | 18.49 | |
| Sensation Seeking | | Income | | 200k+ | | 2.82 | | 0.02 | | 3.23 | | 0.02 | | 17.33 | |
| Sensation Seeking | | Race/ethnicity | | A, A, N | | 2.93 | | 0.03 | | 3.32 | | 0.02 | | 18.35 | |
| Sensation Seeking | | Race/ethnicity | | Hispanic | | 2.67 | | 0.03 | | 3.20 | | 0.02 | | 17.76 | |
| Sensation Seeking | | Race/ethnicity | | NH White | | 2.74 | | 0.01 | | 3.20 | | 0.01 | | 18.18 | |
| Sensation Seeking | | Race/ethnicity | | NH Black/AA | | 2.52 | | 0.03 | | 3.13 | | 0.03 | | 17.97 | |
| Sensation Seeking | | Race/ethnicity | | Unspecified | | 2.62 | | 0.05 | | 3.07 | | 0.04 | | 16.96 | |
| Sensation Seeking | | Site | | A | | 2.56 | | 0.03 | | 3.06 | | 0.02 | | 18.58 | |
| Sensation Seeking | | Site | | B | | 2.74 | | 0.02 | | 3.16 | | 0.02 | | 17.34 | |
| Sensation Seeking | | Site | | C | | 2.65 | | 0.02 | | 3.16 | | 0.02 | | 17.63 | |
| Sensation Seeking | | Site | | D | | 2.76 | | 0.02 | | 3.28 | | 0.02 | | 18.26 | |
| Sensation Seeking | | Site | | E | | 2.78 | | 0.02 | | 3.26 | | 0.01 | | 18.48 | |
| **Inhibitory Control (Anti-Saccade): Statistics Across Individual Trial Types by Sociodemographic Factors** | | | | | | | | | | | | | | | |
| **Trial Type** | | **Population** | | **Level** | | **Mean performance (%)** | | **SE (%)** | | **Mean Peak Performance (%)** | | **SE Peak Performance (%)** | | **Mean Age of Peak Performance (Years)** | |
| Mean | | Sex | | F | | 82.61 | | 0.75 | | 88.40 | | 0.62 | | 18.15 | |
| Mean | | Sex | | M | | 82.49 | | 0.80 | | 88.03 | | 0.64 | | 17.83 | |
| Mean | | Income | | <25k | | 77.29 | | 2.32 | | 83.11 | | 1.83 | | 17.79 | |
| Mean | | Income | | 25-49k | | 82.61 | | 1.38 | | 88.37 | | 1.08 | | 17.84 | |
| Mean | | Income | | 50-74k | | 83.75 | | 1.92 | | 88.69 | | 1.58 | | 17.55 | |
| Mean | | Income | | 75-99k | | 82.33 | | 1.32 | | 88.17 | | 1.09 | | 18.03 | |
| Mean | | Income | | 100k-199k | | 82.62 | | 0.79 | | 88.41 | | 0.66 | | 18.13 | |
| Mean | | Income | | 200k+ | | 86.49 | | 1.91 | | 91.61 | | 1.23 | | 18.08 | |
| Mean | | Race/ethnicity | | A, A, N | | 91.60 | | 2.75 | | 94.28 | | 2.18 | | 17.36 | |
| Mean | | Race/ethnicity | | Hispanic | | 87.30 | | 2.56 | | 91.57 | | 1.29 | | 17.12 | |
| Mean | | Race/ethnicity | | NH White | | 82.51 | | 0.67 | | 88.12 | | 0.55 | | 18.21 | |
| Mean | | Race/ethnicity | | NH Black/AA | | 80.87 | | 1.13 | | 86.57 | | 0.93 | | 17.58 | |
| Mean | | Race/ethnicity | | Unspecified | | 86.55 | | 2.69 | | 94.84 | | 1.02 | | 17.94 | |
| Mean | | Site | | A | | 79.78 | | 0.73 | | 86.05 | | 0.64 | | 18.50 | |
| Mean | | Site | | C | | 85.96 | | 0.78 | | 90.89 | | 0.57 | | 17.37 | |
| Reward | | Sex | | F | | 84.55 | | 0.73 | | 90.39 | | 0.57 | | 18.05 | |
| Reward | | Sex | | M | | 84.44 | | 0.81 | | 90.23 | | 0.61 | | 17.71 | |
| Reward | | Income | | <25k | | 79.49 | | 2.46 | | 86.47 | | 1.68 | | 17.66 | |
| Reward | | Income | | 25-49k | | 84.99 | | 1.37 | | 91.20 | | 1.05 | | 17.95 | |
| Reward | | Income | | 50-74k | | 85.12 | | 1.91 | | 90.29 | | 1.55 | | 17.43 | |
| Reward | | Income | | 75-99k | | 84.03 | | 1.34 | | 89.63 | | 1.09 | | 18.00 | |
| Reward | | Income | | 100k-199k | | 84.42 | | 0.77 | | 90.36 | | 0.59 | | 17.92 | |
| Reward | | Income | | 200k+ | | 89.75 | | 1.63 | | 94.28 | | 0.93 | | 18.07 | |
| Reward | | Race/ethnicity | | A, A, N | | 93.91 | | 1.83 | | 95.14 | | 1.79 | | 17.36 | |
| Reward | | Race/ethnicity | | Hispanic | | 89.98 | | 2.22 | | 94.64 | | 0.96 | | 17.00 | |
| Reward | | Race/ethnicity | | NH White | | 84.26 | | 0.66 | | 90.09 | | 0.52 | | 18.06 | |
| Reward | | Race/ethnicity | | NH Black/AA | | 83.24 | | 1.15 | | 88.97 | | 0.87 | | 17.60 | |
| Reward | | Race/ethnicity | | Unspecified | | 88.47 | | 2.80 | | 96.83 | | 0.79 | | 17.66 | |
| Reward | | Site | | A | | 81.94 | | 0.74 | | 88.46 | | 0.58 | | 18.41 | |
| Reward | | Site | | C | | 87.67 | | 0.76 | | 92.61 | | 0.57 | | 17.24 | |
| Neutral | | Sex | | F | | 80.83 | | 0.84 | | 87.91 | | 0.67 | | 18.20 | |
| Neutral | | Sex | | M | | 81.31 | | 0.83 | | 87.35 | | 0.68 | | 17.86 | |
| Neutral | | Income | | <25k | | 75.72 | | 2.37 | | 81.70 | | 2.00 | | 17.98 | |
| Neutral | | Income | | 25-49k | | 80.23 | | 1.48 | | 86.24 | | 1.22 | | 17.84 | |
| Neutral | | Income | | 50-74k | | 82.39 | | 2.06 | | 87.41 | | 1.73 | | 17.32 | |
| Neutral | | Income | | 75-99k | | 81.52 | | 1.35 | | 88.15 | | 1.10 | | 18.09 | |
| Neutral | | Income | | 100k-199k | | 81.07 | | 0.90 | | 88.33 | | 0.70 | | 18.19 | |
| Neutral | | Income | | 200k+ | | 84.84 | | 2.00 | | 90.77 | | 1.44 | | 18.18 | |
| Neutral | | Race/ethnicity | | A, A, N | | 89.29 | | 3.90 | | 93.42 | | 2.62 | | 17.36 | |
| Neutral | | Race/ethnicity | | Hispanic | | 84.62 | | 3.12 | | 90.50 | | 1.58 | | 16.77 | |
| Neutral | | Race/ethnicity | | NH White | | 81.12 | | 0.73 | | 87.76 | | 0.59 | | 18.27 | |
| Neutral | | Race/ethnicity | | NH Black/AA | | 79.10 | | 1.21 | | 85.72 | | 1.02 | | 17.68 | |
| Neutral | | Race/ethnicity | | Unspecified | | 86.03 | | 2.44 | | 92.96 | | 1.25 | | 17.56 | |
| Neutral | | Site | | A | | 78.16 | | 0.79 | | 85.49 | | 0.69 | | 18.59 | |
| Neutral | | Site | | C | | 84.56 | | 0.85 | | 90.25 | | 0.61 | | 17.36 | |
| **Tissue Iron (nT2*w): Statistics Across Individual Regions by Sociodemographic Factors** | | | | | | | | | | | | | | | |
| **Region** | | **Population** | | **Level** | | **Mean nT2*w** | | **SE nT2*w** | | **Mean Peak nT2*w** | | **SE Peak nT2*w** | | **Mean Age of Peak nT2*w (Years)** | |
| Basal Ganglia | | Sex | | F | | 0.0137 | | .00001 | | 0.0131 | | .00001 | | 20.64 | |
| Basal Ganglia | | Sex | | M | | 0.0134 | | .00001 | | 0.0129 | | .00001 | | 20.27 | |
| Basal Ganglia | | Income | | <25k | | 0.0137 | | .00005 | | 0.0132 | | .00004 | | 19.71 | |
| Basal Ganglia | | Income | | 25-49k | | 0.0137 | | .00001 | | 0.0131 | | .00001 | | 20.28 | |
| Basal Ganglia | | Income | | 50-74k | | 0.0135 | | .00002 | | 0.0129 | | .00002 | | 19.98 | |
| Basal Ganglia | | Income | | 75-99k | | 0.0135 | | .00003 | | 0.0129 | | .00003 | | 19.84 | |
| Basal Ganglia | | Income | | 100k-199k | | 0.0135 | | .00002 | | 0.0130 | | .00002 | | 21.14 | |
| Basal Ganglia | | Income | | 200k+ | | 0.0135 | | .00002 | | 0.0130 | | .00002 | | 19.93 | |
| Basal Ganglia | | Race/eth | | A, A, N | | 0.0136 | | .00004 | | 0.0131 | | .00004 | | 20.45 | |
| Basal Ganglia | | Race/eth | | Hispanic | | 0.0137 | | .00003 | | 0.0132 | | .00002 | | 20.67 | |
| Basal Ganglia | | Race/eth | | NH White | | 0.0135 | | .00003 | | 0.0130 | | .00003 | | 20.46 | |
| Basal Ganglia | | Race/eth | | NH Black/AA | | 0.0137 | | .00001 | | 0.0131 | | .00001 | | 20.25 | |
| Basal Ganglia | | Race/eth | | Unspecified | | 0.0135 | | .00004 | | 0.0129 | | .00003 | | 20.32 | |
| Basal Ganglia | | Site | | A | | 0.0135 | | .00002 | | 0.0129 | | .00002 | | 21.05 | |
| Basal Ganglia | | Site | | B | | 0.0136 | | .00002 | | 0.0131 | | .00002 | | 20.39 | |
| Basal Ganglia | | Site | | C | | 0.0135 | | .00002 | | 0.0130 | | .00002 | | 19.69 | |
| Basal Ganglia | | Site | | D | | 0.0135 | | .00002 | | 0.0130 | | .00001 | | 19.98 | |
| Basal Ganglia | | Site | | E | | 0.0135 | | .00001 | | 0.0130 | | .00001 | | 21.06 | |
| Caudate | | Sex | | F | | 0.0142 | | .00001 | | 0.0136 | | .00001 | | 20.05 | |
| Caudate | | Sex | | M | | 0.0138 | | .00001 | | 0.0132 | | .00001 | | 19.76 | |
| Caudate | | Income | | <25k | | 0.0142 | | .00005 | | 0.0136 | | .00005 | | 18.94 | |
| Caudate | | Income | | 25-49k | | 0.0141 | | .00001 | | 0.0136 | | .00001 | | 19.84 | |
| Caudate | | Income | | 50-74k | | 0.0139 | | .00002 | | 0.0133 | | .00002 | | 19.58 | |
| Caudate | | Income | | 75-99k | | 0.0139 | | .00003 | | 0.0133 | | .00003 | | 19.28 | |
| Caudate | | Income | | 100k-199k | | 0.0140 | | .00003 | | 0.0134 | | .00002 | | 20.48 | |
| Caudate | | Income | | 200k+ | | 0.0140 | | .00002 | | 0.0134 | | .00002 | | 19.56 | |
| Caudate | | Race/eth | | A, A, N | | 0.0140 | | .00004 | | 0.0134 | | .00004 | | 20.24 | |
| Caudate | | Race/eth | | Hispanic | | 0.0142 | | .00003 | | 0.0136 | | .00003 | | 20.20 | |
| Caudate | | Race/eth | | NH White | | 0.0139 | | .00003 | | 0.0134 | | .00003 | | 19.90 | |
| Caudate | | Race/eth | | NH Black/AA | | 0.0141 | | .00001 | | 0.0134 | | .00001 | | 19.39 | |
| Caudate | | Race/eth | | Unspecified | | 0.0139 | | .00005 | | 0.0133 | | .00004 | | 19.93 | |
| Caudate | | Site | | A | | 0.0140 | | .00003 | | 0.0133 | | .00002 | | 19.83 | |
| Caudate | | Site | | B | | 0.0140 | | .00002 | | 0.0134 | | .00002 | | 20.41 | |
| Caudate | | Site | | C | | 0.0140 | | .00002 | | 0.0134 | | .00002 | | 19.20 | |
| Caudate | | Site | | D | | 0.0140 | | .00002 | | 0.0133 | | .00002 | | 19.59 | |
| Caudate | | Site | | E | | 0.0140 | | .00002 | | 0.0135 | | .00002 | | 20.34 | |
| NAcc | | Sex | | F | | 0.0110 | | .00005 | | 0.0094 | | .00005 | | 19.68 | |
| NAcc | | Sex | | M | | 0.0109 | | .00005 | | 0.0095 | | .00005 | | 19.62 | |
| NAcc | | Income | | <25k | | 0.0119 | | 0.0001 | | 0.0106 | | 0.0001 | | 19.40 | |
| NAcc | | Income | | 25-49k | | 0.0113 | | .00005 | | 0.0096 | | .00005 | | 19.73 | |
| NAcc | | Income | | 50-74k | | 0.0106 | | .00008 | | 0.0091 | | .00008 | | 19.24 | |
| NAcc | | Income | | 75-99k | | 0.0109 | | 0.0001 | | 0.0094 | | 0.0001 | | 19.42 | |
| NAcc | | Income | | 100k-199k | | 0.0109 | | 0.0001 | | 0.0094 | | .00009 | | 20.11 | |
| NAcc | | Income | | 200k+ | | 0.0110 | | 0.0001 | | 0.0094 | | 0.0001 | | 19.04 | |
| NAcc | | Race/ethnicity | | A, A, N | | 0.0117 | | 0.0001 | | 0.0103 | | 0.0001 | | 19.88 | |
| NAcc | | Race/ethnicity | | Hispanic | | 0.0114 | | 0.0001 | | 0.0101 | | 0.0001 | | 19.97 | |
| NAcc | | Race/ethnicity | | NH White | | 0.0108 | | 0.0001 | | 0.0093 | | 0.0001 | | 19.57 | |
| NAcc | | Race/ethnicity | | NH Black/AA | | 0.0109 | | .00004 | | 0.0093 | | .00004 | | 19.59 | |
| NAcc | | Race/ethnicity | | Unspecified | | 0.0106 | | 0.0001 | | 0.0089 | | 0.0001 | | 19.59 | |
| NAcc | | Site | | A | | 0.0109 | | 0.0001 | | 0.0093 | | 0.0001 | | 19.87 | |
| NAcc | | Site | | B | | 0.0110 | | .00009 | | 0.0096 | | .00008 | | 19.50 | |
| NAcc | | Site | | C | | 0.0109 | | .00007 | | 0.0094 | | .00007 | | 19.14 | |
| NAcc | | Site | | D | | 0.0110 | | .00008 | | 0.0092 | | .00008 | | 19.43 | |
| NAcc | | Site | | E | | 0.0110 | | .00006 | | 0.0096 | | .00007 | | 20.13 | |
| Pallidum | | Sex | | F | | 0.0112 | | .00001 | | 0.0106 | | .00001 | | 20.99 | |
| Pallidum | | Sex | | M | | 0.0110 | | .00001 | | 0.0106 | | .00001 | | 20.71 | |
| Pallidum | | Income | | <25k | | 0.0112 | | .00005 | | 0.0107 | | .00004 | | 20.45 | |
| Pallidum | | Income | | 25-49k | | 0.0112 | | .00001 | | 0.0107 | | .00001 | | 20.99 | |
| Pallidum | | Income | | 50-74k | | 0.0111 | | .00002 | | 0.0106 | | .00002 | | 20.69 | |
| Pallidum | | Income | | 75-99k | | 0.0111 | | .00003 | | 0.0106 | | .00002 | | 20.62 | |
| Pallidum | | Income | | 100k-199k | | 0.0111 | | .00003 | | 0.0106 | | .00003 | | 21.27 | |
| Pallidum | | Income | | 200k+ | | 0.0111 | | .00003 | | 0.0107 | | .00002 | | 20.18 | |
| Pallidum | | Race/ethnicity | | A, A, N | | 0.0112 | | .00004 | | 0.0107 | | .00004 | | 20.84 | |
| Pallidum | | Race/ethnicity | | Hispanic | | 0.0111 | | .00003 | | 0.0106 | | .00003 | | 21.02 | |
| Pallidum | | Race/ethnicity | | NH White | | 0.0110 | | .00003 | | 0.0106 | | .00003 | | 20.93 | |
| Pallidum | | Race/ethnicity | | NH Black/AA | | 0.0114 | | .00001 | | 0.0107 | | .00001 | | 20.43 | |
| Pallidum | | Race/ethnicity | | Unspecified | | 0.0111 | | .00004 | | 0.0106 | | .00004 | | 20.32 | |
| Pallidum | | Site | | A | | 0.0110 | | .00003 | | 0.0105 | | .00002 | | 21.45 | |
| Pallidum | | Site | | B | | 0.0112 | | .00002 | | 0.0108 | | .00002 | | 20.19 | |
| Pallidum | | Site | | C | | 0.0111 | | .00002 | | 0.0105 | | .00002 | | 19.98 | |
| Pallidum | | Site | | D | | 0.0111 | | .00002 | | 0.0106 | | .00002 | | 21.31 | |
| Pallidum | | Site | | E | | 0.0111 | | .00002 | | 0.0106 | | .00002 | | 21.22 | |
| Putamen | | Sex | | F | | 0.0145 | | .00001 | | 0.0139 | | .00001 | | 20.70 | |
| Putamen | | Sex | | M | | 0.0142 | | .00001 | | 0.0137 | | .00001 | | 20.48 | |
| Putamen | | Income | | <25k | | 0.0145 | | .00005 | | 0.0140 | | .00005 | | 20.34 | |
| Putamen | | Income | | 25-49k | | 0.0145 | | .00001 | | 0.0140 | | .00001 | | 20.42 | |
| Putamen | | Income | | 50-74k | | 0.0143 | | .00002 | | 0.0137 | | .00002 | | 20.28 | |
| Putamen | | Income | | 75-99k | | 0.0143 | | .00003 | | 0.0137 | | .00003 | | 19.82 | |
| Putamen | | Income | | 100k-199k | | 0.0143 | | .00003 | | 0.0138 | | .00002 | | 21.26 | |
| Putamen | | Income | | 200k+ | | 0.0144 | | .00002 | | 0.0139 | | .00002 | | 19.95 | |
| Putamen | | Race/ethnicity | | A, A, N | | 0.0144 | | .00004 | | 0.0138 | | .00004 | | 20.58 | |
| Putamen | | Race/ethnicity | | Hispanic | | 0.0145 | | .00002 | | 0.0140 | | .00002 | | 20.77 | |
| Putamen | | Race/ethnicity | | NH White | | 0.0143 | | .00003 | | 0.0138 | | .00003 | | 20.64 | |
| Putamen | | Race/ethnicity | | NH Black/AA | | 0.0146 | | .00001 | | 0.0139 | | .00001 | | 20.21 | |
| Putamen | | Race/ethnicity | | Unspecified | | 0.0143 | | .00004 | | 0.0138 | | .00003 | | 20.31 | |
| Putamen | | Site | | A | | 0.0143 | | .00002 | | 0.0138 | | .00002 | | 21.20 | |
| Putamen | | Site | | B | | 0.0144 | | .00002 | | 0.0139 | | .00002 | | 20.15 | |
| Putamen | | Site | | C | | 0.0144 | | .00002 | | 0.0138 | | .00002 | | 19.81 | |
| Putamen | | Site | | D | | 0.0144 | | .00002 | | 0.0138 | | .00001 | | 20.30 | |
| Putamen | | Site | | E | | 0.0143 | | .00002 | | 0.0138 | | .00001 | | 21.29 | |

**Supplementary Table S3. Full Sample: Developmental GAMM Model Statistics Across Individual Substances, Subscales, Trial Types, and Regions of Interest**

| **Substance Use: Statistics Across Individual Substances** | | | | | | | |
| --- | --- | --- | --- | --- | --- | --- | --- |
| **Substance** | **Population** | **s(age) F** | **s(age) p** | **Period of increase** | | **Period of decrease** | **Rate of Change** |
| All | Full Sample | 785.08 | <.001 | 12.00 – 25.05 | | 26.94 – 29.91 | .74 |
| Alcohol | Full Sample | 670.22 | <.001 | 15.51 – 25.32 | | 27.57 – 29.91 | .29 |
| Binge | Full Sample | 160.49 | <.001 | 13.80 – 23.07 | | 23.88 – 29.91 | .01 |
| Cannabis | Full Sample | 260.23 | <.001 | 12.00 – 24.78 | |  | .40 |
| Nicotine | Full Sample | 128.46 | <.001 | 12.00 – 24.06 | |  | .23 |
| **Impulsivity (UPPS): Statistics Across Individual Subscales** | | | | | | | |
| **Subscale** | **Population** | **s(age) F** | **s(age) p** | **Period of increase** | | **Period of decrease** | **Rate of Change** |
| Mean | Full Sample | -75.66 | <.001 | NA | | 12.00 – 29.30 | -.02 |
| Negative Urgency | Full Sample | -23.18 | <.001 | NA | | 12.00 – 25.74 | -.02 |
| Perseverance | Full Sample | -4.76 | <.001 | NA | | 15.30 – 22.08 | -.01 |
| Positive Urgency | Full Sample | -80.75 | <.001 | NA | | 12.00 – 26.95 | -.03 |
| Premeditation | Full Sample | -31.45 | <.001 | NA | | 12.00 – 22.52 | -.01 |
| Sensation seeking | Full Sample | -21.25 | <.001 | NA | | 18.26 – 29.30 | -.02 |
| **Inhibitory Control (Anti-Saccade): Statistics Across Individual Trial Types** | | | | | | | |
| **Trial Type** | **Population** | **s(age) F** | **s(age) p** | | **Period of increase** | **Period of decrease** | **Rate of Change** |
| Mean | Full Sample | 14.82 | < .001 | | 12.27 – 20.34 |  | 1.45 |
| Reward | Full Sample | 10.17 | < .001 | | 12.27 – 20.52 |  | 1.21 |
| Neutral | Full Sample | 14.29 | < .001 | | 12.27 – 20.63 |  | 1.57 |
| **Basal Ganglia Tissue Iron (nT2*w): Statistics Across Individual Regions of Interest** | | | | | | | |
| **Region of Interest** | **Population** | **s(age) F** | **s(age) p** | | **Period of increase** | **Period of decrease** | **Peak Change Age (years)** |
| Basal Ganglia | Full Sample | -253.25 | <.001 | | NA | 12.04 – 29.3 | 25.05 |
| Caudate | Full Sample | -76.48 | <.001 | | NA | 14.47 – 29.3 | 27.08 |
| NAcc | Full Sample | -42.32 | <.001 | | NA | 12.04 – 29.3 | 12.56 |
| Pallidum | Full Sample | -513.65 | <.001 | | NA | 12.04 – 29.3 | 13.12 |
| Putamen | Full Sample | -331.63 | <.001 | | NA | 12.04 – 29.3 | 24.35 |

*Note. All metrics result from independent generalized additive mixed models (GAMMs) described in the main text. F and p values describe the significance of the smooth term for age.*

**Supplementary Table S4. Sociodemographic Subgroups: Developmental GAMM Model Statistics Across Individual Substances, Subscales, Trial Types, and Regions of Interest**

| **Substance Use Model Statistics across Sociodemographic Factors** | | | | | | | |
| --- | --- | --- | --- | --- | --- | --- | --- |
| Substance | Population | Level | s(age) F | s(age) p | Period of increase | Period of decrease | Rate of Change |
| All | Sex | F | 361.58 | <.001 | 14.27 – 25.42 |  | 0.68 |
| All | Sex | M | 435.51 | <.001 | 12.00 – 24.36 | 26.78 – 29.83 | 0.82 |
| All | Income | <25k | 31.73 | <.001 | 12.04 – 28.23 |  | 1.24 |
| All | Income | 25 – 49k | 44.82 | <.001 | 12.14 – 29.90 |  | 1.09 |
| All | Income | 50 – 74k | 98.77 | <.001 | 15.66 – 24.16 |  | 0.67 |
| All | Income | 75 – 99k | 115.10 | <.001 | 13.53 – 24.42 |  | 0.92 |
| All | Income | 100k – 199k | 306.16 | <.001 | 12.02 – 24.51 | 26.80 – 29.53 | 0.70 |
| All | Income | 200k+ | 216.04 | <.001 | 14.31 – 24.42 |  | 0.96 |
| All | Race/ethnicity | A,A,N | 38.31 | <.001 | 12.00 – 24.19 |  | 0.66 |
| All | Race/ethnicity | Hispanic | 108.19 | <.001 | 13.50 – 24.05 |  | 0.89 |
| All | Race/ethnicity | NH White | 570.70 | <.001 | 12.02 – 24.70 |  | 0.73 |
| All | Race/ethnicity | NH Black/AA | 60.66 | <.001 | 14.48 – 28.78 | 26.67 – 29.91 | 1.05 |
| All | Race/ethnicity | Unspecified | 26.48 | <.001 | 14.64 – 25.83 |  | 1.08 |
| All | Site | A | 68.77 | <.001 | 14.66 – 25.92 |  | 0.81 |
| All | Site | B | 187.31 | <.001 | 14.35 – 23.48 |  | 0.74 |
| All | Site | C | 175.94 | <.001 | 15.09 – 24.34 |  | 0.65 |
| All | Site | D | 172.01 | <.001 | 13.58 – 23.97 | 26.70 – 29.70 | 0.68 |
| All | Site | E | 202.42 | <.001 | 12.00 – 24.91 |  | 0.89 |
| Alcohol | Sex | F | 325.55 | <.001 | 15.89 – 25.50 |  | 0.29 |
| Alcohol | Sex | M | 350.04 | <.001 | 15.58 – 24.90 | 29.02 – 29.83 | 0.33 |
| Alcohol | Income | <25k | 27.73 | <.001 | 16.84 – 28.23 |  | 0.37 |
| Alcohol | Income | 25 – 49k | 38.25 | <.001 | 16.87 – 24.72 |  | 0.22 |
| Alcohol | Income | 50 – 74k | 73.12 | <.001 | 16.36 – 25.30 |  | 0.35 |
| Alcohol | Income | 75-99K | 108.71 | <.001 | 16.05 – 25.32 |  | 0.43 |
| Alcohol | Income | 100k – 199k | 252.44 | <.001 | 15.72 – 24.25 | 25.92 – 29.53 | 0.24 |
| Alcohol | Income | 200k+ | 213.11 | <.001 | 15.43 – 29.05 |  | 0.67 |
| Alcohol | Race/ethnicity | A,A,N | 38.80 | <.001 | 16.21 – 24.19 |  | 0.23 |
| Alcohol | Race/ethnicity | Hispanic | 68.51 | <.001 | 16.05 – 24.14 |  | 0.28 |
| Alcohol | Race/ethnicity | NH White | 486.03 | <.001 | 15.53 – 24.97 |  | 0.32 |
| Alcohol | Race/ethnicity | NH Black/AA | 52.55 | <.001 | 16.58 – 28.78 | 27.57 – 29.91 | 0.39 |
| Alcohol | Race/ethnicity | Unspecified | 36.35 | <.001 | 16.28 – 26.40 |  | 0.58 |
| Alcohol | Site | A | 79.53 | <.001 | 16.44 – 25.39 |  | 0.32 |
| Alcohol | Site | B | 152.98 | <.001 | 15.39 – 24.43 |  | 0.42 |
| Alcohol | Site | C | 150.09 | <.001 | 16.08 – 25.95 |  | 0.37 |
| Alcohol | Site | D | 146.00 | <.001 | 16.22 – 24.24 | 27.41 – 29.70 | 0.30 |
| Alcohol | Site | E | 158.69 | <.001 | 16.03 – 24.91 |  | 0.28 |
| Binge | Sex | F | 63.03 | <.001 | 15.62 –  23.08 | 25.15 – 29.91 | 0.02 |
| Binge | Sex | M | 100.80 | <.001 | 13.70 – 22.57 | 23.56 – 29.83 | 0.01 |
| Binge | Income | <25k | 5.18 | <.001 | 15.95 – 22.45 |  | 0.06 |
| Binge | Income | 25 – 49k | 5.49 | <.001 | 15.89 – 23.30 |  | 0.07 |
| Binge | Income | 50 – 74k | 18.46 | <.001 | 16.36 – 22.41 |  | 0.05 |
| Binge | Income | 75-99K | 25.26 | <.001 | 15.87 – 22.71 |  | 0.05 |
| Binge | Income | 100k – 199k | 57.11 | <.001 | 13.96 – 22.67 | 23.90 – 29.53 | 0.02 |
| Binge | Income | 200k+ | 55.37 | <.001 | 15.26 – 21.94 | 24.25 – 29.05 | 0.03 |
| Binge | Race/ethnicity | A,A,N | 9.46 | <.001 | 15.32 – 22.04 |  | 0.05 |
| Binge | Race/ethnicity | Hispanic | 18.18 | <.001 | 16.22 – 22.12 | 25.46 – 29.51 | 0.01 |
| Binge | Race/ethnicity | NH White | 113.49 | <.001 | 12.47 – 22.81 | 23.80 – 29.91 | 0.02 |
| Binge | Race/ethnicity | NH Black/AA | 8.32 | <.001 | 16.83 – 21.88 |  | 0.03 |
| Binge | Race/ethnicity | Unspecified | 10.01 | <.001 | 16.12 – 23.36 |  | 0.22 |
| Binge | Site | A | -16.00 | <.001 | 16.61 – 22.29 | 24.15 – 29.91 | -0.02 |
| Binge | Site | B | 42.69 | <.001 | 15.04 – 21.74 | 24.17 – 29.30 | 0.02 |
| Binge | Site | C | 39.79 | <.001 | 15.99 – 22.36 | 24.96 – 29.90 | 0.01 |
| Binge | Site | D | 28.89 | <.001 | 15.34 – 22.74 | 28.91 – 29.70 | 0.05 |
| Binge | Site | E | 38.47 | <.001 | 15.41 – 22.94 | 25.81 – 29.84 | 0.04 |
| Cannabis | Sex | F | 116.57 | <.001 | 14.00 – 25.42 |  | 0.40 |
| Cannabis | Sex | M | 145.90 | <.001 | 12.00 – 23.83 |  | 0.42 |
| Cannabis | Income | <25k | 18.99 | <.001 | 12.04 – 23.43 |  | 0.81 |
| Cannabis | Income | 25 – 49k | 25.06 | <.001 | 12.14 – 25.26 |  | 0.73 |
| Cannabis | Income | 50 – 74k | 43.51 | <.001 | 15.74 – 23.90 |  | 0.46 |
| Cannabis | Income | 75-99K | 30.66 | <.001 | 12.00 – 26.49 |  | 0.63 |
| Cannabis | Income | 100k – 199k | 99.76 | <.001 | 12.02 – 24.60 |  | 0.40 |
| Cannabis | Income | 200k+ | 49.24 | <.001 | 15.68 – 22.20 | 25.54 – 29.05 | 0.12 |
| Cannabis | Race/Eth | A,A,N | 8.85 | <.001 | 14.24 – 23.65 |  | 0.25 |
| Cannabis | Race/ethnicity | Hispanic | 39.36 | <.001 | 12.00 – 24.58 |  | 0.68 |
| Cannabis | Race/ethnicity | NH White | 180.24 | <.001 | 12.92 – 24.34 |  | 0.37 |
| Cannabis | Race/ethnicity | NH Black/AA | 25.68 | <.001 | 12.04 – 25.50 |  | 0.62 |
| Cannabis | Race/ethnicity | Unspecified | 6.51 | <.001 | 15.71 – 21.38 |  | 0.34 |
| Cannabis | Site | A | 18.08 | <.001 | 12.27 – 29.91 |  | 0.45 |
| Cannabis | Site | B | 59.61 | <.001 | 15.22 – 22.95 |  | 0.36 |
| Cannabis | Site | C | 51.16 | <.001 | 15.81 – 23.80 |  | 0.32 |
| Cannabis | Site | D | 55.46 | <.001 | 12.17 – 23.53 |  | 0.40 |
| Cannabis | Site | E | 85.09 | <.001 | 12.00 – 24.10 |  | 0.53 |
| Nicotine | Sex | F | 43.40 | <.001 | 15.62 – 23.89 |  | 0.14 |
| Nicotine | Sex | M | 86.35 | <.001 | 12.00 – 23.47 |  | 0.33 |
| Nicotine | Income | <25k | 8.29 | <.001 | 14.97 – 24.16 |  | 0.50 |
| Nicotine | Income | 25 – 49k | 8.47 | <.001 | 14.73 – 26.95 |  | 0.34 |
| Nicotine | Income | 50 – 74k | 9.48 | <.001 | 16.27 – 22.32 |  | 0.18 |
| Nicotine | Income | 75-99K | 20.05 | <.001 | 12.00 – 23.34 |  | 0.40 |
| Nicotine | Income | 100k – 199k | 53.73 | <.001 | 12.55 – 23.46 |  | 0.20 |
| Nicotine | Income | 200k+ | 30.11 | <.001 | 12.77 – 22.80 |  | 0.31 |
| Nicotine | Race/ethnicity | A,A,N | 11.95 | <.001 | 12.00 – 20.07 |  | 0.26 |
| Nicotine | Race/ethnicity | Hispanic | 16.82 | <.001 | 13.58 – 22.38 |  | 0.26 |
| Nicotine | Race/ethnicity | NH White | 89.50 | <.001 | 12.02 – 23.71 |  | 0.23 |
| Nicotine | Race/ethnicity | NH Black/AA | 11.62 | <.001 | 16.75 – 25.50 |  | 0.35 |
| Nicotine | Race/ethnicity | Unspecified | 5.67 | <.001 | 16.45 – 24.92 |  | 0.51 |
| Nicotine | Site | A | 11.71 | <.001 | 15.64 – 25.30 |  | 0.31 |
| Nicotine | Site | B | 31.30 | <.001 | 15.39 – 22.17 |  | 0.16 |
| Nicotine | Site | C | 22.78 | <.001 | 15.00 – 23.35 |  | 0.24 |
| Nicotine | Site | D | 24.17 | <.001 | 12.17 – 22.83 |  | 0.27 |
| Nicotine | Site | E | 36.86 | <.001 | 12.00 – 23.47 |  | 0.30 |
| **Impulsivity (UPPS) Model Statistics across Sociodemographic Factors** | | | | | | | |
| **Subscale** | **Population** | **Level** | **S(age) F** | **s(age)p** | **Period of increase** | **Period of decrease** | **Rate of change** |
| Mean | Sex | F | -60.34 | <.001 |  | 12.02 - 25.22 | -0.021 |
| Mean | Sex | M | -20.13 | <.001 |  | 14.91 - 29.05 | -0.015 |
| Mean | Income | <25k | -7.11 | <.001 |  | 12.04 - 18.31 | -0.021 |
| Mean | Income | 25k-49k | -9.80 | <.001 |  | 12.14 - 20.88 | -0.019 |
| Mean | Income | 50k-74k | -22.60 | <.001 |  | 12.06 - 25.39 | -0.029 |
| Mean | Income | 75k-99k | -3.74 | <.001 |  | 17.60 - 23.11 | -0.012 |
| Mean | Income | 100k-199k | -25.67 | <.001 |  | 14.97 - 29.30 | -0.017 |
| Mean | Income | 200k+ | -14.41 | <.001 |  | 15.94- 25.62 | -0.018 |
| Mean | Race/ethnicity | A, A, N | -8.01 | <.001 |  | 16.13 - 27.63 | -0.020 |
| Mean | Race/ethnicity | Hispanic | -18.09 | <.001 |  | 12.00 - 27.72 | -0.027 |
| Mean | Race/ethnicity | NH White | -42.48 | <.001 |  | 16.01 - 25.31 | -0.014 |
| Mean | Race/ethnicity | NH Black/AA | -13.96 | <.001 |  | 12.04 - 19.61 | -0.023 |
| Mean | Race/ethnicity | Other | -1.27 | 0.03 |  |  | -0.009 |
| Mean | Site | A | -9.80 | <.001 |  | 12.27 - 22.35 | -0.015 |
| Mean | Site | B | -8.36 | <.001 |  | 15.39 - 25.91 | -0.016 |
| Mean | Site | C | -7.02 | <.001 |  | 12.04 - 18.71 | -0.014 |
| Mean | Site | D | -20.29 | <.001 |  | 16.57 - 24.63 | -0.019 |
| Mean | Site | E | -41.05 | <.001 |  | 12.00 - 29.05 | -0.025 |
| Negative Urgency | Sex | F | -13.94 | <.001 |  | 16.10 - 24.87 | -0.022 |
| Negative Urgency | Sex | M | -10.73 | <.001 |  | 12.00 - 22.62 | -0.020 |
| Negative Urgency | Income | <25k | -2.18 | 0.01 |  |  | -0.020 |
| Negative Urgency | Income | 25k-49k | -3.25 | <.001 |  | 15.46 - 18.86 | -0.024 |
| Negative Urgency | Income | 50k-74k | -5.25 | <.001 |  | 15.81 - 22.81 | -0.028 |
| Negative Urgency | Income | 75k-99k | -1.36 | 0.03 |  |  | -0.013 |
| Negative Urgency | Income | 100k-199k | -5.28 | <.001 |  | 16.88 - 24.96 | -0.016 |
| Negative Urgency | Income | 200k+ | -8.11 | <.001 |  | 15.77 - 25.71 | -0.030 |
| Negative Urgency | Race/ethnicity | A, A, N | -2.51 | 0.01 |  | 20.51 - 22.77 | -0.025 |
| Negative Urgency | Race/ethnicity | Hispanic | -5.10 | 0.01 |  | 15.68 - 23.29 | -0.027 |
| Negative Urgency | Race/ethnicity | NH White | -10.61 | <.001 |  | 15.32 - 24.35 | -0.017 |
| Negative Urgency | Race/ethnicity | NH Black/AA | -7.26 | <.001 |  | 12.04 - 21.40 | -0.037 |
| Negative Urgency | Race/ethnicity | Other | 0.00 | 0.48 |  |  | 0.000 |
| Negative Urgency | Site | A | -3.93 | <.001 |  | 16.19 -20.93 | -0.019 |
| Negative Urgency | Site | B | -1.62 | 0.02 |  |  | -0.013 |
| Negative Urgency | Site | C | -3.14 | <.001 |  | 15.37 - 17.99 | -0.018 |
| Negative Urgency | Site | D | -5.31 | <.001 |  | 17.79 - 22.35 | -0.014 |
| Negative Urgency | Site | E | -15.09 | <.001 |  | 14.91 - 28.28 | -0.032 |
| Perseverance | Sex | F | -5.06 | 0.01 |  | 15.41 - 21.22 | -0.008 |
| Perseverance | Sex | M | -0.83 | 0.06 |  |  | -0.004 |
| Perseverance | Income | <25k | -0.75 | 0.08 |  |  | -0.012 |
| Perseverance | Income | 25k-49k | -2.31 | 0.01 |  | 16.99 - 18.05 | -0.015 |
| Perseverance | Income | 50k-74k | -0.76 | 0.07 |  |  | -0.007 |
| Perseverance | Income | 75k-99k | 0.00 | 0.68 |  |  | 0.000 |
| Perseverance | Income | 100k-199k | -1.66 | 0.02 |  |  | -0.005 |
| Perseverance | Income | 200k+ | 0.00 | 0.55 |  |  | 0.000 |
| Perseverance | Race/ethnicity | A, A, N | 0.00 | 0.96 |  |  | 0.000 |
| Perseverance | Race/ethnicity | Hispanic | -3.55 | <.001 |  | 15.85 - 22.45 | -0.017 |
| Perseverance | Race/ethnicity | NH White | -2.08 | 0.01 |  |  | -0.005 |
| Perseverance | Race/ethnicity | NH Black/AA | -1.99 | 0.01 |  |  | -0.012 |
| Perseverance | Race/ethnicity | Other | 0.03 | 0.30 |  |  | 0.001 |
| Perseverance | Site | A | .00001 | 0.73 |  |  | .00001 |
| Perseverance | Site | B | 0.00 | 0.74 |  |  | 0.000 |
| Perseverance | Site | C | -0.27 | 0.18 |  |  | -0.003 |
| Perseverance | Site | D | -0.12 | 0.24 |  |  | -0.001 |
| Perseverance | Site | E | -7.91 | <.001 |  | 12.00 - 22.88 | -0.016 |
| Positive Urgency | Sex | F | -51.43 | <.001 |  | 12.02 - 25.48 | -0.035 |
| Positive Urgency | Sex | M | -31.18 | <.001 |  | 13.88 - 26.14 | -0.031 |
| Positive Urgency | Income | <25k | -6.56 | <.001 |  | 12.04 - 18.23 | -0.043 |
| Positive Urgency | Income | 25k-49k | -6.81 | <.001 |  | 12.14 - 21.36 | -0.036 |
| Positive Urgency | Income | 50k-74k | -23.39 | <.001 |  | 14.23 - 23.47 | -0.046 |
| Positive Urgency | Income | 75k-99k | -7.64 | <.001 |  | 17.10 - 22.70 | -0.023 |
| Positive Urgency | Income | 100k-199k | -26.20 | <.001 |  | 12.02 - 27.30 | -0.031 |
| Positive Urgency | Income | 200k+ | -17.29 | <.001 |  | 15.60 - 26.05 | -0.033 |
| Positive Urgency | Race/ethnicity | A, A, N | -7.37 | <.001 |  | 14.51 - 23.34 | -0.039 |
| Positive Urgency | Race/ethnicity | Hispanic | -16.55 | <.001 |  | 12.00 - 22.03 | -0.043 |
| Positive Urgency | Race/ethnicity | NH White | -55.03 | <.001 |  | 15.84 - 24.78 | -0.025 |
| Positive Urgency | Race/ethnicity | NH Black/AA | -10.17 | <.001 |  | 12.04 - 20.83 | -0.044 |
| Positive Urgency | Race/ethnicity | Other | -0.91 | <.001 |  |  | -0.014 |
| Positive Urgency | Site | A | -10.07 | <.001 |  | 12.27 - 23.77 | -0.033 |
| Positive Urgency | Site | B | -11.90 | <.001 |  | 14.26 - 25.65 | -0.033 |
| Positive Urgency | Site | C | -12.45 | <.001 |  | 12.04 - 18.55 | -0.037 |
| Positive Urgency | Site | D | -24.85 | <.001 |  | 16.49 - 22.68 | -0.021 |
| Positive Urgency | Site | E | -30.61 | <.001 |  | 12.00 -26.14 | -0.037 |
| Premeditation | Sex | F | -26.78 | <.001 |  | 12.02 - 22.01 | -0.017 |
| Premeditation | Sex | M | -6.74 | <.001 |  | 13.11 - 22.20 | -0.011 |
| Premeditation | Income | <25k | -2.16 | 0.01 |  |  | -0.021 |
| Premeditation | Income | 25k-49k | -5.60 | <.001 |  | 14.65 - 20.47 | -0.023 |
| Premeditation | Income | 50k-74k | -14.18 | <.001 |  | 12.06 - 23.72 | -0.034 |
| Premeditation | Income | 75k-99k | 0.00 | .37 |  |  | 0.000 |
| Premeditation | Income | 100k-199k | -7.48 | <.001 |  | 12.02 - 22.01 | -0.012 |
| Premeditation | Income | 200k+ | -5.12 | <.001 |  | 15.17- 20.48 | -0.013 |
| Premeditation | Race/ethnicity | A, A, N | -1.72 | .01 |  |  | -0.013 |
| Premeditation | Race/ethnicity | Hispanic | -4.12 | <.001 |  | 12.00 -18.69 | -0.016 |
| Premeditation | Race/ethnicity | NH White | -16.83 | <.001 |  | 12.02 - 22.09 | -0.013 |
| Premeditation | Race/ethnicity | NH Black/AA | -5.07 | <.001 |  | 13.67 - 19.86 | -0.019 |
| Premeditation | Race/ethnicity | Other | -1.70 | 0.02 |  |  | -0.017 |
| Premeditation | Site | A | -3.84 | <.001 |  | 12.77 - 18.77 | -0.012 |
| Premeditation | Site | B | -5.05 | <.001 |  | 15.22 - 22.08 | -0.016 |
| Premeditation | Site | C | -1.25 | 0.03 |  |  | -0.008 |
| Premeditation | Site | D | -7.04 | <.001 |  | 14.21 - 22.27 | -0.020 |
| Premeditation | Site | E | -14.90 | <.001 |  | 12.00 - 21.85 | -0.019 |
| Sensation Seeking | Sex | F | -24.64 | <.001 |  | 15.06 - 29.30 | -0.032 |
| Sensation Seeking | Sex | M | -13.02 | <.001 |  | 20.91 - 29.05 | -0.007 |
| Sensation Seeking | Income | <25k | 0.00 | 0.82 |  |  | 0.000 |
| Sensation Seeking | Income | 25k-49k | 0.00 | 0.43 |  |  | 0.000 |
| Sensation Seeking | Income | 50k-74k | -2.70 | <.001 |  | 16.89 - 17.48 | -0.019 |
| Sensation Seeking | Income | 75k-99k | -7.82 | <.001 |  | 20.97 - 28.38 | -0.033 |
| Sensation Seeking | Income | 100k-199k | -17.66 | <.001 |  | 18.36 - 29.30 | -0.019 |
| Sensation Seeking | Income | 200k+ | -2.08 | 0.02 |  |  | -0.011 |
| Sensation Seeking | Race/ethnicity | A, A, N | -4.27 | <.001 |  | 20.83 - 28.12 | -0.027 |
| Sensation Seeking | Race/ethnicity | Hispanic | -6.27 | <.001 |  | 18.44 - 28.64 | -0.030 |
| Sensation Seeking | Race/ethnicity | NH White | -18.46 | <.001 |  | 18.53 - 29.30 | -0.016 |
| Sensation Seeking | Race/ethnicity | NH Black/AA | 0.00 | 0.78 |  |  | 0.000 |
| Sensation Seeking | Race/ethnicity | Other | -0.51 | 0.12 |  |  | -0.009 |
| Sensation Seeking | Site | A | -2.42 | 0.01 |  | 20.68 -22.93 | -0.016 |
| Sensation Seeking | Site | B | -0.77 | 0.09 |  |  | -0.008 |
| Sensation Seeking | Site | C | 0.00 | 0.71 |  |  | 0.000 |
| Sensation Seeking | Site | D | -10.30 | <.001 |  | 19.01 - 28.38 | -0.028 |
| Sensation Seeking | Site | E | -11.69 | <.001 |  | 18.34 - 29.05 | -0.02 |
| **Inhibitory Control (Anti-Saccade) Model Statistics across Sociodemographic Factors** | | | | | | | |
| **Trial Type** | **Population** | **Level** | **s(age) F** | **s(age) p** | **Period of increase** | **Period of decrease** | **Rate of Change** |
| Mean | Sex | F | 2.45 | <.001 |  |  | 0.81 |
| Mean | Sex | M | 8.49 | <.001 | 12.27 – 19.90 |  | 1.69 |
| Mean | Income | <25k | 1.59 | 0.02 |  |  | 1.99 |
| Mean | Income | 25 – 49k | 0.75 | 0.08 |  |  | 0.78 |
| Mean | Income | 50 – 74k | 0.89 | 0.07 |  |  | 1.16 |
| Mean | Income | 75 – 99k | 2.36 | 0.01 | 15.61 – 16.23 |  | 1.11 |
| Mean | Income | 100k – 199k | 2.95 | <.001 | 14.72 -17.51 |  | 0.98 |
| Mean | Income | 200k+ | 1.65 | 0.02 |  |  | 1.54 |
| Mean | Race/ethnicity | A,A,N | 0.00 | 0.97 |  |  | 0.00 |
| Mean | Race/ethnicity | Hispanic | 0.75 | 0.11 |  |  | 1.49 |
| Mean | Race/ethnicity | NH White | 7.56 | <.001 | 12.27 - 19.80 |  | 1.27 |
| Mean | Race/ethnicity | NH Black/AA | 2.08 | 0.01 |  |  | 1.18 |
| Mean | Race/ethnicity | Unspecified | 0.98 | 0.07 |  |  | 2.08 |
| Mean | Site | A | 14.45 | <.001 | 12.27 – 19.98 |  | 1.71 |
| Mean | Site | C | 1.99 | 0.01 |  |  | 0.82 |
| Reward | Sex | F | 1.39 | 0.03 |  |  | 0.56 |
| Reward | Sex | M | 5.50 | <.001 | 14.27 – 20.06 |  | 1.41 |
| Reward | Income | <25k | 1.17 | 0.05 |  |  | 1.82 |
| Reward | Income | 25 – 49k | 0.61 | 0.10 |  |  | 0.67 |
| Reward | Income | 50 – 74k | 0.58 | 0.11 |  |  | 0.89 |
| Reward | Income | 75 – 99k | 1.51 | 0.02 |  |  | 0.95 |
| Reward | Income | 100k – 199k | 1.38 | 0.03 |  |  | 0.63 |
| Reward | Income | 200k+ | 1.07 | 0.05 |  |  | 0.98 |
| Reward | Race/ethnicity | A,A,N | 0.00 | 0.65 |  |  | 0.00 |
| Reward | Race/ethnicity | Hispanic | 0.94 | 0.08 |  |  | 1.48 |
| Reward | Race/ethnicity | NH White | 3.50 | <.001 | 14.24 – 18.78 |  | 0.84 |
| Reward | Race/ethnicity | NH Black/AA | 2.85 | <.001 | 17.13 – 19.41 |  | 1.40 |
| Reward | Race/ethnicity | Unspecified | 0.82 | 0.08 |  |  | 1.94 |
| Reward | Site | A | 8.55 | <.001 | 12.27 – 19.56 |  | 1.36 |
| Reward | Site | C | 1.64 | 0.02 |  |  | 0.71 |
| Neutral | Sex | F | 2.48 | <.001 | 16.09 – 16.80 |  | 0.90 |
| Neutral | Sex | M | 8.04 | <.001 | 13.05 – 19.40 |  | 1.70 |
| Neutral | Income | <25k | 1.71 | 0.02 |  |  | 2.03 |
| Neutral | Income | 25 – 49k | 0.71 | 0.08 |  |  | 0.80 |
| Neutral | Income | 50 – 74k | 1.05 | 0.05 |  |  | 1.37 |
| Neutral | Income | 75 – 99k | 2.15 | 0.01 |  |  | 1.01 |
| Neutral | Income | 100k – 199k | 2.54 | <.001 | 15.34 - 17.17 | 17.17 | 1.05 |
| Neutral | Income | 200k+ | 2.26 | 0.01 |  |  | 1.99 |
| Neutral | Race/ethnicity | A,A,N | .00001 | 0.84 |  |  | .00001 |
| Neutral | Race/ethnicity | Hispanic | 0.48 | 0.16 |  |  | 1.34 |
| Neutral | Race/ethnicity | NH White | 7.54 | <.001 | 12.27 – 20.28 | 20.28 | 1.42 |
| Neutral | Race/ethnicity | NH Black/AA | 1.02 | 0.05 |  |  | 0.85 |
| Neutral | Race/ethnicity | Unspecified | 1.98 | 0.02 |  |  | 2.73 |
| Neutral | Site | A | 13.93 | <.001 | 12.27 – 20.22 |  | 1.91 |
| Neutral | Site | C | 1.73 | 0.01 |  |  | 0.81 |
| **Tissue Iron (nT2*w) Model Statistics across Sociodemographic Factors** | | | | | | | |
| **Region of Interest** | **Population** | **Level** | **S(age) F** | **s(age)p** | **Period of increase** | **Period of decrease** | **Peak Change Age (years)** |
| Basal Ganglia | Sex | F | -131.37 | <.001 |  | 12.04 – 29.3 | 23.74 |
| Basal Ganglia | Sex | M | -122.32 | <.001 |  | 12.05 – 29.05 | 27.04 |
| Basal Ganglia | Income | <25k | -16.46 | <.001 |  | 15.01 – 24.93 | 20.68 |
| Basal Ganglia | Income | 25k-49k | -32.62 | <.001 |  | 12.14 – 28.18 | 26.44 |
| Basal Ganglia | Income | 50k-74k | -28.56 | <.001 |  | 12.06 – 28.64 | 13.76 |
| Basal Ganglia | Income | 75k-99k | -21.83 | <.001 |  | 14.54 – 28.35 | 24.34 |
| Basal Ganglia | Income | 100k-199k | -118.60 | <.001 |  | 12.27 – 29.3 | 24.07 |
| Basal Ganglia | Income | 200k+ | -37.02 | <.001 |  | 14.35 – 29.05 | 20.37 |
| Basal Ganglia | Race/ethnicity | A, A, N | -19.38 | <.001 |  | 12.78 – 24.88 | 16.78 |
| Basal Ganglia | Race/ethnicity | Hispanic | -52.34 | <.001 |  | 12.09 – 28.64 | 25.68 |
| Basal Ganglia | Race/ethnicity | Non-Hispanic Black/AA | -25.26 | <.001 |  | 14.61 – 28.03 | 25.78 |
| Basal Ganglia | Race/ethnicity | Non-Hispanic White | -145.83 | <.001 |  | 12.05 – 29.30 | 26.69 |
| Basal Ganglia | Race/ethnicity | Other | -15.59 | <.001 |  | 15.62 – 27.11 | 25.45 |
| Basal Ganglia | Site | A | -24.63 | <.001 |  | 13.80 – 28.35 | 25.19 |
| Basal Ganglia | Site | B | -68.20 | <.001 |  | 13.61 – 29.3 | 19.50 |
| Basal Ganglia | Site | C | -41.93 | <.001 |  | 12.04 – 28.03 | 12.56 |
| Basal Ganglia | Site | D | -24.06 | <.001 |  | 17.12 – 28.3 | 26.03 |
| Basal Ganglia | Site | E | -126.76 | <.001 |  | 12.09 – 29.05 | 15.96 |
| Caudate | Sex | F | -42.65 | <.001 |  | 15.85 – 29.30 | 28.04 |
| Caudate | Sex | M | -33.77 | <.001 |  | 13.07 – 29.05 | 25.16 |
| Caudate | Income | <25k | -5.74 | <.001 |  | 16.31 – 22.98 | 20.33 |
| Caudate | Income | 25k-49k | -12.78 | <.001 |  | 16.00 – 28.18 | 27.45 |
| Caudate | Income | 50k-74k | -6.24 | <.001 |  | 16.05 – 25.97 | 24.68 |
| Caudate | Income | 75k-99k | -3.99 | <.001 |  | 17.57 – 23.36 | 22.83 |
| Caudate | Income | 100k-199k | -35.74 | <.001 |  | 12.27 – 29.30 | 26.34 |
| Caudate | Income | 200k+ | -14.17 | <.001 |  | 16.49 – 29.05 | 28.53 |
| Caudate | Race/ethnicity | A, A, N | -5.34 | <.001 |  | 15.53 – 22.99 | 17.61 |
| Caudate | Race/ethnicity | Hispanic | -20.09 | <.001 |  | 15.83 – 28.64 | 26.97 |
| Caudate | Race/ethnicity | Non-Hispanic Black/AA | -8.07 | <.001 |  | 17.02 – 28.03 | 26.54 |
| Caudate | Race/ethnicity | Non-Hispanic White | -39.84 | <.001 |  | 14.56 – 29.30 | 26.74 |
| Caudate | Race/ethnicity | Other | -6.03 | <.001 |  | 17.95 – 24.63 | 23.28 |
| Caudate | Site | A | -7.78 | <.001 |  | 16.55 – 28.35 | 26.81 |
| Caudate | Site | B | -38.26 | <.001 |  | 15.95 – 29.30 | 26.74 |
| Caudate | Site | C | -10.92 | <.001 |  | 13.48 – 25.37 | 19.39 |
| Caudate | Site | D | -3.35 | .005 |  | 20.36 – 22.38 | 21.33 |
| Caudate | Site | E | -36.27 | <.001 |  | 12.09 – 29.05 | 20.72 |
| NAcc | Sex | F | -14.80 | <.001 |  | 12.64 – 29.30 | 28.38 |
| NAcc | Sex | M | -32.84 | <.001 |  | 12.05 – 25.37 | 12.56 |
| NAcc | Income | <25k | -0.81 | 0.068 |  | NA | NA |
| NAcc | Income | 25k-49k | -6.05 | <.001 |  | 20.20 – 28.18 | 27.17 23 |
| NAcc | Income | 50k-74k | -5.02 | <.001 |  | 12.06 – 18.39 | 12.80 |
| NAcc | Income | 75k-99k | -7.25 | <.001 |  | 12.10 – 18.22 | 12.67 |
| NAcc | Income | 100k-199k | -22.93 | <.001 |  | 12.27 – 29.04 | 13.25 |
| NAcc | Income | 200k+ | -2.55 | .003 |  | 16.74 – 20.67 | 17.43 |
| NAcc | Race/ethnicity | A, A, N | -6.36 | <.001 |  | 12.78 – 19.85 | 13.56 |
| NAcc | Race/ethnicity | Hispanic | -7.85 | <.001 |  | 15.66 – 28.64 | 28.09 |
| NAcc | Race/ethnicity | Non-Hispanic Black/AA | -4.89 | <.001 |  | 15.49 – 23.04 | 19.23 |
| NAcc | Race/ethnicity | Non-Hispanic White | -22.03 | <.001 |  | 12.05 – 25.48 | 12.70 |
| NAcc | Race/ethnicity | Other | -3.99 | <.001 |  | 16.37 – 22.53 | 21.85 |
| NAcc | Site | A | -5.48 | <.001 |  | 12.27 – 23.25 | 13.23 |
| NAcc | Site | B | -13.43 | <.001 |  | 13.78 – 27.30 | 22.32 |
| NAcc | Site | C | -7.47 | <.001 |  | 12.04 – 18.14 | 12.44 |
| NAcc | Site | D | -3.10 | .003 |  | 14.93 – 18.17 | 15.13 |
| NAcc | Site | E | -14.78 | <.001 |  | 12.09 – 25.72 | 13.36 |
| Pallidum | Sex | F | -268.73 | <.001 |  | 12.04 – 29.30 | 12.77 |
| Pallidum | Sex | M | -250.55 | <.001 |  | 12.05 -29.05 | 14.74 |
| Pallidum | Income | <25k | -30.63 | <.001 |  | 12.04 -24.02 | 13.15 |
| Pallidum | Income | 25k-49k | -68.57 | <.001 |  | 12.14 – 24.55 | 12.98 |
| Pallidum | Income | 50k-74k | -55.67 | <.001 |  | 12.06 – 28.64 | 12.55 |
| Pallidum | Income | 75k-99k | -54.40 | <.001 |  | 12.10 – 25.90 | 18.91 |
| Pallidum | Income | 100k-199k | -203.87 | <.001 |  | 12.27 – 29.30 | 14.06 |
| Pallidum | Income | 200k+ | -109.35 | <.001 |  | 12.05 –  29.05 | 15.97 |
| Pallidum | Race/ethnicity | A, A, N | -36.739 | <.001 |  | 12.78 – 24.88 | 14.35 |
| Pallidum | Race/ethnicity | Hispanic | -107.03 | <.001 |  | 12.09 – 28.64 | 12.88 |
| Pallidum | Race/ethnicity | Non-Hispanic Black/AA | -45.59 | <.001 |  | 12.04 – 28.03 | 13.80 |
| Pallidum | Race/ethnicity | Non-Hispanic White | -311.94 | <.001 |  | 12.05 – 29.3 | 15.17 |
| Pallidum | Race/ethnicity | Other | -50.07 | <.001 |  | 13.52 – 22.98 | 18.10 |
| Pallidum | Site | A | -46.23 | <.001 |  | 12.27 – 28.35 | 19.38 |
| Pallidum | Site | B | -150.03 | <.001 |  | 12.05 – 25.39 | 16.77 |
| Pallidum | Site | C | -67.02 | <.001 |  | 12.04 – 23.85 | 12.96 |
| Pallidum | Site | D | -112.31 | <.001 |  | 12.18 – 28.30 | 19.95 |
| Pallidum | Site | E | -221.95 | <.001 |  | 12.09 – 29.05 | 13.75 |
| Putamen | Sex | F | -156.05 | <.001 |  | 12.04 – 29.30 | 23.83 |
| Putamen | Sex | M | -178.64 | <.001 |  | 12.05 – 29.05 | 25.24 |
| Putamen | Income | <25k | -19.58 | <.001 |  | 14.95 – 24.93 | 23.47 |
| Putamen | Income | 25k-49k | -34.80 | <.001 |  | 12.14 – 28.18 | 13.42 |
| Putamen | Income | 50k-74k | -39.12 | <.001 |  | 12.06 – 28.64 | 13.97 |
| Putamen | Income | 75k-99k | -35.54 | <.001 |  | 15.77 – 28.35 | 25.73 |
| Putamen | Income | 100k-199k | -159.33 | <.001 |  | 12.27 – 29.30 | 23.73 |
| Putamen | Income | 200k+ | -45.97 | <.001 |  | 14.27 – 28.96 | 19.39 |
| Putamen | Race/ethnicity | A, A, N | -19.47 | <.001 |  | 12.78 – 26.21 | 19.10 |
| Putamen | Race/ethnicity | Hispanic | -65.51 | <.001 |  | 12.09 – 28.64 | 27.60 |
| Putamen | Race/ethnicity | Non-Hispanic Black/AA | -26.43 | <.001 |  | 14.93 – 28.03 | 26.26 |
| Putamen | Race/ethnicity | Non-Hispanic White | -210.30 | <.001 |  | 12.05 – 29.30 | 24.74 |
| Putamen | Race/ethnicity | Other | -20.14 | <.001 |  | 15.47 – 27.11 | 22.71 |
| Putamen | Site | A | -31.66 | <.001 |  | 14.77 – 28.35 | 24.91 |
| Putamen | Site | B | -78.14 | <.001 |  | 14.13 – 29.30 | 19.15 |
| Putamen | Site | C | -50.85 | <.001 |  | 12.04 – 28.03 | 20.80 |
| Putamen | Site | D | -40.20 | <.001 |  | 16.87 – 28.30 | 26.72 |
| Putamen | Site | E | -164.63 | <.001 |  | 12.09 – 29.05 | 18.05 |
| NAcc | Site | A | -5.48 | <.001 |  | 12.27 – 23.25 | 13.23 |
| NAcc | Site | B | -13.43 | <.001 |  | 13.78 – 27.30 | 22.32 |
| NAcc | Site | C | -7.47 | <.001 |  | 12.04 – 18.14 | 12.44 |
| NAcc | Site | D | -3.10 | .003 |  | 14.93 – 18.17 | 15.13 |
| NAcc | Site | E | -14.78 | <.001 |  | 12.09 – 25.72 | 13.36 |
| Pallidum | Sex | F | -268.73 | <.001 |  | 12.04 – 29.30 | 12.77 |
| Pallidum | Sex | M | -250.55 | <.001 |  | 12.05 -29.05 | 14.74 |
| Pallidum | Income | <25k | -30.63 | <.001 |  | 12.04 -24.02 | 13.15 |
| Pallidum | Income | 25k-49k | -68.57 | <.001 |  | 12.14 – 24.55 | 12.98 |
| Pallidum | Income | 50k-74k | -55.67 | <.001 |  | 12.06 – 28.64 | 12.55 |
| Pallidum | Income | 75k-99k | -54.40 | <.001 |  | 12.10 – 25.90 | 18.91 |
| Pallidum | Income | 100k-199k | -203.87 | <.001 |  | 12.27 – 29.30 | 14.06 |
| Pallidum | Income | 200k+ | -109.35 | <.001 |  | 12.05 –  29.05 | 15.97 |
| Pallidum | Race/ethnicity | A, A, N | -36.739 | <.001 |  | 12.78 – 24.88 | 14.35 |
| Pallidum | Race/ethnicity | Hispanic | -107.03 | <.001 |  | 12.09 – 28.64 | 12.88 |
| Pallidum | Race/ethnicity | Non-Hispanic Black/AA | -45.59 | <.001 |  | 12.04 – 28.03 | 13.80 |
| Pallidum | Race/ethnicity | Non-Hispanic White | -311.94 | <.001 |  | 12.05 – 29.3 | 15.17 |
| Pallidum | Race/ethnicity | Other | -50.07 | <.001 |  | 13.52 – 22.98 | 18.10 |
| Pallidum | Site | A | -46.23 | <.001 |  | 12.27 – 28.35 | 19.38 |
| Pallidum | Site | B | -150.03 | <.001 |  | 12.05 – 25.39 | 16.77 |
| Pallidum | Site | C | -67.02 | <.001 |  | 12.04 – 23.85 | 12.96 |
| Pallidum | Site | D | -112.31 | <.001 |  | 12.18 – 28.30 | 19.95 |
| Pallidum | Site | E | -221.95 | <.001 |  | 12.09 – 29.05 | 13.75 |
| Putamen | Sex | F | -156.05 | <.001 |  | 12.04 – 29.30 | 23.83 |
| Putamen | Sex | M | -178.64 | <.001 |  | 12.05 – 29.05 | 25.24 |
| Putamen | Income | <25k | -19.58 | <.001 |  | 14.95 – 24.93 | 23.47 |
| Putamen | Income | 25k-49k | -34.80 | <.001 |  | 12.14 – 28.18 | 13.42 |
| Putamen | Income | 50k-74k | -39.12 | <.001 |  | 12.06 – 28.64 | 13.97 |
| Putamen | Income | 75k-99k | -35.54 | <.001 |  | 15.77 – 28.35 | 25.73 |
| Putamen | Income | 100k-199k | -159.33 | <.001 |  | 12.27 – 29.30 | 23.73 |
| Putamen | Income | 200k+ | -45.97 | <.001 |  | 14.27 – 28.96 | 19.39 |
| Putamen | Race/ethnicity | A, A, N | -19.47 | <.001 |  | 12.78 – 26.21 | 19.10 |
| Putamen | Race/ethnicity | Hispanic | -65.51 | <.001 |  | 12.09 – 28.64 | 27.60 |
| Putamen | Race/ethnicity | Non-Hispanic Black/AA | -26.43 | <.001 |  | 14.93 – 28.03 | 26.26 |
| Putamen | Race/ethnicity | Non-Hispanic White | -210.30 | <.001 |  | 12.05 – 29.30 | 24.74 |
| Putamen | Race/ethnicity | Other | -20.14 | <.001 |  | 15.47 – 27.11 | 22.71 |
| Putamen | Site | A | -31.66 | <.001 |  | 14.77 – 28.35 | 24.91 |
| Putamen | Site | B | -78.14 | <.001 |  | 14.13 – 29.30 | 19.15 |
| Putamen | Site | C | -50.85 | <.001 |  | 12.04 – 28.03 | 20.80 |
| Putamen | Site | D | -40.20 | <.001 |  | 16.87 – 28.30 | 26.72 |
| Putamen | Site | E | -164.63 | <.001 |  | 12.09 – 29.05 | 18.05 |
| Putamen | Income | <25k | -19.58 | <.001 |  | 14.95 – 24.93 | 23.47 |
| Putamen | Income | 25k-49k | -34.80 | <.001 |  | 12.14 – 28.18 | 13.42 |
| Putamen | Income | 50k-74k | -39.12 | <.001 |  | 12.06 – 28.64 | 13.97 |
| Putamen | Income | 75k-99k | -35.54 | <.001 |  | 15.77 – 28.35 | 25.73 |
| Putamen | Income | 100k-199k | -159.33 | <.001 |  | 12.27 – 29.30 | 23.73 |
| Putamen | Income | 200k+ | -45.97 | <.001 |  | 14.27 – 28.96 | 19.39 |
| Putamen | Race/ethnicity | A, A, N | -19.47 | <.001 |  | 12.78 – 26.21 | 19.10 |
| Putamen | Race/ethnicity | Hispanic | -65.51 | <.001 |  | 12.09 – 28.64 | 27.60 |
| Putamen | Race/ethnicity | Non-Hispanic Black/AA | -26.43 | <.001 |  | 14.93 – 28.03 | 26.26 |
| Putamen | Race/ethnicity | Non-Hispanic White | -210.30 | <.001 |  | 12.05 – 29.30 | 24.74 |
| Putamen | Race/ethnicity | Other | -20.14 | <.001 |  | 15.47 – 27.11 | 22.71 |
| Putamen | Site | A | -31.66 | <.001 |  | 14.77 – 28.35 | 24.91 |
| Putamen | Site | B | -78.14 | <.001 |  | 14.13 – 29.30 | 19.15 |
| Putamen | Site | C | -50.85 | <.001 |  | 12.04 – 28.03 | 20.80 |
| Putamen | Site | D | -40.20 | <.001 |  | 16.87 – 28.30 | 26.72 |
| Putamen | Site | E | -164.63 | <.001 |  | 12.09 – 29.05 | 18.05 |

*Note. All metrics result from independent generalized additive mixed models (GAMMs) described in the main text. F and p values describe the significance of the smooth term for age.*

**Supplementary Table S5. Sociodemographic Subgroups: Differences in the Magnitude of Substance use, Impulsivity, Inhibitory Control, and Tissue Iron across Individual Substances, Subscales, Trial Types, and Regions of Interest**

| **Mean Substance Use across Sociodemographic Factors** | | | |
| --- | --- | --- | --- |
| **Substance** | **Population** | ***F* statistic Mean** | **P value Mean** |
| All | Sex | 7.23 | 0.0072 |
|  | Income | 0.88 | 0.49 |
|  | Race/ethnicity | 1.06 | 0.37 |
|  | Study site | 1.56 | 0.18 |
| Alcohol | Sex | 3.41 | 0.065 |
|  | Income | 2.64 | 0.022 |
|  | Race/ethnicity | 1.68 | 0.15 |
|  | Study site | 2.16 | 0.071 |
| Cannabis | Sex | 4.55 | 0.033 |
|  | Income | 2.17 | 0.055 |
|  | Race/ethnicity | 1.06 | 0.37 |
|  | Study site | 1.44 | 0.22 |
| Nicotine | Sex | 6.03 | 0.014 |
|  | Income | 0.7 | 0.62 |
|  | Race/ethnicity | 0.99 | 0.41 |
|  | Study site | 1.5 | 0.2 |
| Binge | Sex | 3.79 | 0.052 |
|  | Income | 0.66 | 0.65 |
|  | Race/ethnicity | 0.44 | 0.78 |
|  | Study site | 0.28 | 0.89 |
| **Mean Impulsivity (UPPS) across Sociodemographic Factors** | | | |
| **Subscale** | **Population** | ***F* statistic Mean** | **P value Mean** |
| Mean | Sex | 23.63 | <.001 |
| Mean | Income | 0.25 | 0.94 |
| Mean | Race/ethnicity | 4.07 | <.001 |
| Mean | Site | 4.87 | <.001 |
| Negative Urgency | Sex | 0.25 | 0.62 |
| Negative Urgency | Income | 1.01 | 0.41 |
| Negative Urgency | Race/ethnicity | 2.8 | 0.02 |
| Negative Urgency | Site | 3.02 | 0.02 |
| Perseverance | Sex | 0.49 | 0.48 |
| Perseverance | Income | 0.72 | 0.61 |
| Perseverance | Race/ethnicity | 1.98 | 0.09 |
| Perseverance | Site | 1.88 | 0.11 |
| Positive Urgency | Sex | 35.52 | <.001 |
| Positive Urgency | Income | 2.68 | 0.02 |
| Positive Urgency | Race/ethnicity | 4.18 | <.001 |
| Positive Urgency | Site | 2.27 | 0.06 |
| Premeditation | Sex | 0.02 | 0.89 |
| Premeditation | Income | 0.67 | 0.64 |
| Premeditation | Race/ethnicity | 1.87 | 0.11 |
| Premeditation | Site | 3.83 | <.001 |
| Sensation Seeking | Sex | 65.94 | <.001 |
| Sensation Seeking | Income | 2.39 | 0.04 |
| Sensation Seeking | Race/ethnicity | 5.87 | <.001 |
| Sensation Seeking | Site | 3.01 | 0.02 |
| **Mean Inhibitory Control (Anti-Saccade) across Sociodemographic Factors** | | | |
| **Trial Type** | **Population** | ***F* statistic Mean** | **P value Mean** |
| All | Sex | .04 | .84 |
|  | Income | 1.59 | .16 |
|  | Race/ethnicity | 2.57 | .037 |
|  | Study site | 32.01 | <.001 |
| Reward | Sex | .02 | .89 |
|  | Income | 1.84 | .10 |
|  | Race/ethnicity | 2.45 | .045 |
|  | Study site | 27.57 | <.001 |
| Neutral | Sex | .36 | .55 |
|  | Income | 1.65 | .14 |
|  | Race/ethnicity | 2.40 | .049 |
|  | Study site | 31.99 | <.001 |
| **Mean Tissue Iron (nT2*w) across Sociodemographic Factors** | | | |
| **Region of Interest** | **Population** | ***F* statistic** | **P value** |
| Basal Ganglia | Sex | 51.94 | <.001 |
| Basal Ganglia | Income | 2.07 | .07 |
| Basal Ganglia | Race/ethnicity | 4.76 | <.001 |
| Basal Ganglia | Site | 0.34 | .85 |
| Caudate | Sex | 99.16 | <.001 |
| Caudate | Income | 2.00 | .08 |
| Caudate | Race/ethnicity | 1.83 | .12 |
| Caudate | Site | 0.44 | .78 |
| NAcc | Sex | 0.76 | .38 |
| NAcc | Income | 2.17 | .06 |
| NAcc | Race/ethnicity | 2.07 | .08 |
| NAcc | Site | 0.72 | .58 |
| Pallidum | Sex | 13.74 | <.001 |
| Pallidum | Income | 0.53 | .75 |
| Pallidum | Race/ethnicity | 4.89 | <.001 |
| Pallidum | Site | 0.72 | .58 |
| Putamen | Sex | 50.85 | <.001 |
| Putamen | Income | 1.54 | .17 |
| Putamen | Race/ethnicity | 7.47 | <.001 |
| Putamen | Site | 0.47 | .76 |

*Note. All metrics result from independent generalized additive mixed models (GAMMs) described in the main text. F and p values describe the significance an ANOVA test across sociodemographic factors, controlling for a smooth term for age.*

**Supplementary Table S6. Sociodemographic Subgroups: Differences in the Magnitude of Peak Substance use, Impulsivity, Inhibitory Control, and Tissue Iron across Individual Substances, Subscales, Trial Types, and Regions of Interest**

| **Peak Substance Use across Sociodemographic Factors** | | | |
| --- | --- | --- | --- |
| **Substance** | **Population** | ***F* statistic Peak** | **P value Peak** |
| All | Sex | 21.83 | <.001 |
|  | Income | 1.81 | 0.11 |
|  | Race/ethnicity | 4.95 | <.001 |
|  | Site | 0.23 | 0.92 |
| Alcohol | Sex | 15.3 | .001 |
|  | Income | 5.26 | <.001 |
|  | Race/ethnicity | 5.8 | <.001 |
|  | Site | 1.29 | 0.27 |
| Binge | Sex | 6.44 | 0.01 |
|  | Income | 0.64 | 0.67 |
|  | Race/ethnicity | 1.29 | 0.27 |
|  | Site | 0.81 | 0.52 |
| Cannabis | Sex | 9.82 | 0.002 |
|  | Income | 2.4 | 0.037 |
|  | Race/ethnicity | 3.16 | 0.014 |
|  | Site | 0.6 | 0.67 |
| Nicotine | Sex | 2.79 | 0.096 |
|  | Income | 1.29 | 0.27 |
|  | Race/ethnicity | 2.14 | 0.076 |
|  | Site | 1.14 | 0.34 |
| **Peak Impulsivity (UPPS) across Sociodemographic Factors** | | | |
| **Subscale** | **Population** | ***F* statistic Mean Peak** | **P value Mean Peak** |
| Mean | Sex | 13.57 | <.001 |
| Mean | Income | 0.73 | 0.6 |
| Mean | Race/ethnicity | 0.95 | 0.43 |
| Mean | Site | 2.83 | 0.02 |
| Negative Urgency | Sex | 0.16 | 0.69 |
| Negative Urgency | Income | 1.49 | 0.19 |
| Negative Urgency | Race/ethnicity | 1.77 | 0.13 |
| Negative Urgency | Site | 1.39 | 0.23 |
| Perseverance | Sex | 1.57 | 0.21 |
| Perseverance | Income | 0.97 | 0.44 |
| Perseverance | Race/ethnicity | 1.34 | 0.25 |
| Perseverance | Site | 1.95 | 0.1 |
| Positive Urgency | Sex | 31.08 | <.001 |
| Positive Urgency | Income | 3.33 | 0.01 |
| Positive Urgency | Race/ethnicity | 3.19 | 0.01 |
| Positive Urgency | Site | 0.67 | 0.61 |
| Premeditation | Sex | 1.75 | 0.19 |
| Premeditation | Income | 0.62 | 0.68 |
| Premeditation | Race/ethnicity | 0.82 | 0.51 |
| Premeditation | Site | 4.77 | <.001 |
| Sensation Seeking | Sex | 59.27 | <.001 |
| Sensation Seeking | Income | 0.49 | 0.78 |
| Sensation Seeking | Race/ethnicity | 2 | 0.09 |
| Sensation Seeking | Site | 2.95 | 0.02 |
| **Peak inhibitory Control (Anti-Saccade) Performance across Sociodemographic Factors** | | | |
| **Trial Type** | **Population** | ***F* statistic Mean Peak** | **P value Mean Peak** |
| Mean | Sex | 0.04 | .84 |
|  | Income | 1.42 | .22 |
|  | Race/ethnicity | 2.07 | .086 |
|  | Site | 12.46 | <.001 |
| Reward | Sex | .05 | .82 |
|  | Income | 1.33 | .25 |
|  | Race/ethnicity | 1.92 | .11 |
|  | Site | 9.42 | .002 |
| Neutral | Sex | 0 | .98 |
|  | Income | 1.53 | .18 |
|  | Race/ethnicity | 1.83 | .12 |
|  | Site | 11.49 | <.001 |
| **Peak Tissue Iron (nT2*w) across Sociodemographic Factors** | | | |
| **Region of Interest** | **Population** | ***F* statistic Mean Peak** | **P value Mean Peak** |
| Basal Ganglia | Sex | 27.77 | <.001 |
| Basal Ganglia | Income | 3.06 | .01 |
| Basal Ganglia | Race/ethnicity | 2.54 | .04 |
| Basal Ganglia | Site | 1.38 | .24 |
| Caudate | Sex | 50.39 | <.001 |
| Caudate | Income | 2.37 | .04 |
| Caudate | Race/ethnicity | 1.51 | .2 |
| Caudate | Site | 0.59 | .67 |
| NAcc | Sex | 0.2 | .66 |
| NAcc | Income | 2.41 | .04 |
| NAcc | Race/ethnicity | 3.03 | .02 |
| NAcc | Site | 0.95 | .43 |
| Pallidum | Sex | 4.8 | .03 |
| Pallidum | Income | 1.6 | .16 |
| Pallidum | Race/ethnicity | 1.43 | .22 |
| Pallidum | Site | 4.5 | <.001 |
| Putamen | Sex | 27.34 | <.001 |
| Putamen | Income | 2.67 | .02 |
| Putamen | Race/ethnicity | 4.53 | <.001 |
| Putamen | Site | 2.28 | .06 |

*Note. All metrics result from independent generalized additive mixed models (GAMMs) described in the main text. F and p values describe the significance an ANOVA test across sociodemographic factors, controlling for a smooth term for age.*

**Supplementary Table S7A. Characteristics Among Substance use Frequency & Use-Type Categories across Individual Substances**

| **Frequency Categories (never, ever, regular use)** | | | | | | |
| --- | --- | --- | --- | --- | --- | --- |
| Substance | Category | Mean use (days) | SE use (days) | Mean Age (years) | SE Age (years) | N visits |
| All | Never | 0.00 | 0.00 | 16.89 | 0.07 | 578 |
| All | Ever | 1.58 | 0.06 | 20.17 | 0.07 | 633 |
| All | Regular | 13.81 | 0.21 | 22.12 | 0.06 | 551 |
| Alcohol | Never | 0.00 | 0.00 | 17.00 | 0.06 | 607 |
| Alcohol | Ever | 1.30 | 0.04 | 20.37 | 0.06 | 650 |
| Alcohol | Regular | 6.88 | 0.14 | 22.57 | 0.06 | 489 |
| Binge | Never | 0.00 | 0.00 | 18.13 | 0.06 | 723 |
| Binge | Ever | 0.68 | 0.03 | 21.63 | 0.06 | 574 |
| Binge | Regular | 3.44 | 0.15 | 22.29 | 0.10 | 221 |
| Cannabis | Never | 0.00 | 0.00 | 18.60 | 0.07 | 688 |
| Cannabis | Ever | 0.72 | 0.04 | 20.87 | 0.08 | 495 |
| Cannabis | Regular | 12.33 | 0.32 | 21.78 | 0.07 | 324 |
| Nicotine | Never | 0.00 | 0.00 | 19.02 | 0.06 | 747 |
| Nicotine | Ever | 0.34 | 0.02 | 21.71 | 0.08 | 411 |
| Nicotine | Regular | 18.81 | 0.46 | 21.28 | 0.11 | 171 |
| **Use-Type Categories (no-use, single-use, co-use, poly-use)** | | | | | | |
| **Substance** | **Category** | **Mean use (days)** | **SE use (days)** | **Mean Age (years)** | **SE Age (years)** | **N visits** |
| All | no-use | 0.00 | 0.00 | 16.89 | 0.07 | 578 |
| All | single-use | 5.62 | 0.18 | 21.27 | 0.08 | 575 |
| All | co-use | 8.19 | 0.28 | 21.43 | 0.09 | 457 |
| All | poly-use | 11.60 | 0.31 | 21.04 | 0.08 | 445 |
| Alcohol | no-use | 0.00 | 0.00 | 17.00 | 0.06 | 607 |
| Alcohol | single-use | 2.40 | 0.09 | 21.28 | 0.08 | 497 |
| Alcohol | co-use | 4.61 | 0.12 | 21.59 | 0.07 | 526 |
| Alcohol | poly-use | 5.10 | 0.25 | 20.96 | 0.11 | 234 |
| Binge | no-use | 0.00 | 0.00 | 18.13 | 0.06 | 723 |
| Binge | single-use | 0.91 | 0.06 | 21.70 | 0.09 | 349 |
| Binge | co-use | 1.38 | 0.07 | 21.81 | 0.07 | 457 |
| Binge | poly-use | 1.74 | 0.13 | 21.81 | 0.12 | 237 |
| Cannabis | no-use | 0.00 | 0.00 | 18.60 | 0.07 | 688 |
| Cannabis | single-use | 2.82 | 0.26 | 20.97 | 0.13 | 277 |
| Cannabis | co-use | 6.01 | 0.23 | 21.46 | 0.07 | 502 |
| Cannabis | poly-use | 7.41 | 0.44 | 20.96 | 0.11 | 234 |
| Nicotine | no-use | 0.00 | 0.00 | 19.02 | 0.06 | 747 |
| Nicotine | single-use | 0.67 | 0.10 | 22.60 | 0.12 | 222 |
| Nicotine | co-use | 3.74 | 0.28 | 21.42 | 0.10 | 319 |
| Nicotine | poly-use | 9.78 | 0.49 | 20.96 | 0.11 | 234 |

**Supplementary Table S7B. Associations Between Past 30-day Substance use and Impulsivity, Inhibitory Control, and Tissue Iron across Subscales, Trial Types, and Regions of Interest**

| **Impulsivity (UPPS)** | | | | |
| --- | --- | --- | --- | --- |
| **Substance** | **Subscale** | **Population** | ***F* statistic** | ***p* statistic** |
| All | Mean | Full Sample | 33.54 | <.001 |
| All | Negative Urgency | Full Sample | 16.50 | <.001 |
| All | Perseverance | Full Sample | 7.71 | .007 |
| All | Positive Urgency | Full Sample | 13.23 | <.001 |
| All | Premeditation | Full Sample | 24.22 | <.001 |
| All | Sensation Seeking | Full Sample | 18.60 | <.001 |
| Alcohol | Mean | Full Sample | 26.83 | <.001 |
| Alcohol | Negative Urgency | Full Sample | 2.76 | .10 |
| Alcohol | Perseverance | Full Sample | 5.78 | .02 |
| Alcohol | Positive Urgency | Full Sample | 9.77 | <.001 |
| Alcohol | Premeditation | Full Sample | 14.92 | <.001 |
| Alcohol | Sensation Seeking | Full Sample | 18.39 | <.001 |
| Binge | Mean | Full Sample | 16.98 | <.001 |
| Binge | Negative Urgency | Full Sample | 3.89 | .05 |
| Binge | Perseverance | Full Sample | 1.51 | .22 |
| Binge | Positive Urgency | Full Sample | 14.09 | <.001 |
| Binge | Premeditation | Full Sample | 14.37 | <.001 |
| Binge | Sensation Seeking | Full Sample | 6.11 | .02 |
| Cannabis | Mean | Full Sample | 31.65 | <.001 |
| Cannabis | Negative Urgency | Full Sample | 26.91 | <.001 |
| Cannabis | Perseverance | Full Sample | 10.45 | .001 |
| Cannabis | Positive Urgency | Full Sample | 8.94 | .002 |
| Cannabis | Premeditation | Full Sample | 16.48 | <.001 |
| Cannabis | Sensation Seeking | Full Sample | 14.30 | <.001 |
| Nicotine | Mean | Full Sample | 33.05 | <.001 |
| Nicotine | Negative Urgency | Full Sample | 16.90 | <.001 |
| Nicotine | Perseverance | Full Sample | 2.41 | .18 |
| Nicotine | Positive Urgency | Full Sample | 13.27 | <.001 |
| Nicotine | Premeditation | Full Sample | 13.59 | <.001 |
| Nicotine | Sensation Seeking | Full Sample | 8.24 | .007 |
| **Inhibitory Control (Anti-Saccade)** | | | | |
| **Substance** | **Trial Type** | **Population** | ***F* statistic** | ***p* statistic** |
| All | Mean | Full Sample | 0.48 | 0.49 |
| All | Reward | Full Sample | 0.83 | 0.36 |
| All | Neutral | Full Sample | 0.01 | 0.90 |
| Alcohol | Mean | Full Sample | 0.07 | 0.79 |
| Alcohol | Reward | Full Sample | 0.79 | 0.38 |
| Alcohol | Neutral | Full Sample | 0.02 | 0.90 |
| Binge | Mean | Full Sample | 0.01 | 0.93 |
| Binge | Reward | Full Sample | 0.36 | 0.55 |
| Binge | Neutral | Full Sample | 0.05 | 0.82 |
| Cannabis | Mean | Full Sample | 1.48 | 0.22 |
| Cannabis | Reward | Full Sample | 1.01 | 0.32 |
| Cannabis | Neutral | Full Sample | 1.68 | 0.31 |
| Nicotine | Mean | Full Sample | 0.07 | 0.80 |
| Nicotine | Reward | Full Sample | 0.03 | 0.85 |
| Nicotine | Neutral | Full Sample | 0.01 | 0.94 |
| **Tissue Iron (nT2*w)** | | | | |
| **Substance** | **ROI** | **Population** | ***F* statistic** | ***p* statistic** |
| All | Basal Ganglia | Full Sample | 4.22 | 0.01 |
| All | Caudate | Full Sample | 1.62 | 0.2 |
| All | NAcc | Full Sample | 1.39 | 0.25 |
| All | Pallidum | Full Sample | 0.61 | 0.54 |
| All | Putamen | Full Sample | 4.03 | 0.018 |
| Alcohol | Basal Ganglia | Full Sample | 5.69 | 0.0034 |
| Alcohol | Caudate | Full Sample | 2.06 | 0.13 |
| Alcohol | NAcc | Full Sample | 1.61 | 0.2 |
| Alcohol | Pallidum | Full Sample | 1.52 | 0.22 |
| Alcohol | Putamen | Full Sample | 4.55 | 0.011 |
| Binge | Basal Ganglia | Full Sample | 3.51 | 0.03 |
| Binge | Caudate | Full Sample | 0.71 | 0.49 |
| Binge | NAcc | Full Sample | 1.51 | 0.22 |
| Binge | Pallidum | Full Sample | 1.17 | 0.31 |
| Binge | Putamen | Full Sample | 3.31 | 0.036 |
| Cannabis | Basal Ganglia | Full Sample | 4.66 | 0.0095 |
| Cannabis | Caudate | Full Sample | 3.96 | 0.019 |
| Cannabis | NAcc | Full Sample | 0.44 | 0.65 |
| Cannabis | Pallidum | Full Sample | 0.28 | 0.76 |
| Cannabis | Putamen | Full Sample | 5.76 | 0.0032 |
| Nicotine | Basal Ganglia | Full Sample | 1.62 | 0.2 |
| Nicotine | Caudate | Full Sample | 1.66 | 0.19 |
| Nicotine | NAcc | Full Sample | 0.67 | 0.51 |
| Nicotine | Pallidum | Full Sample | 0.17 | 0.84 |
| Nicotine | Putamen | Full Sample | 1.19 | 0.3 |

*Note. All metrics result from independent generalized additive mixed models (GAMMs) described in the main text. F and p values describe the significance an ANOVA test across sociodemographic factors, controlling for a smooth term for age and relevant sociodemographic variables.*

**Supplementary Table S7C. Differences in Impulsivity, Inhibitory Control, and Tissue Iron within Substance Use Category Groups across Subscales, Trial Types, and Regions of Interest**

| **Impulsivity (UPPS)** | | | | | |
| --- | --- | --- | --- | --- | --- |
| **Substance** | **Subscale** | **Population** | **Grouping** | ***F* statistic** | ***p* statistic** |
| All | Mean | Full Sample | Frequency Category | 24.63 | <.001 |
| All | Negative Urgency | Full Sample | Frequency Category | 12.46 | <.001 |
| All | Perseverance | Full Sample | Frequency Category | 3.01 | .0494 |
| All | Positive Urgency | Full Sample | Frequency Category | 8.13 | .00029 |
| All | Premeditation | Full Sample | Frequency Category | 11.30 | <.001 |
| All | Sensation Seeking | Full Sample | Frequency Category | 13.03 | <.001 |
| Alcohol | Mean | Full Sample | Frequency Category | 23.77 | <.001 |
| Alcohol | Negative Urgency | Full Sample | Frequency Category | 11.32 | <.001 |
| Alcohol | Perseverance | Full Sample | Frequency Category | 5.88 | .0028 |
| Alcohol | Positive Urgency | Full Sample | Frequency Category | 5.99 | .0025 |
| Alcohol | Premeditation | Full Sample | Frequency Category | 10.35 | <.001 |
| Alcohol | Sensation Seeking | Full Sample | Frequency Category | 11.70 | <.001 |
| Binge | Mean | Full Sample | Frequency Category | 10.45 | <.001 |
| Binge | Negative Urgency | Full Sample | Frequency Category | 3.61 | .02 |
| Binge | Perseverance | Full Sample | Frequency Category | 1.75 | .17 |
| Binge | Positive Urgency | Full Sample | Frequency Category | 4.27 | .0139 |
| Binge | Premeditation | Full Sample | Frequency Category | 7.56 | <.001 |
| Binge | Sensation Seeking | Full Sample | Frequency Category | 5.65 | .003 |
| Cannabis | Mean | Full Sample | Frequency Category | 28.97 | <.001 |
| Cannabis | Negative Urgency | Full Sample | Frequency Category | 19.55 | <.001 |
| Cannabis | Perseverance | Full Sample | Frequency Category | 5.54 | .0039 |
| Cannabis | Positive Urgency | Full Sample | Frequency Category | 13.94 | <.001 |
| Cannabis | Premeditation | Full Sample | Frequency Category | 13.58 | <.001 |
| Cannabis | Sensation Seeking | Full Sample | Frequency Category | 5.92 | .0027 |
| Nicotine | Mean | Full Sample | Frequency Category | 24.69 | <.001 |
| Nicotine | Negative Urgency | Full Sample | Frequency Category | 19.66 | <.001 |
| Nicotine | Perseverance | Full Sample | Frequency Category | 3.97 | .0189 |
| Nicotine | Positive Urgency | Full Sample | Frequency Category | 8.54 | <.001 |
| Nicotine | Premeditation | Full Sample | Frequency Category | 12.26 | <.001 |
| Nicotine | Sensation Seeking | Full Sample | Frequency Category | 6.67 | .001 |
| **Inhibitory Control (Anti-Saccade)** | | | | | |
| **Substance** | **Trial Type** | **Population** | **Grouping** | ***F* statistic** | ***p* statistic** |
| All | Mean | Full Sample | Frequency Category | 5.11 | 0.006 |
| All | Reward | Full Sample | Frequency Category | 2.63 | 0.073 |
| All | Neutral | Full Sample | Frequency Category | 1.37 | 0.26 |
| Alcohol | Mean | Full Sample | Frequency Category | 1.97 | 0.14 |
| Alcohol | Reward | Full Sample | Frequency Category | 0.96 | 0.38 |
| Alcohol | Neutral | Full Sample | Frequency Category | 0.3 | 0.74 |
| Binge | Mean | Full Sample | Frequency Category | 0.96 | 0.38 |
| Binge | Reward | Full Sample | Frequency Category | 0.15 | 0.86 |
| Binge | Neutral | Full Sample | Frequency Category | 0.33 | 0.72 |
| Cannabis | Mean | Full Sample | Frequency Category | 3.35 | 0.036 |
| Cannabis | Reward | Full Sample | Frequency Category | 0.72 | 0.49 |
| Cannabis | Neutral | Full Sample | Frequency Category | 1.24 | 0.29 |
| Nicotine | Mean | Full Sample | Frequency Category | 4.01 | 0.018 |
| Nicotine | Reward | Full Sample | Frequency Category | 2.2 | 0.11 |
| Nicotine | Neutral | Full Sample | Frequency Category | 0.94 | 0.39 |
| **Tissue Iron (nT2*w)** | | | | | |
| **Substance** | **ROI** | **Population** | **Grouping** | ***F* statistic** | ***p* statistic** |
| All | Basal Ganglia | Full Sample | Frequency Category | 3.37 | 0.02 |
| All | Caudate | Full Sample | Frequency Category | 2.27 | 0.13 |
| All | NAcc | Full Sample | Frequency Category | 1.63 | 0.20 |
| All | Pallidum | Full Sample | Frequency Category | 3.83 | 0.01 |
| All | Putamen | Full Sample | Frequency Category | 5.20 | 0.02 |
| Alcohol | Basal Ganglia | Full Sample | Frequency Category | 9.99 | <.001 |
| Alcohol | Caudate | Full Sample | Frequency Category | 6.30 | 0.01 |
| Alcohol | NAcc | Full Sample | Frequency Category | 1.22 | 0.27 |
| Alcohol | Pallidum | Full Sample | Frequency Category | 2.80 | 0.04 |
| Alcohol | Putamen | Full Sample | Frequency Category | 6.18 | 0.01 |
| Binge | Basal Ganglia | Full Sample | Frequency Category | 1.50 | 0.18 |
| Binge | Caudate | Full Sample | Frequency Category | 0.50 | 0.48 |
| Binge | NAcc | Full Sample | Frequency Category | 0.06 | 0.81 |
| Binge | Pallidum | Full Sample | Frequency Category | 0.74 | 0.31 |
| Binge | Putamen | Full Sample | Frequency Category | 2.19 | 0.14 |
| Cannabis | Basal Ganglia | Full Sample | Frequency Category | 1.08 | 0.24 |
| Cannabis | Caudate | Full Sample | Frequency Category | 0.17 | 0.68 |
| Cannabis | NAcc | Full Sample | Frequency Category | 0.09 | 0.91 |
| Cannabis | Pallidum | Full Sample | Frequency Category | 1.49 | 0.18 |
| Cannabis | Putamen | Full Sample | Frequency Category | 0.97 | 0.24 |
| Nicotine | Basal Ganglia | Full Sample | Frequency Category | 0.80 | 0.29 |
| Nicotine | Caudate | Full Sample | Frequency Category | 0.37 | 0.54 |
| Nicotine | NAcc | Full Sample | Frequency Category | 1.77 | 0.14 |
| Nicotine | Pallidum | Full Sample | Frequency Category | 0.00 | 0.94 |
| Nicotine | Putamen | Full Sample | Frequency Category | 0.60 | 0.44 |

*Note. All metrics result from independent generalized additive mixed models (GAMMs) described in the main text. F and p values describe the significance an ANOVA test across sociodemographic factors, controlling for a smooth term for age and relevant sociodemographic variables.*

**Table S7D. Differences in Impulsivity, Inhibitory Control, and Tissue Iron within Substance Use-Type Groups across Subscales, Trial Types, and Regions of Interest**

| **Impulsivity (UPPS)** | | | | | |
| --- | --- | --- | --- | --- | --- |
| **Substance** | **Subscale** | **Population** | **Grouping** | ***F* statistic** | ***p* statistic** |
| All | Mean | Full Sample | Usetype Category | 18.33 | <.001 |
| All | Negative Urgency | Full Sample | Usetype Category | 7.52 | <.001 |
| All | Perseverance | Full Sample | Usetype Category | 3.42 | .016 |
| All | Positive Urgency | Full Sample | Usetype Category | 8.29 | <.001 |
| All | Premeditation | Full Sample | Usetype Category | 11.81 | <.001 |
| All | Sensation Seeking | Full Sample | Usetype Category | 8.24 | <.001 |
| Alcohol | Mean | Full Sample | Usetype Category | 24.46 | <.001 |
| Alcohol | Negative Urgency | Full Sample | Usetype Category | 12.24 | <.001 |
| Alcohol | Perseverance | Full Sample | Usetype Category | 7.42 | <.001 |
| Alcohol | Positive Urgency | Full Sample | Usetype Category | 8.61 | <.001 |
| Alcohol | Premeditation | Full Sample | Usetype Category | 8.61 | <.001 |
| Alcohol | Sensation Seeking | Full Sample | Usetype Category | 9.91 | <.001 |
| Binge | Mean | Full Sample | Usetype Category | 7.92 | <.001 |
| Binge | Negative Urgency | Full Sample | Usetype Category | 2.58 | .05 |
| Binge | Perseverance | Full Sample | Usetype Category | 3.23 | .02 |
| Binge | Positive Urgency | Full Sample | Usetype Category | 2.92 | .03 |
| Binge | Premeditation | Full Sample | Usetype Category | 4.91 | .002 |
| Binge | Sensation Seeking | Full Sample | Usetype Category | 3.58 | .01 |
| Cannabis | Mean | Full Sample | Usetype Category | 18.35 | <.001 |
| Cannabis | Negative Urgency | Full Sample | Usetype Category | 11.26 | <.001 |
| Cannabis | Perseverance | Full Sample | Usetype Category | 3.76 | .01 |
| Cannabis | Positive Urgency | Full Sample | Usetype Category | 9.36 | <.001 |
| Cannabis | Premeditation | Full Sample | Usetype Category | 8.59 | <.001 |
| Cannabis | Sensation Seeking | Full Sample | Usetype Category | 4.74 | .0026 |
| Nicotine | Mean | Full Sample | Usetype Category | 14.22 | <.001 |
| Nicotine | Negative Urgency | Full Sample | Usetype Category | 12.35 | <.001 |
| Nicotine | Perseverance | Full Sample | Usetype Category | 2.68 | .04 |
| Nicotine | Positive Urgency | Full Sample | Usetype Category | 3.72 | .01 |
| Nicotine | Premeditation | Full Sample | Usetype Category | 5.71 | <.001 |
| Nicotine | Sensation Seeking | Full Sample | Usetype Category | 5.31 | .001 |
| **Inhibitory Control (Anti-Saccade)** | | | | | |
| **Substance** | **Trial Type** | **Population** | **Grouping** | ***F* statistic** | ***p* statistic** |
| All | Mean | Full Sample | Usetype Category | 5.22 | 0.0014 |
| All | Reward | Full Sample | Usetype Category | 1.79 | 0.15 |
| All | Neutral | Full Sample | Usetype Category | 1.83 | 0.14 |
| Alcohol | Mean | Full Sample | Usetype Category | 1.35 | 0.26 |
| Alcohol | Reward | Full Sample | Usetype Category | 1.06 | 0.36 |
| Alcohol | Neutral | Full Sample | Usetype Category | 0.13 | 0.94 |
| Binge | Mean | Full Sample | Usetype Category | 0.59 | 0.62 |
| Binge | Reward | Full Sample | Usetype Category | 0.05 | 0.99 |
| Binge | Neutral | Full Sample | Usetype Category | 0.36 | 0.78 |
| Cannabis | Mean | Full Sample | Usetype Category | 1.69 | 0.17 |
| Cannabis | Reward | Full Sample | Usetype Category | 0.61 | 0.61 |
| Cannabis | Neutral | Full Sample | Usetype Category | 0.49 | 0.69 |
| Nicotine | Mean | Full Sample | Usetype Category | 3.37 | 0.018 |
| Nicotine | Reward | Full Sample | Usetype Category | 2.35 | 0.072 |
| Nicotine | Neutral | Full Sample | Usetype Category | 1.33 | 0.26 |
| **Tissue Iron (nT2*w)** | | | | | |
| **Substance** | **ROI** | **Population** | **Grouping** | ***F* statistic** | ***p* statistic** |
| All | Basal Ganglia | Full Sample | Usetype Category | 1.14 | 0.33 |
| All | Caudate | Full Sample | Usetype Category | 1.51 | 0.21 |
| All | NAcc | Full Sample | Usetype Category | 0.97 | 0.41 |
| All | Pallidum | Full Sample | Usetype Category | 0.91 | 0.43 |
| All | Putamen | Full Sample | Usetype Category | 0.38 | 0.77 |
| Alcohol | Basal Ganglia | Full Sample | Usetype Category | 0.88 | 0.45 |
| Alcohol | Caudate | Full Sample | Usetype Category | 0.83 | 0.48 |
| Alcohol | NAcc | Full Sample | Usetype Category | 0.04 | 0.99 |
| Alcohol | Pallidum | Full Sample | Usetype Category | 0.84 | 0.47 |
| Alcohol | Putamen | Full Sample | Usetype Category | 1.2 | 0.31 |
| Binge | Basal Ganglia | Full Sample | Usetype Category | 3.63 | 0.01 |
| Binge | Caudate | Full Sample | Usetype Category | 2.04 | 0.11 |
| Binge | NAcc | Full Sample | Usetype Category | 0.74 | 0.53 |
| Binge | Pallidum | Full Sample | Usetype Category | 1.06 | 0.36 |
| Binge | Putamen | Full Sample | Usetype Category | 4.42 | 0.004 |
| Cannabis | Basal Ganglia | Full Sample | Usetype Category | 3.62 | 0.013 |
| Cannabis | Caudate | Full Sample | Usetype Category | 2.15 | 0.091 |
| Cannabis | NAcc | Full Sample | Usetype Category | 0.94 | 0.42 |
| Cannabis | Pallidum | Full Sample | Usetype Category | 0.31 | 0.82 |
| Cannabis | Putamen | Full Sample | Usetype Category | 4.58 | 0.003 |
| Nicotine | Basal Ganglia | Full Sample | Usetype Category | 1.65 | 0.18 |
| Nicotine | Caudate | Full Sample | Usetype Category | 1.43 | 0.23 |
| Nicotine | NAcc | Full Sample | Usetype Category | 0.52 | 0.67 |
| Nicotine | Pallidum | Full Sample | Usetype Category | 0.31 | 0.82 |
| Nicotine | Putamen | Full Sample | Usetype Category | 1.48 | 0.22 |

*Note. All metrics result from independent generalized additive mixed models (GAMMs) described in the main text. F and p values describe the significance an ANOVA test across sociodemographic factors, controlling for a smooth term for age and relevant sociodemographic variables.*

**Supplementary Table S8. Trajectory Groups: Statistics Across Individual Substances, Subscales, Trial Types, and Regions of Interest**

| **Substance Use: Statistics Across Individual Substances and Trajectory Groups** | | | | | | | | |
| --- | --- | --- | --- | --- | --- | --- | --- | --- |
| Substance | Population | | Mean n days | SE n days | Mean Peak Use (days) | SE Peak Use (days) | Mean Age of initiation (years) | Mean Age of peak use (years) |
| All | Low | | 0.35 | 0.04 | 3.25 | 0.12 | 17.77 | 21.59 |
| All | Youth Peak | | 11.9 | 0.27 | 22.63 | 0.2 | 15.65 | 21.89 |
| All | Adolescent Increasing | | 8.17 | 0.32 | 23.28 | 0.21 | 16.39 | 21.78 |
| All | Adult Increasing | | 3.26 | 0.13 | 10.76 | 0.18 | 17.38 | 23.01 |
| Alcohol | Low | | 0.23 | 0.02 | 2.3 | 0.05 | 18.19 | 21.85 |
| Alcohol | Youth Peak | | 4.82 | 0.14 | 12.27 | 0.17 | 16.08 | 22.09 |
| Alcohol | Adolescent Increasing | | 3.31 | 0.17 | 11.1 | 0.22 | 16.72 | 21.75 |
| Alcohol | Adult Increasing | | 2.1 | 0.08 | 6.82 | 0.12 | 17.68 | 22.97 |
| Binge | Low | | 0.02 | 0 | 1.32 | 0.06 | 19.38 | 20.81 |
| Binge | Youth Peak | | 1.36 | 0.06 | 4.88 | 0.11 | 17.51 | 21.1 |
| Binge | Adolescent Increasing | | 0.93 | 0.07 | 4.63 | 0.16 | 17.98 | 21.33 |
| Binge | Adult Increasing | | 0.31 | 0.02 | 2.24 | 0.07 | 19.1 | 22.08 |
| Cannabis | Low | | 0.08 | 0.03 | 3.32 | 0.41 | 16.97 | 20.54 |
| Cannabis | Youth Peak | | 6.28 | 0.25 | 16.86 | 0.31 | 16.51 | 21.47 |
| Cannabis | Adolescent Increasing | | 4.33 | 0.26 | 16.23 | 0.36 | 17.21 | 21.48 |
| Cannabis | Adult Increasing | | 0.8 | 0.08 | 7.15 | 0.29 | 18.33 | 21.99 |
| Nicotine | Low | | 0.03 | 0.01 | 2.69 | 0.27 | 17.42 | 20.28 |
| Nicotine | Youth Peak | | 4.03 | 0.22 | 15.65 | 0.36 | 17.4 | 20.97 |
| Nicotine | Adolescent Increasing | | 2.24 | 0.21 | 14.9 | 0.46 | 18.24 | 21.61 |
| Nicotine | Adult Increasing | | 0.22 | 0.05 | 6.63 | 0.57 | 18.35 | 21.01 |
| **Impulsivity (UPPS): Statistics Across Individual Subscales and Trajectory Groups** | | | | | | | | |
| **Subscale** | | **Population** | | **Mean Score** | **SE Score** | **Mean Peak Score** | **SE Peak Score** | **Mean Age of peak Score** |
| Mean across all subscales | | Low | | 1.87 | 0.01 | 2.18 | 0.01 | 17.97 |
| Mean across all subscales | | Youth Peak | | 2.03 | 0.01 | 2.32 | 0.01 | 18.58 |
| Mean across all subscales | | Adolescent Increasing | | 1.99 | 0.01 | 2.31 | 0.01 | 17.48 |
| Mean across all subscales | | Adult Increasing | | 1.83 | 0.01 | 2.1 | 0.01 | 18.57 |
| Negative Urgency | | Low | | 1.83 | 0.02 | 2.48 | 0.02 | 17.84 |
| Negative Urgency | | Youth Peak | | 1.95 | 0.02 | 2.57 | 0.02 | 18.82 |
| Negative Urgency | | Adolescent Increasing | | 1.95 | 0.02 | 2.65 | 0.02 | 17.79 |
| Negative Urgency | | Adult Increasing | | 1.73 | 0.02 | 2.29 | 0.02 | 18.41 |
| Perseverance | | Low | | 1.74 | 0.01 | 2.19 | 0.01 | 17.82 |
| Perseverance | | Youth Peak | | 1.79 | 0.01 | 2.19 | 0.01 | 18.69 |
| Perseverance | | Adolescent Increasing | | 1.77 | 0.02 | 2.19 | 0.01 | 17.35 |
| Perseverance | | Adult Increasing | | 1.75 | 0.01 | 2.16 | 0.01 | 18.7 |
| Positive Urgency | | Low | | 1.65 | 0.02 | 2.17 | 0.02 | 17.31 |
| Positive Urgency | | Youth Peak | | 1.75 | 0.02 | 2.33 | 0.02 | 18.41 |
| Positive Urgency | | Adolescent Increasing | | 1.72 | 0.02 | 2.32 | 0.02 | 17.12 |
| Positive Urgency | | Adult Increasing | | 1.47 | 0.02 | 1.94 | 0.02 | 18.28 |
| Premeditation | | Low | | 1.6 | 0.01 | 2.06 | 0.01 | 17.92 |
| Premeditation | | Youth Peak | | 1.72 | 0.02 | 2.17 | 0.01 | 18.6 |
| Premeditation | | Adolescent Increasing | | 1.69 | 0.02 | 2.14 | 0.01 | 17.35 |
| Premeditation | | Adult Increasing | | 1.55 | 0.01 | 1.96 | 0.01 | 18.53 |
| Sensation Seeking | | Low | | 2.52 | 0.02 | 3.03 | 0.02 | 17.78 |
| Sensation Seeking | | Youth Peak | | 2.93 | 0.02 | 3.39 | 0.01 | 18.63 |
| Sensation Seeking | | Adolescent Increasing | | 2.83 | 0.02 | 3.33 | 0.02 | 17.07 |
| Sensation Seeking | | Adult Increasing | | 2.63 | 0.02 | 3.09 | 0.02 | 18.46 |
| **Inhibitory Control (Anti-Saccade): Statistics Across Individual Subscales and Trajectory Groups** | | | | | | | | |
| **Subscale** | | **Population** | | **Mean performance (%)** | **SE (%)** | **Mean Peak performance (%)** | **SE Peak performance (%)** | **Mean Age of peak performance (years)** |
| Mean | | Low | | 82.03 | 0.88 | 88.11 | 0.72 | 17.76 |
| Mean | | Youth Peak | | 81.32 | 1.24 | 86.14 | 1.07 | 18.47 |
| Mean | | Adolescent Increasing | | 83.33 | 1.57 | 89.56 | 1.17 | 16.93 |
| Mean | | Adult Increasing | | 84.02 | 0.96 | 89.45 | 0.74 | 18.61 |
| Reward | | Low | | 83.95 | 0.86 | 89.95 | 0.66 | 17.69 |
| Reward | | Youth Peak | | 83.06 | 1.26 | 88.08 | 1.02 | 18.25 |
| Reward | | Adolescent Increasing | | 85.31 | 1.49 | 91.92 | 1.11 | 16.76 |
| Reward | | Adult Increasing | | 86.16 | 0.98 | 91.88 | 0.69 | 18.58 |
| Neutral | | Low | | 80.58 | 0.94 | 87.52 | 0.77 | 17.71 |
| Neutral | | Youth Peak | | 79.81 | 1.36 | 85.8 | 1.16 | 18.6 |
| Neutral | | Adolescent Increasing | | 82.58 | 1.64 | 88.93 | 1.24 | 17.14 |
| Neutral | | Adult Increasing | | 81.99 | 1.06 | 88.72 | 0.78 | 18.6 |
| **Tissue Iron (nT2*w): Statistics Across Individual Subscales and Trajectory Groups** | | | | | | | | |
| **Region** | | **Population** | | **Mean nT2*w** | **SE nT2*w** | **Mean Peak nT2*w** | **SE Peak nT2*2** | **Mean Age of peak nT2*w** |
| Basal Ganglia | | Low | | 0.01356 | .00002 | 0.01303 | .00002 | 19.88 |
| Basal Ganglia | | Youth Peak | | 0.01352 | .00002 | 0.01301 | .00002 | 20.93 |
| Basal Ganglia | | Adolescent Increasing | | 0.01355 | .00002 | 0.01301 | .00002 | 19.50 |
| Basal Ganglia | | Adult Increasing | | 0.01349 | .00002 | 0.01297 | .00002 | 21.27 |
| Caudate | | Low | | 0.014 | .00002 | 0.01338 | .00002 | 19.31 |
| Caudate | | Youth Peak | | 0.01401 | .00002 | 0.01344 | .00002 | 20.54 |
| Caudate | | Adolescent Increasing | | 0.01398 | .00002 | 0.01335 | .00002 | 19.02 |
| Caudate | | Adult Increasing | | 0.01399 | .00002 | 0.01341 | .00002 | 20.55 |
| NAcc | | Low | | 0.01109 | .00007 | 0.00964 | .00007 | 19.30 |
| NAcc | | Youth Peak | | 0.01116 | .00007 | 0.00961 | .00008 | 20.37 |
| NAcc | | Adolescent Increasing | | 0.01088 | .00008 | 0.00923 | .00008 | 18.81 |
| NAcc | | Adult Increasing | | 0.01065 | .00007 | 0.00919 | .00007 | 19.92 |
| Pallidum | | Low | | 0.01113 | .00002 | 0.01064 | .00002 | 20.47 |
| Pallidum | | Youth Peak | | 0.01106 | .00002 | 0.01059 | .00002 | 21.25 |
| Pallidum | | Adolescent Increasing | | 0.01118 | .00003 | 0.01068 | .00002 | 20.23 |
| Pallidum | | Adult Increasing | | 0.01106 | .00002 | 0.01055 | .00002 | 21.29 |
| Putamen | | Low | | 0.01439 | .00002 | 0.01385 | .00002 | 20.10 |
| Putamen | | Youth Peak | | 0.01432 | .00002 | 0.0138 | .00002 | 21.12 |
| Putamen | | Adolescent Increasing | | 0.01438 | .00002 | 0.01383 | .00002 | 19.62 |
| Putamen | | Adult Increasing | | 0.01432 | .00002 | 0.01378 | .00002 | 21.27 |

**Supplementary Table S9. Trajectory Groups: Developmental GAMM Model Statistics Across Individual Substances, Subscales, Trial Types, and Regions of Interest**

| **Substance Use: Statistics Across Individual Substances and Trajectory Groups** | | | | | | |
| --- | --- | --- | --- | --- | --- | --- |
| **Substance** | **Population** | **s(age) F** | **s(age) p** | **Period of increase** | **Period of decrease** | **Rate of Change** |
| All | Low | 11.70 | <.001 | 14.97 – 22.53 |  | 0.041 |
| All | Youth Peak | 407.98 | <.001 | 12.10 – 23.73 | 25.43 – 29.90 | 1.507 |
| All | Adolescent Increasing | 841.07 | <.001 | 15.86 – 27.54 |  | 2.096 |
| All | Adult Increasing | 248.17 | <.001 | 15.20 – 29.84 |  | 0.879 |
| Alcohol | Low | 38.95 | <.001 | 16.32 – 23.61 |  | 0.023 |
| Alcohol | Youth Peak | 242.88 | <.001 | 14.34 – 24.09 | 25.43 – 29.90 | 0.358 |
| Alcohol | Adolescent Increasing | 327.74 | <.001 | 15.94 – 27.54 |  | 1.070 |
| Alcohol | Adult Increasing | 268.01 | <.001 | 16.19 – 29.84 |  | 0.456 |
| Binge | Low | 0.0001 | .30 |  |  | 0.0001 |
| Binge | Youth Peak | 62.23 | <.001 | 12.10 – 22.21 | 23.46 – 29.90 | 0.068 |
| Binge | Adolescent Increasing | 86.72 | <.001 | 16.18 – 22.87 |  | 0.143 |
| Binge | Adult Increasing | 26.30 | <.001 | 12.08 – 24.22 |  | 0.051 |
| Cannabis | Low | 0.0001 | .72 |  |  | 0.0001 |
| Cannabis | Youth Peak | 149.67 | <.001 | 12.10 – 23.55 |  | 1.008 |
| Cannabis | Adolescent Increasing | 212.93 | <.001 | 16.18 – 27.54 |  | 1.230 |
| Cannabis | Adult Increasing | 38.69 | <.001 | 20.92 – 29.84 |  | 0.353 |
| Nicotine | Low | 0.23 | .20 |  |  | 0.001 |
| Nicotine | Youth Peak | 88.18 | <.001 | 12.10 – 22.92 |  | 0.749 |
| Nicotine | Adolescent Increasing | 86.43 | <.001 | 16.41 – 27.54 |  | 0.913 |
| Nicotine | Adult Increasing | 5.05 | <.001 | 21.54 – 29.84 |  | 0.061 |
| **Impulsivity (UPPS): Statistics Across Individual Substances and Trajectory Groups** | | | | | | |
| **Subscale** | **Population** | **s(age) F** | **s(age) p** | **Period of increase** | **Period of decrease** | **Rate of Change** |
| Mean | Low | -30.11 | <.001 | 12.00 | 12.00 - 29.30 | -0.025 |
| Mean | Youth Peak | -26.78 | <.001 | 13.43 | 13.43 - 26.23 | -0.022 |
| Mean | Adolescent Increasing | -3.18 | <.001 | 15.59 | 15.59 - 19.75 | -0.008 |
| Mean | Adult Increasing | -22.12 | <.001 | 12.42 | 12.42 - 24.87 | -0.017 |
| Negative Urgency | Low | -11.93 | <.001 | 12.00 | 12.00 - 24.43 | -0.030 |
| Negative Urgency | Youth Peak | -7.25 | <.001 | 16.67 | 16.67 - 24.32 | -0.023 |
| Negative Urgency | Adolescent Increasing | 0.00 | .80 |  |  | 0.000 |
| Negative Urgency | Adult Increasing | -11.55 | <.001 | 12.08 | 12.08 - 24.02 | -0.026 |
| Perseverance | Low | -5.22 | <.001 | 15.13 | 15.13 - 21.56 | -0.012 |
| Perseverance | Youth Peak | -2.29 | .01 | 17.50 | 17.50 - 19.41 | -0.009 |
| Perseverance | Adolescent Increasing | 0.00 | .32 |  |  | 0.000 |
| Perseverance | Adult Increasing | 0.00 | .92 |  |  | 0.000 |
| Positive Urgency | Low | -23.19 | <.001 | 12.00 | 12.00 - 29.30 | -0.039 |
| Positive Urgency | Youth Peak | -24.09 | <.001 | 12.10 | 12.10 - 24.73 | -0.038 |
| Positive Urgency | Adolescent Increasing | -8.47 | <.001 | 14.34 | 14.34 - 21.42 | -0.026 |
| Positive Urgency | Adult Increasing | -26.73 | 0.00 | 12.08 | 12.08 - 24.36 | -0.032 |
| Premeditation | Low | -9.45 | <.001 | 12.00 | 12.00 - 22.87 | -0.019 |
| Premeditation | Youth Peak | -11.57 | <.001 | 12.10 | 12.10 - 22.16 | -0.020 |
| Premeditation | Adolescent Increasing | -3.45 | <.001 | 14.96 | 14.96 - 17.46 | -0.008 |
| Premeditation | Adult Increasing | -8.42 | <.001 | 14.47 | 14.47 - 22.06 | -0.014 |
| Sensation Seeking | Low | -7.07 | <.001 | 19.56 | 19.56 - 29.30 | -0.026 |
| Sensation Seeking | Youth Peak | -8.30 | <.001 | 19.50 | 19.50 - 28.64 | -0.024 |
| Sensation Seeking | Adolescent Increasing | -1.44 | .07 |  |  | -0.010 |
| Sensation Seeking | Adult Increasing | -3.47 | <.001 | 19.58 | 19.58 - 23.08 | -0.011 |
| **Inhibitory Control (Anti-Saccade): Statistics Across Individual Substances and Trajectory Groups** | | | | | | |
| **Trial Type** | **Population** | **s(age) F** | **s(age) p** | **Period of increase** | **Period of decrease** | **Rate of Change** |
| Mean | Low | 5.32 | <.001 | 14.15 – 20.10 |  | 1.494 |
| Mean | Youth Peak | 2.18 | .01 |  |  | 1.300 |
| Mean | Adolescent Increasing | 2.56 | .01 | 15.04 – 16.11 |  | 1.717 |
| Mean | Adult Increasing | 0.43 | .13 |  |  | 0.361 |
| Reward | Low | 2.86 | <.001 | 15.39 – 18.68 |  | 1.025 |
| Reward | Youth Peak | 2.29 | .01 | 16.63 – 17.84 |  | 1.396 |
| Reward | Adolescent Increasing | 1.33 | .03 |  |  | 1.243 |
| Reward | Adult Increasing | 0.05 | .29 |  |  | 0.056 |
| Neutral | Low | 5.49 | <.001 | 13.74 – 20.21 |  | 1.683 |
| Neutral | Youth Peak | 2.06 | .01 |  |  | 1.368 |
| Neutral | Adolescent Increasing | 2.29 | .01 | 15.36 – 16.11 |  | 1.697 |
| Neutral | Adult Increasing | 0.21 | .20 |  |  | 0.225 |
| **Tissue Iron (nT2*w): Statistics Across Individual Substances and Trajectory Groups** | | | | | | |
| **Region** | **Population** | **s(age) F** | **s(age) p** | **Period of increase** | **Period of decrease** | **Rate of Change** |
| Basal Ganglia | Low | -68.91 | <.001 |  | 12.99 - 29.30 | -0.00008 |
| Basal Ganglia | Youth Peak | -83.37 | <.001 |  | 12.10 - 28.64 | -0.00009 |
| Basal Ganglia | Adolescent Increasing | -33.81 | <.001 |  | 14.64 - 26.76 | -0.00007 |
| Basal Ganglia | Adult Increasing | -75.49 | <.001 |  | 12.13 - 29.05 | -0.00007 |
| Caudate | Low | -18.25 | <.001 |  | 16.29 - 29.30 | -0.00005 |
| Caudate | Youth Peak | -30.07 | <.001 |  | 12.10 - 28.64 | -0.00006 |
| Caudate | Adolescent Increasing | -9.17 | <.001 |  | 18.78 - 26.76 | -0.00004 |
| Caudate | Adult Increasing | -24.67 | <.001 |  | 13.92 - 29.05 | -0.00005 |
| NAcc | Low | -7.76 | <.001 |  | 12.04 - 23.23 | -0.00007 |
| NAcc | Youth Peak | -20.54 | <.001 |  | 12.10 - 26.65 | -0.00014 |
| NAcc | Adolescent Increasing | -7.19 | <.001 |  | 12.05 - 16.93 | -0.00012 |
| NAcc | Adult Increasing | -10.85 | <.001 |  | 12.13 - 24.54 | -0.00009 |
| Pallidum | Low | -142.21 | <.001 |  | 12.04 - 29.30 | -0.00009 |
| Pallidum | Youth Peak | -126.92 | <.001 |  | 12.10 - 28.64 | -0.00009 |
| Pallidum | Adolescent Increasing | -126.77 | <.001 |  | 12.05 - 26.76 | -0.00010 |
| Pallidum | Adult Increasing | -132.35 | <.001 |  | 12.13 - 29.05 | -0.00009 |
| Putamen | Low | -93.80 | <.001 |  | 12.04 - 29.30 | -0.00009 |
| Putamen | Youth Peak | -97.87 | <.001 |  | 12.10 - 28.64 | -0.00009 |
| Putamen | Adolescent Increasing | -52.03 | <.001 |  | 14.79 - 26.76 | -0.00008 |
| Putamen | Adult Increasing | -92.95 | <.001 |  | 12.13 - 29.05 | -0.00007 |

*Note. All metrics result from independent generalized additive mixed models (GAMMs) described in the main text. F and p values describe the significance of the smooth term for age.*

**Supplementary Table S10. Trajectory Groups: Differences in the Magnitude of Substance use, Impulsivity, Inhibitory Control, and Tissue Iron across Individual Substances, Subscales, Trial Types, and Regions of Interest**

| **Mean Substance Use across Trajectory Groups** | | | |
| --- | --- | --- | --- |
| **Substance** | **Population** | ***F* statistic: Mean** | **P value** |
| All | Trajectory Group | 113.88 | <.001 |
| Alcohol | Trajectory Group | 66.05 | <.001 |
| Binge | Trajectory Group | 44.34 | <.001 |
| Cannabis | Trajectory Group | 25.79 | <.001 |
| Nicotine | Trajectory Group | 17.91 | <.001 |
| **Mean Impulsivity (UPPS) across Trajectory Groups** | | | |
| **Subscale** | **Population** | ***F* statistic: Mean** | **P value** |
| Mean | Trajectory Group | 14.13 | <.001 |
| Negative Urgency | Trajectory Group | 6.07 | <.001 |
| Perseverance | Trajectory Group | .58 | .60 |
| Positive Urgency | Trajectory Group | 7.69 | <.001 |
| Premeditation | Trajectory Group | 7.03 | <.001 |
| Sensation Seeking | Trajectory Group | 12.61 | <.001 |
| **Mean Inhibitory Control (Anti-Saccade) Performance across Trajectory Groups** | | | |
| **Trial Type** | **Population** | ***F* statistic: Mean** | **P value** |
| Mean | Trajectory Group | 1.52 | .21 |
| Reward | Trajectory Group | 1.29 | .28 |
| Neutral | Trajectory Group | 1.48 | .22 |
| **Mean Tissue Iron (nT2*w) across Trajectory Groups** | | | |
| **Region** | **Population** | ***F* statistic: Mean** | **P value** |
| Basal Ganglia | Trajectory Group | 1.80 | .150 |
| Caudate | Trajectory Group | 2.53 | .056 |
| NAcc | Trajectory Group | 3.07 | .027 |
| Pallidum | Trajectory Group | .32 | .810 |
| Putamen | Trajectory Group | .81 | .490 |

*Note. All metrics result from independent generalized additive mixed models (GAMMs) described in the main text. F and p values describe the significance an ANOVA test across trajectory groups, controlling for a smooth term for age and relevant sociodemographic variables.*

**Supplementary Table S11. Trajectory Groups: Differences in the Magnitude of Peak Substance use, Impulsivity, Inhibitory Control, and Tissue Iron across Individual Substances, Subscales, Trial Types, and Regions of Interest**

| **Peak Substance Use across Trajectory Groups** | | | |
| --- | --- | --- | --- |
| **Substance** | **Population** | ***F* statistic: Mean Peak** | **P value** |
| All | Trajectory Group | 252.42 | <.001 |
| Alcohol | Trajectory Group | 69.86 | <.001 |
| Binge | Trajectory Group | 15.76 | <.001 |
| Cannabis | Trajectory Group | 32.02 | <.001 |
| Nicotine | Trajectory Group | 12.33 | <.001 |
| **Peak Impulsivity across Trajectory Groups** | | | |
| **Subscale** | **Population** | ***F* statistic: Mean Peak** | **P value** |
| Mean | Trajectory Group | 13.35 | <.001 |
| Negative Urgency | Trajectory Group | 8.78 | <.001 |
| Perseverance | Trajectory Group | .61 | .60 |
| Positive Urgency | Trajectory Group | 10.04 | <.001 |
| Premeditation | Trajectory Group | 6.93 | <.001 |
| Sensation Seeking | Trajectory Group | 11.83 | <.001 |
| **Peak Inhibitory Control (Anti-Saccade) Performance across Trajectory Groups** | | | |
| **Trial Type** | **Population** | ***F* statistic: Mean Peak** | **P value** |
| Mean | Trajectory Group | 1.17 | .32 |
| Reward | Trajectory Group | 1.90 | .13 |
| Neutral | Trajectory Group | .59 | .62 |
| **Peak Tissue Iron (nT2*w) across Trajectory Groups** | | | |
| **Region** | **Population** | ***F* statistic: Mean Peak** | **P value** |
| Basal Ganglia | Trajectory Group | 2.31 | .075 |
| Caudate | Trajectory Group | 1.76 | .150 |
| NAcc | Trajectory Group | 2.72 | .044 |
| Pallidum | Trajectory Group | 3.30 | .020 |
| Putamen | Trajectory Group | 2.06 | .100 |

*Note. All metrics result from independent generalized additive mixed models (GAMMs) described in the main text. F and p values describe the significance an ANOVA test across trajectory groups, controlling for a smooth term for age and relevant sociodemographic variables.*

**Supplementary Table S12. Age by Trajectory Group Interaction Test Statistics on Impulsivity, Inhibitory Control, and Tissue Iron across Subscales, Trial Types, and Regions of Interest**

| **Impulsivity (UPPS)** | | | |
| --- | --- | --- | --- |
| **Subscale** | **Population** | ***F* statistic** | **P value** |
| Mean | Trajectory Group | 9.89 | <.001 |
| Negative Urgency | Trajectory Group | 4.70 | <.001 |
| Perseverance | Trajectory Group | 4.68 | <.001 |
| Positive Urgency | Trajectory Group | 2.677 | .015 |
| Premeditation | Trajectory Group | 4.68 | <.001 |
| Sensation Seeking | Trajectory Group | 6.969 | <.001 |
| **Inhibitory Control (Anti-Saccade)** | | | |
| **Trial Type** | **Population** | ***F* statistic** | **P value** |
| Mean | Trajectory Group | 1.13 | .316 |
| Reward | Trajectory Group | .802 | .502 |
| Neutral | Trajectory Group | 1.305 | .314 |
| **Tissue Iron (nT2*w)** | | | |
| **Region** | **Population** | ***F* statistic** | **P value** |
| Basal Ganglia | Trajectory Group | 2.60 | .0038 |
| Caudate | Trajectory Group | 2.18 | .020 |
| NAcc | Trajectory Group | 3.32 | .007 |
| Pallidum | Trajectory Group | .835 | .57 |
| Putamen | Trajectory Group | 1.885 | .0467 |

*Note. All metrics result from independent generalized additive mixed models (GAMMs) described in the main text. F and p values describe the significance an ANOVA test across trajectory groups, controlling for a smooth term for age and relevant sociodemographic variables.*

**Supplementary Table S13. Post-Hoc Comparisons for Age by Trajectory Group Interactions on Impulsivity, Inhibitory Control, and Tissue Iron across Subscales, Trial Types, and Regions of Interest**

| **Impulsivity (UPPS)** | | | | |
| --- | --- | --- | --- | --- |
| **Subscale** | **Population** | **Age 14 *p* value** | **Age 25 *p* value** | **Slope *p* value** |
| Mean | Low – Adult Increase | .327 | .312 | .076 |
| Mean | Low – Adolescent Increase | .196 | <.001 | <.001 |
| Mean | Low – Youth Peak | <.001 | <.001 | .648 |
| Mean | Adult Increase – Adolescent Increase | .033 | <.001 | .019 |
| Mean | Adult Increase – Youth Peak | <.001 | <.001 | .190 |
| Mean | Adolescent Increase – Youth Peak | .002 | .315 | <.001 |
| Negative Urgency | Low – Adult Increase | .303 | .662 | .698 |
| Negative Urgency | Low – Adolescent Increase | .955 | <.001 | <.001 |
| Negative Urgency | Low – Youth Peak | .148 | .026 | .288 |
| Negative Urgency | Adult Increase – Adolescent Increase | .332 | <.001 | <.001 |
| Negative Urgency | Adult Increase – Youth Peak | .021 | .007 | .492 |
| Negative Urgency | Adolescent Increase – Youth Peak | .227 | .018 | .011 |
| Perseverance | Low – Adult Increase | .828 | .753 | .702 |
| Perseverance | Low – Adolescent Increase | .366 | .002 | .025 |
| Perseverance | Low – Youth Peak | <.001 | .007 | .590 |
| Perseverance | Adult Increase – Adolescent Increase | .281 | .002 | .053 |
| Perseverance | Adult Increase – Youth Peak | <.001 | .013 | .357 |
| Perseverance | Adolescent Increase – Youth Peak | .012 | .287 | .006 |
| Positive Urgency | Low – Adult Increase | .155 | .550 | .562 |
| Positive Urgency | Low – Adolescent Increase | .598 | .027 | .096 |
| Positive Urgency | Low – Youth Peak | .003 | .033 | .631 |
| Positive Urgency | Adult Increase – Adolescent Increase | .071 | .005 | .279 |
| Positive Urgency | Adult Increase – Youth Peak | <.001 | .005 | .291 |
| Positive Urgency | Adolescent Increase – Youth Peak | .026 | .708 | .039 |
| Premeditation | Low – Adult Increase | .828 | .753 | .702 |
| Premeditation | Low – Adolescent Increase | .366 | .002 | .025 |
| Premeditation | Low – Youth Peak | <.001 | .007 | .590 |
| Premeditation | Adult Increase – Adolescent Increase | .281 | .002 | .053 |
| Premeditation | Adult Increase – Youth Peak | <.001 | .013 | .357 |
| Premeditation | Adolescent Increase – Youth Peak | .012 | .287 | .006 |
| Sensation Seeking | Low – Adult Increase | .441 | .028 | .063 |
| Sensation Seeking | Low – Adolescent Increase | .001 | .007 | .864 |
| Sensation Seeking | Low – Youth Peak | <.001 | <.001 | .827 |
| Sensation Seeking | Adult Increase – Adolescent Increase | .028 | .400 | .208 |
| Sensation Seeking | Adult Increase – Youth Peak | <.001 | .037 | .099 |
| Sensation Seeking | Adolescent Increase – Youth Peak | .122 | .411 | .994 |
| **Inhibitory Control (Anti-Saccade)** | | | | |
| **Trial Type** | **Population** | **Age 14 *p* value** | **Age 23 *p* value** | **Slope *p* value** |
| Mean | Low – Adult Increase | .841 | .484 | .616 |
| Mean | Low – Adolescent Increase | .260 | .631 | .649 |
| Mean | Low – Youth Peak | .087 | .276 | .682 |
| Mean | Adult Increase – Adolescent Increase | .270 | .934 | .430 |
| Mean | Adult Increase – Youth Peak | .098 | .694 | .414 |
| Mean | Adolescent Increase – Youth Peak | .552 | .884 | .910 |
| Reward | Low – Adult Increase | .687 | .794 | .763 |
| Reward | Low – Adolescent Increase | .522 | .898 | .593 |
| Reward | Low – Youth Peak | .076 | .833 | .364 |
| Reward | Adult Increase – Adolescent Increase | .368 | .751 | .470 |
| Reward | Adult Increase – Youth Peak | .058 | .968 | .269 |
| Reward | Adolescent Increase – Youth Peak | .295 | .776 | .839 |
| Neutral | Low – Adult Increase | .794 | .190 | .594 |
| Neutral | Low – Adolescent Increase | .563 | .488 | .984 |
| Neutral | Low – Youth Peak | .135 | .156 | .936 |
| Neutral | Adult Increase – Adolescent Increase | .775 | .964 | .689 |
| Neutral | Adult Increase – Youth Peak | .260 | .858 | .591 |
| Neutral | Adolescent Increase – Youth Peak | .396 | .880 | .967 |
| **Tissue Iron (nT2*w)** | | | | |
| **Region** | **Population** | **Age 14 *p* value** | **Age 25 *p* value** | **Slope *p* value** |
| Basal Ganglia | Low – Adult Increase | .982 | .671 | .865 |
| Basal Ganglia | Low – Adolescent Increase | .324 | .244 | .099 |
| Basal Ganglia | Low – Youth Peak | .016 | .051 | .263 |
| Basal Ganglia | Adult Increase – Adolescent Increase | .348 | .391 | .141 |
| Basal Ganglia | Adult Increase – Youth Peak | .027 | .108 | .210 |
| Basal Ganglia | Adolescent Increase – Youth Peak | .002 | .688 | .013 |
| Caudate | Low – Adult Increase | .544 | .990 | .537 |
| Caudate | Low – Adolescent Increase | .639 | .648 | .473 |
| Caudate | Low – Youth Peak | <.001 | .105 | .073 |
| Caudate | Adult Increase – Adolescent Increase | .318 | .627 | .216 |
| Caudate | Adult Increase – Youth Peak | .007 | .083 | .209 |
| Caudate | Adolescent Increase – Youth Peak | <.001 | .393 | .028 |
| NAcc | Low – Adult Increase | .376 | .051 | .124 |
| NAcc | Low – Adolescent Increase | .677 | .777 | .584 |
| NAcc | Low – Youth Peak | .021 | .830 | .006 |
| NAcc | Adult Increase – Adolescent Increase | .701 | .183 | .493 |
| NAcc | Adult Increase – Youth Peak | .002 | .073 | .203 |
| NAcc | Adolescent Increase – Youth Peak | .012 | .899 | .086 |
| Pallidum | Low – Adult Increase | .931 | .690 | .810 |
| Pallidum | Low – Adolescent Increase | .984 | .716 | .616 |
| Pallidum | Low – Youth Peak | .871 | .343 | .395 |
| Pallidum | Adult Increase – Adolescent Increase | .925 | .977 | .794 |
| Pallidum | Adult Increase – Youth Peak | .945 | .584 | .558 |
| Pallidum | Adolescent Increase – Youth Peak | .869 | .667 | .816 |
| Putamen | Low – Adult Increase | .078 | .823 | .518 |
| Putamen | Low – Adolescent Increase | .307 | .180 | .936 |
| Putamen | Low – Youth Peak | .165 | .362 | .133 |
| Putamen | Adult Increase – Adolescent Increase | .016 | .322 | .590 |
| Putamen | Adult Increase – Youth Peak | .003 | .318 | .393 |
| Putamen | Adolescent Increase – Youth Peak | .788 | .044 | .239 |

*Note. All metrics result from independent generalized additive mixed models (GAMMs) described in the main text investigating fixed values for ages that approximated the range of the data and the slope of the age effect. F and p values describe the significance an ANOVA test across trajectory group comparisons, controlling for relevant sociodemographic variables.*
