## Supplemental figures for "Developmental variation in dopamine neurobiology, neurocognitive functioning, and impulsivity shape substance use trajectories in youth"

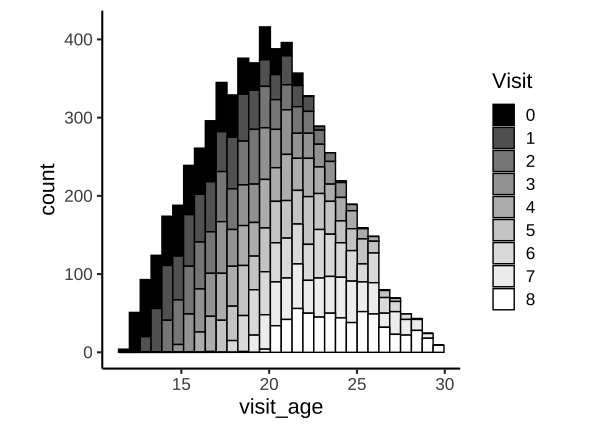

**Figure S1. Histogram of participant age and visit number.** Histogram of participant age and visit number. NCANDA-A used an accelerated longitudinal design with up to 9 visits per-participant (807 participants; 6164 sessions total following exclusions), spanning the full adolescent period and relevant transitional periods from adolescence to young adulthood (total age-range: 12-30 years old).
